# Parametric and AI-Assisted Design of Right-Handed Four-Helix Coiled Coils

**DOI:** 10.64898/2026.11.14.766675

**Authors:** Paul Kluge, Abibe Useini, Faezeh Shabani, Björn Kieslich, Aleksandr Kazimir, Norbert Sträter, Christina Lamers, Sibylle Gemming, Georg Künze

## Abstract

Right-handed, antiparallel α-helical coiled coils are rare in nature and remain challenging targets for de novo protein design. Here, we combined parametric backbone generation with deep-learning-based sequence design and structure prediction to systematically explore this underrepresented region of coiled-coil structural space. We compared three ProteinMPNN design strategies: repeat-constrained homotetramer design, unconstrained homotetramer design, and unconstrained heterotetramer design. Removing strict sequence repeat constraints increased the fraction of designs satisfying structural confidence and topology criteria approximately sixfold, from 0.44% to 2.80%, while simultaneous optimization of two distinct peptide sequences yielded a success rate of 3.04%. Selected designs were evaluated for assembly using a split-luciferase complementation assay in mammalian cells, followed by circular dichroism spectroscopy and molecular dynamics simulations. Five of six experimentally characterized designs exhibited predominantly α-helical structures, with apparent melting temperatures ranging from 55.2 to 72.5 °C. The crystal structure of the most stable design, Heteromer-2, determined at 1.92 Å resolution, confirmed the intended right-handed, antiparallel architecture, with a backbone RMSD of only 0.45 Å relative to the computational model. Our findings establish a strategy for designing noncanonical coiled-coil architectures by combining precise parametric control of backbone geometry with flexible, AI-assisted sequence optimization.

## Introduction

De novo protein design aims to create proteins with predefined structures and properties, extending the accessible structural repertoire beyond that sampled by natural evolution.^[1]^ Among the various protein architectures, α-helical coiled coils have long served as particularly tractable models for investigating sequence-structure relationships and developing protein design strategies. Coiled coils consist of two or more amphipathic α-helices that associate through predominantly hydrophobic interactions, typically involving knobs-into-holes side-chain packing, and wrap around a common superhelical axis.^[2]^ Their architectures can be described by a relatively small number of geometric parameters, including the number and relative orientation of helices, superhelical pitch and radius, and helix-helix packing geometry. Moreover, well-established relationships between sequence periodicity, side-chain interactions, and coiled-coil geometry provide a basis for rational design.^[3]^ This detailed understanding enables the construction of coiled-coil architectures with prescribed geometries rather than relying exclusively on naturally occurring structures as design templates.^[4]^ The structural principles and sequence determinants governing coiled-coil formation have been reviewed extensively elsewhere.^[3]^

Analysis of experimentally determined coiled-coil structures, such as those collected in the CC+ database, using the Coiled-Coil Crick Parameterization (CCCP) framework has revealed that naturally occurring coiled coils occupy a restricted region of the geometrically accessible parameter space.^[4, 5]^ Parametric backbone generation combined with computational sequence design has demonstrated that this repertoire can be expanded to include alternative geometries, yielding de novo helical bundles with high thermodynamic stability and close agreement between designed and experimentally determined structures.^[6]^ However, generating a geometrically feasible backbone represents only one part of the design challenge; identifying an amino acid sequence that reliably adopts the intended structure remains equally important. For canonical coiled coils, the underlying sequence-structure relationships are particularly well characterized. Their sequences typically follow a seven-residue heptad repeat, designated ***a–g***, in which predominantly hydrophobic residues at positions ***a*** and ***d*** form the buried core, while residues at the flanking ***e*** and ***g*** positions contribute to interhelical packing and electrostatic interactions. The identity and packing preferences of residues at the ***a*** and ***d*** positions strongly influence oligomerization state, as demonstrated by mutations in the GCN4 leucine zipper that switch its assembly between dimeric, trimeric, and tetrameric coiled coils.^[7]^ Electrostatic interactions involving the ***e*** and ***g*** positions, together with other interfacial interactions, further contribute to assembly stability, specificity, and helix orientation.^[8]^ Together, these findings established a robust knowledge-based framework for rational coiled-coil sequence design and have enabled the development of coiled-coil assemblies for applications in synthetic biology and biosensing.^[9]^

More recently, knowledge-based coiled-coil design has increasingly been complemented by computational approaches based on molecular energy functions and deep learning.^[9]^ A landmark study by Huang et al. demonstrated that systematic variation of backbone geometries generated using the Crick equations, followed by Rosetta-based sequence optimization, enables the design of hyperstable helical bundles with distinct superhelical geometries and topologies.^[6]^ Subsequently, Boyken et al. extended computational helical assembly design by introducing modular hydrogen-bond networks into protein-protein interfaces, enabling the generation of homo-oligomeric assemblies with precisely defined structures and interaction specificities.^[10]^ More recently, deep-learning-based sequence design methods such as ProteinMPNN have substantially improved the efficiency of identifying sequences compatible with prescribed protein backbones.^[11]^ These methods are frequently integrated with structure prediction tools, including AlphaFold2 and AlphaFold3, to evaluate the structural plausibility of designed sequences and prioritize candidates for experimental characterization.^[12, 13]^ Deep learning has also enabled generative backbone design, notably through diffusion-based approaches such as RFdiffusion, which can generate novel protein structures and assemblies without predefined structural templates.^[14, 15]^ Nevertheless, parametric backbone generation remains advantageous when precise and systematic control over geometric features is required. Combining such geometric control with computational sequence optimization offers a promising strategy for accessing coiled-coil architectures that are difficult to obtain using conventional sequence-based design rules. This motivates the present study, in which we focus on two defining structural features of helical bundles: superhelical handedness, described by the superhelical twist parameter ω₀, and the relative parallel or antiparallel orientation of the constituent helices.

The handedness of a coiled coil is closely related to the periodicity of its underlying sequence repeat. An ideal, undistorted α-helix contains approximately 3.63 residues per turn, whereas the canonical heptad repeat corresponds to seven residues distributed over two helical turns, equivalent to 3.5 residues per turn. To maintain the alignment of hydrophobic core residues along the helix-helix interface, this mismatch is accommodated by a left-handed superhelical twist. Consequently, canonical heptad-based coiled coils predominantly adopt left-handed geometries, characterized by negative values of ω₀. Right-handed supercoiling, by contrast, can arise when the periodicity of the hydrophobic repeat exceeds that of an undistorted α-helix. Noncanonical sequence repeats, including the eleven-residue hendecad and fifteen-residue pentadecad, provide alternative packing arrangements with average periodicities of 11/3 (approximately 3.67) and 15/4 (3.75) residues per turn, respectively. Hendecad repeats generally favor nearly straight or weakly right-handed assemblies, whereas pentadecad repeats can support more pronounced right-handed supercoiling.^[16, 17]^ Thus, systematic variation of sequence periodicity provides a means of modulating superhelical geometry from left-handed through approximately untwisted to right-handed arrangements.

The design of right-handed coiled coils therefore requires departures from the sequence periodicity and packing geometry characteristic of canonical heptad-based assemblies. Consistent with these additional constraints, right-handed coiled coils are considerably less common than their left-handed counterparts in structurally characterized proteins and occupy a sparsely populated region of coiled-coil parameter space.^[4]^ Nevertheless, naturally occurring examples demonstrate that right-handed coiled coils are structurally feasible, including the highly thermostable tetrameric coiled coil of the *Staphylothermus marinus* surface-layer protein tetrabrachion.^[18]^ Early computational design studies established that right-handed coiled coils can also be generated de novo. Harbury et al. designed a family of right-handed helical bundles by explicitly optimizing backbone geometry and side-chain packing, rather than restricting the design to a fixed natural backbone.^[19]^ Subsequent work demonstrated that stable right-handed tetrameric assemblies could be constructed entirely from genetically encoded amino acids.^[20]^ Huang et al. further showed that parametric backbone generation combined with Rosetta-based sequence design enables the construction of highly stable right-handed and untwisted helical bundles with experimentally validated structures.^[6]^ Despite these advances, right-handed coiled coils remain comparatively underexplored as de novo design scaffolds, particularly in combination with antiparallel helix arrangements.

The relative orientation of helices represents a second important determinant of coiled-coil architecture. In parallel assemblies, the constituent helices are aligned in the same N-to-C-terminal direction, whereas antiparallel assemblies contain helices oriented in opposite directions, resulting in distinct interhelical contacts and packing arrangements. Helix orientation is influenced by the distribution of hydrophobic and polar residues, electrostatic complementarity, and the relative register of interacting helices. In conventional heptad-based designs, favorable interactions between residues at the ***e*** and ***g*** positions can stabilize parallel arrangements, whereas alternative electrostatic patterns can promote antiparallel assembly.^[8]^ Antiparallel specificity can also be achieved by strategically positioning buried polar residues to favor one orientation over competing configurations.^[21]^ More recently, systematic sequence-based design strategies have enabled the construction of antiparallel four-helix bundles as homo- and heterotetrameric assemblies and as single-chain proteins.^[22]^ Nevertheless, controlling helix orientation remains challenging because the energetic differences between alternative arrangements can be small and depend on subtle changes in interfacial packing. Moreover, predicting the correct topology is not always straightforward, even with modern deep-learning tools. For example, Chubb et al. demonstrated that AlphaFold-Multimer can generate high-confidence coiled-coil models with incorrect helix orientations, highlighting the importance of extensive structural sampling and experimental validation.^[23]^ Thus, the reliable design of antiparallel assemblies remains an important challenge for both sequence design and structure prediction.

Superhelical handedness and helix orientation are governed by distinct but coupled structural and sequence determinants. Whereas superhelical twist is strongly influenced by sequence periodicity and the resulting hydrophobic packing pattern, the relative orientation of helices depends on the compatibility of interfacial interactions and packing arrangements. Simultaneously satisfying both requirements can therefore impose additional constraints on the sequences capable of stabilizing a given architecture. In particular, four-helix bundles combining right-handed supercoiling with antiparallel helix orientation remain poorly represented among experimentally characterized coiled coils. An important precedent was established by Huang et al., who reported a de novo designed, single-chain, antiparallel right-handed four-helix bundle with exceptional thermodynamic stability and a crystal structure closely matching the computational model.^[6]^ However, this structure represented one design within a broader investigation of distinct helical bundle geometries. Whether antiparallel right-handed four-helix bundles can be obtained systematically across different sequence architectures and design strategies remains insufficiently explored. Consequently, the combined design space of right-handed supercoiling and antiparallel helix orientation represents an attractive target for further investigation.

In this study, we systematically explored the design of right-handed four-helix bundles using a workflow that combines parametric backbone generation with deep-learning-based sequence design and structure prediction. Starting from Crick-parameterized backbones with positive superhelical twist values (ω₀ > 0), we explored geometries compatible with hendecad and pentadecad sequence periodicities. We compared design strategies imposing different degrees of sequence and architectural constraints, ranging from designs with prescribed sequence repeats to unconstrained sequence optimization of single-chain bundles and two-component A₂B₂ heterotetrameric assemblies. Candidate sequences were generated and evaluated using a deep-learning design and prediction pipeline, followed by experimental screening and structural characterization. Assembly of the designed peptides was initially assessed using a split-luciferase complementation assay, while their α-helical structure and thermal stability were characterized by circular dichroism spectroscopy. X-ray crystallographic analysis of a particularly stable design confirmed the intended right-handed, antiparallel four-helix architecture, with sub-ångström structural agreement between the experimentally determined structure and the computational design model.

Together, our results demonstrate that right-handed, antiparallel four-helix bundles can be accessed through a systematic combination of parametric backbone generation and AI- assisted sequence design. The developed workflow provides a framework for exploring noncanonical coiled-coil geometries and expands the repertoire of structurally characterized de novo helical bundle architectures.

## Results

### A deep-learning pipeline to explore the design space of right-handed helical bundles

To design coiled-coil architectures, we developed a computational workflow combining parametric backbone generation with deep-learning-based sequence design and structure prediction (**Figure 1A**). Four-helix bundle backbones were generated using the BundleGridSampler mover in Rosetta, which constructs helical assemblies based on Crick’s parameters.^[24]^ Backbone geometries were systematically varied by sampling the superhelical angular frequency (ω₀), superhelical radius (R₀), and rotational phase of the individual helices (Δω₁) (**Figure 1B**). The parameter ranges were informed by the analysis of naturally occurring coiled coils using the CCCP program^[4]^ and structural data from the SamCC-Turbo database^[25]^ (**Figure S1**). For antiparallel four-helix bundles, R₀ was sampled from 5 to 9 Å, ω₀ from 0 to +3.5 °/res, and Δω₁ from 0° to 350°. Throughout this study, positive values of ω₀ denote right-handed supercoiling, negative values denote left-handed supercoiling, and ω₀ = 0 corresponds to a straight, untwisted bundle.

**Figure 1.**
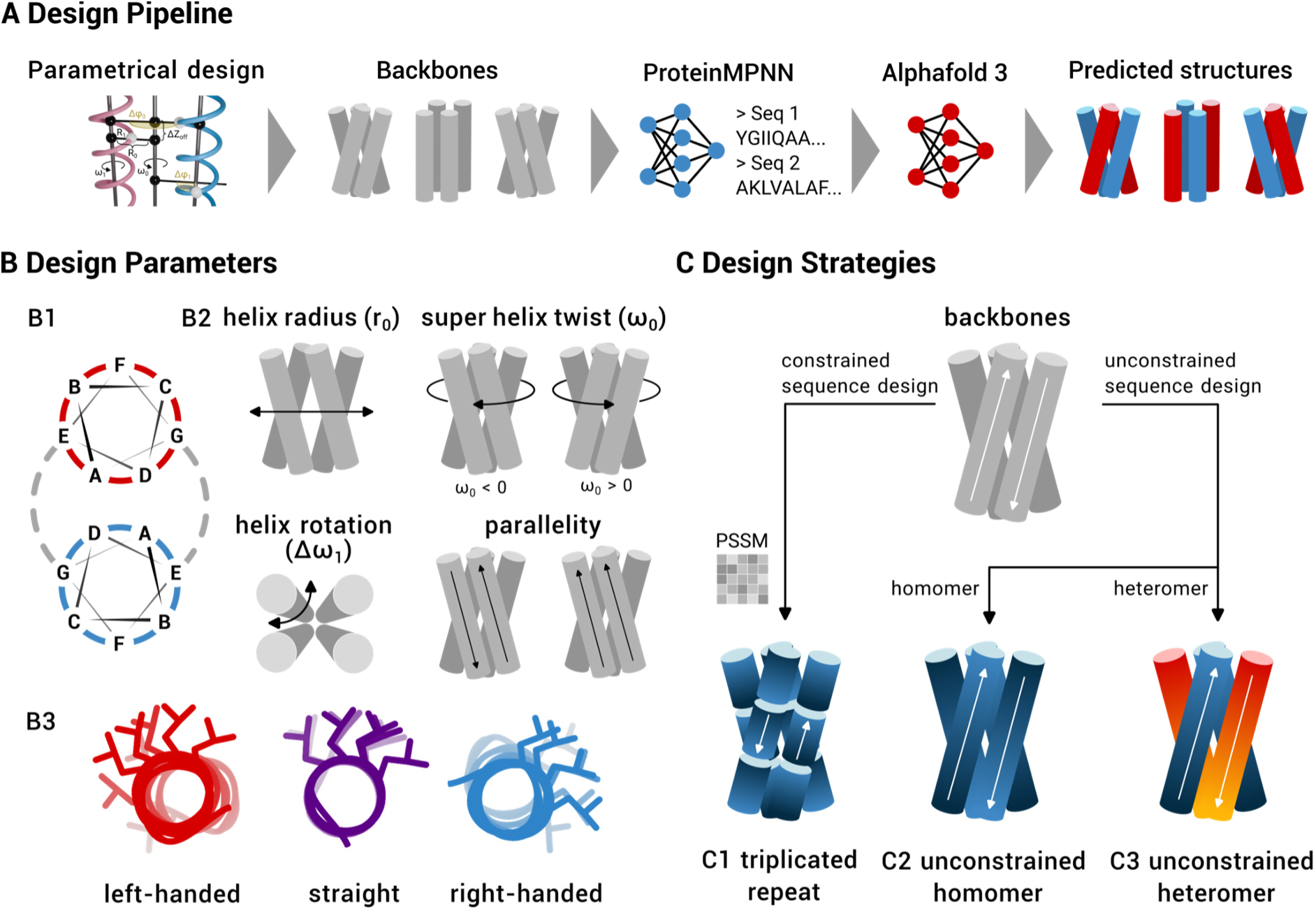
Computational design workflow, sampled backbone geometries, and sequence design strategies for right-handed four-helix bundles. (A) Overview of the computational design workflow comprising five steps: (1) definition and sampling of coiled-coil geometries based on Crick’s parameters, (2) generation of polyalanine four-helix bundle backbones, (3) sequence design using ProteinMPNN, (4) structure prediction of the designed sequences using AlphaFold3, and (5) selection of candidates based on structural confidence metrics and agreement with the intended bundle topology. (B) Geometric parameters defining the sampled coiled-coil backbones. (B1) Helical-wheel representation illustrating the sequence register and arrangement of core and interfacial positions. (B2) Schematic representation of the principal parameters varied during backbone generation: superhelical angular frequency (ω_0_), superhelical radius (R_0_), and rotational phase of the individual helices (Δω_1_). (B3) Illustration of the three superhelical handedness regimes: left-handed (ω_0_ < 0), untwisted (ω_0_ = 0), and right-handed (ω_0_ > 0). (C) Comparison of the three ProteinMPNN sequence design strategies. (C1) Repeat-constrained design, in which a single hendecad or pentadecad repeat unit is optimized and repeated along the peptide sequence. (C2) Unconstrained homotetramer design, in which the full-length peptide sequence is optimized while enforcing sequence identity across all four helices. (C3) Unconstrained heterotetramer design, in which two distinct peptide sequences are optimized simultaneously, with sequence identity enforced only between the two copies of each peptide.

Three sequence design strategies were investigated, differing in the extent of sequence periodicity and symmetry constraints imposed during ProteinMPNN optimization (**Figure 1C**). In **strategy 1 (repeat-constrained homotetramer design)**, a single hendecad or pentadecad sequence repeat was optimized and subsequently repeated to generate the full-length peptide, enforcing the intended sequence periodicity. In **strategy 2 (unconstrained homotetramer design)**, the complete peptide sequence was optimized without imposing an internal repeat, while maintaining sequence symmetry between the four helices to generate an A_4_ homotetramer. In **strategy 3 (unconstrained heterotetramer design)**, two distinct peptide sequences were optimized simultaneously to form an A₂B₂ heterotetramer. All three strategies were integrated into a common design and structure prediction workflow to assess their ability to generate sequences predicted to adopt the targeted right-handed coiled-coil geometries.

### Repeat-constrained sequence design yields few high-confidence helical bundles

To generate coiled coils with sequence periodicities characteristic of right-handed superhelical geometries, we initially employed a repeat-constrained design strategy based on hendecad and pentadecad repeats. In this approach, a single repeat unit was optimized and repeated two or three times to generate the full-length peptide sequence. Three variants of this strategy were evaluated. In the first approach, an 11-residue backbone was used for sequence design, and the resulting sequence was triplicated to generate a 33-residue peptide. In the second approach, a complete 33-residue backbone was generated, but only the central 11 residues were allowed to vary during ProteinMPNN optimization, while the two flanking repeats were retained as alanine residues. The optimized central repeat was subsequently triplicated before structure prediction with AlphaFold3 (AF3). In the third approach, sequence optimization was additionally guided by a position-specific scoring matrix (PSSM) derived from the amino acid frequencies of naturally occurring coiled coils with hendecad and pentadecad periodicities in the SamCC-Turbo database.^[25]^ This PSSM was used to bias ProteinMPNN toward amino acid preferences observed at equivalent repeat positions in natural coiled-coil structures (see **Table S1** and **S2**).

All three approaches yielded relatively few designs satisfying the computational selection criteria. Of 900 generated sequences, only four (0.44%) passed the stringent confidence and topology filter, requiring a mean pLDDT ≥ 90, an ipTM score ≥ 0.9, and the intended antiparallel four-helix bundle arrangement (**Table 1**). Although a somewhat larger proportion of sequences was predicted to form helical bundles, most failed to satisfy the combined structural confidence and topology criteria. These results indicate that imposing a strict sequence repeat strongly restricted the number of sequences predicted to adopt the targeted bundle architecture with high confidence.

**Table 1.** Success rates per design strategy. Fraction of designs passing the confidence filter (mean pLDDT ≥ 90 and ipTM ≥ 0.9, antiparallel bundle) for every strategy.

| **strategy** | **ω<sub>0</sub>** | **R<sub>0</sub>** | **Δω<sub>1</sub>** | **No. backbones generated** | **No. sequences generated** | **No. successful designs** | **Success rate (%)** |
|---|---|---|---|---|---|---|---|
| strict repeats | 0-1 °/res | 6-8 Å | 0° | 9 | 900 | 4 | 0.44 |
| unconstrained homomer | 0-3.5 °/res | 5-9 Å | 0° | 40 | 4000 | 112 | 2.80 |
| unconstrained heteromer | 0-3.5 °/res | 5-9 Å | 0-350° | 1656 | 33120 | 1010 | 3.04 |

We next investigated whether the ProteinMPNN sequence score could be used to prioritize designs before computationally more expensive structure prediction. However, no significant association was observed between the ProteinMPNN score and successful bundle formation (**Figure S2**). The point-biserial correlation coefficients were 0.032 (*p* = 0.51) for the first 11-residue repeat design set, 0.082 (*p* = 0.34) for the wrapped-repeat design set, and 0.027 (*p* = 0.76) for the PSSM-biased wrapped-repeat design set. Thus, within the investigated sequence libraries, ProteinMPNN scores alone did not reliably discriminate between sequences predicted to form the desired bundle architecture and those that failed the structural selection criteria. Consequently, preselection based on these scores would not provide an effective means of enriching the design libraries for successful candidates.

### Unconstrained sequence design increases the yield of high-confidence coiled coils

To investigate whether the imposed sequence periodicity limited design success, we next removed the repeat constraints and PSSM bias and optimized the entire 30-residue peptide sequence. Sequence identity between the four helices was maintained to enforce homotetrameric assembly, and the restricted amino acid alphabet excluding cysteine, proline, and aromatic residues was retained. Removing the internal sequence constraints substantially increased the fraction of designs satisfying the computational selection criteria. Of 4,000 generated sequences, 112 (2.80%) passed the confidence and topology filter, compared with only 0.44% for the repeat-constrained strategy (**Table 1**). Thus, allowing greater sequence flexibility increased the in silico design success rate by approximately sixfold, although the overall fraction of high-confidence predictions remained relatively low.

Further exploratory calculations revealed a pronounced dependence of design success on the rotational phase of the individual helices (Δω₁), which had been fixed at 0° in the preceding repeat-constrained design calculations. Sampling seven phase values distributed across the helical register resulted in substantial differences in the number of sequences satisfying the confidence filter, ranging from 32 to 143 designs per phase value. The highest numbers of successful predictions were obtained at phase angles near 296–347° and 41°, indicating that the rotational orientation of the helices relative to the bundle axis strongly influences the compatibility of designed sequences with the target structure. Notably, the previously selected phase angle of 0° was located close to this favorable region. These results identify Δω₁ as an important geometric parameter affecting the computational design success rate.

To increase confidence in the selected designs, we introduced two additional computational validation steps. First, sequences that satisfied the AF3 selection criteria were independently evaluated using Boltz-2.^[26]^ Designs were retained only if both prediction methods supported the intended coiled-coil architecture. This consensus-based approach was intended to reduce the likelihood of selecting designs whose predicted structures resulted from method-specific errors, although agreement between the two predictors does not establish experimental foldability. Second, we assessed sequence novelty by comparing the designed sequences against naturally occurring coiled coils represented in the CC+ database^[27]^ and the hendecad and pentadecad subsets of SamCC-Turbo.^[25]^ Among 24,790 natural sequences retrieved from these databases, no exact matches to the examined designed sequence segments were identified, and only two designs exhibited matches differing by three residues. These findings indicate that the generated sequences are distinct from the natural coiled-coil sequences represented in the reference datasets, supporting their sequence novelty within the scope of the database search.

### One-shot heterotetramer design outperforms iterative sequence optimization

Having investigated sequence design strategies for homotetrameric coiled coils, we next explored the design of heterotetrameric assemblies composed of two distinct peptide sequences (A₂B₂). We initially adopted an iterative design strategy in which the sequences of one pair of helices were optimized while those of the complementary pair were held fixed. The newly designed sequences were then threaded onto the corresponding backbone positions, followed by structure prediction and another round of optimization targeting the other helix pair. Repeated application of this procedure increased the proportion of designs satisfying the numerical structural confidence thresholds from 1% after the first round to 5% after the third round. However, this improvement was accompanied by a progressive deviation from the intended bundle geometry. From the second design round onward, the predicted assemblies predominantly adopted left-handed rather than the targeted right-handed superhelical geometry. Following the third round, most predicted assemblies also exhibited an arrangement of the two chain types that differed from the intended helix-pairing topology.

To overcome these limitations, we investigated a one-shot design strategy in which both peptide sequences were optimized simultaneously, allowing the interactions between the two chain types to be considered during a single ProteinMPNN calculation. This approach improved the recovery of designs with the intended topology. Of 33,120 generated heterotetrameric designs, 1010 (3.04%) passed the combined confidence and topology filter (**Table 1**). In contrast to the iterative approach, these designs retained the targeted right-handed superhelical geometry and antiparallel arrangement of the four helices. Thus, simultaneous optimization of both peptide sequences proved more effective than sequential redesign for generating high-confidence A₂B₂ assemblies with the desired backbone geometry.

### Designed coiled coils exhibit assembly-dependent luciferase activity in mammalian cells

For experimental characterization, we selected 12 homomeric designs from strategy 1 (1-1 to 1-12), comprising four that passed the structural confidence filter and eight additional designs with the highest scores among those that failed. From strategy 2, we selected 15 homomeric designs (2-1 to 2-15), and from strategy 3, 17 heteromeric designs comprising 34 distinct peptide sequences (3-1 to 3-17). The sequences and identifiers of all selected designs are provided in **Table S3**.

To investigate whether the designed peptides assemble in a cellular environment, we employed a split-firefly luciferase (split-Luc) complementation assay in human HEK293T cells. For each design, two constructs carrying complementary fragments of firefly luciferase were coexpressed, comprising two A₂ constructs for the homomeric designs or two AB constructs for the heteromeric designs. Association of the designed peptides brings the luciferase fragments into proximity, allowing reconstitution of enzymatic activity. The resulting firefly luciferase signal was normalized to the activity of a cotransfected Renilla luciferase reporter to account for differences in transfection efficiency and expression. Preliminary control experiments with the previously characterized, left-handed coiled coil DHD15^[28]^ established that firefly/Renilla luminescence ratios of approximately 1 or higher were consistent with substantial coiled-coil-dependent reporter complementation (**Figure S3**). This value therefore served as an empirical reference for evaluating the designed coiled coils and guiding candidate selection.

Of the 12 constructs generated using strategy 1 (repeat-constrained homotetramer design), only two, 1-2 and 1-7, produced appreciable luciferase signals, with relative luminescence ratios of 0.33 and 0.16, respectively. The remaining ten constructs exhibited ratios below 0.03, and none reached the empirical reference value of approximately 1 (**Figure 2A**).

**Figure 2.**
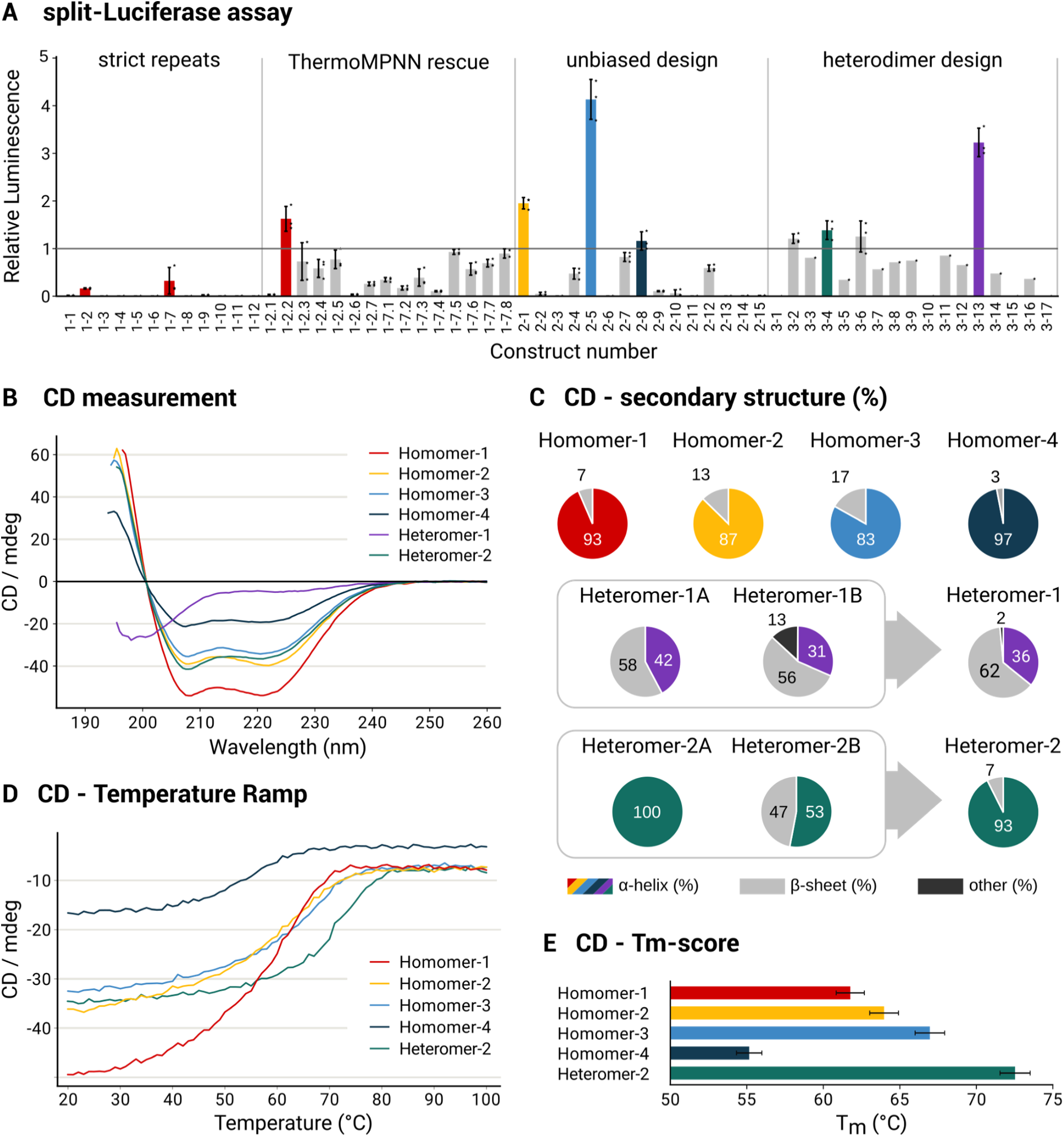
Cellular assembly and biophysical characterization of designed four-helix bundles. (A) Split-luciferase complementation assay showing firefly/Renilla luminescence ratios of the designed coiled coils (mean ± SD of three biological replicates). Designs selected for subsequent biophysical and structural characterization are highlighted in color. The horizontal line at 1 indicates the empirical reference value established using the DHD15 reference coiled coil (**Figure S3**). (B) Far-UV CD spectra of six selected helical bundles recorded at 20 °C. (C) Secondary-structure composition of the six bundles estimated from their CD spectra using BeStSel. (D) Thermal unfolding of Homomers 1–4 and Heteromer-2 monitored by temperature-dependent CD spectroscopy. (E) Melting temperatures (Tₘ) determined from the thermal unfolding measurements shown in (D).

To investigate whether the assembly propensity of the two initially active designs could be improved by stabilizing the intended coiled-coil structures, we performed computational sequence optimization using ThermoMPNN.^[29]^ For each parent design, ThermoMPNN was used to predict the effects of all possible single amino acid substitutions on folding free energy (ΔΔG). Substitutions predicted to improve stability were introduced into the parent sequences, followed by AF3 structure prediction to assess their compatibility with the intended bundle architecture. This procedure was repeated iteratively, using the most promising variants as starting points for subsequent optimization rounds. However, most substitutions did not improve the AF3 confidence metrics relative to the parent sequences, and many resulted in lower prediction confidence.

A total of 15 ThermoMPNN-derived variants were selected for experimental testing, comprising seven variants of design 1-2 (1-2.1 to 1-2.7) and eight variants of design 1-7 (1-7.1 to 1-7.8). The variants contained 1 or 2 amino acid substitutions relative to their respective parent sequences. Several variants exhibited substantially increased luciferase activity compared with their parent designs (**Figure 2A**). Two variants, 1-7.5 and 1-7.8, approached the luminescence level of the control, whereas variant 1-2.2 exceeded the control, reaching a relative luminescence of 1.63 ± 0.26, which represented an approximately tenfold increase over the parent design. These results demonstrate that ThermoMPNN-guided sequence optimization can improve the assembly-dependent reporter signal of selected designs, although the effects were strongly dependent on the parent sequence.

Among the 15 constructs generated using strategy 2 (unconstrained homotetramer design), three exhibited luciferase activities exceeding that of the control (**Figure 2A**). The strongest signal was observed for design 2-5, with a relative luminescence of 4.13 ± 0.42, followed by 2-1 (1.95 ± 0.12) and 2-8 (1.16 ± 0.19). In contrast, five constructs produced no detectable luciferase signal. Of the 17 heteromeric coiled-coil designs generated using strategy 3, four constructs (3-2, 3-4, 3-6, and 3-13) exhibited luciferase activities above the control level (**Figure 2A**). The strongest response was obtained for design 3-13, which reached a relative luminescence of 3.23 ± 0.30.

Collectively, the split-Luc complementation experiments revealed substantial differences in assembly-dependent reporter activity among the three design strategies. Unconstrained homotetramer and one-shot heterotetramer designs yielded the strongest luciferase signals, with several constructs exceeding the activity of the control. In contrast, repeat-constrained homotetramer designs exhibited generally weak reporter activity, which could be improved for selected variants through ThermoMPNN-guided sequence optimization. These findings indicate that relaxing sequence constraints facilitates the identification of coiled-coil designs capable of assembling in a cellular environment, while subsequent stability-oriented sequence optimization can further improve the performance of individual designs.

### Circular dichroism spectroscopy confirms α-helical structure and thermal stability of designed bundles

The four homotetrameric and two heterotetrameric designs with the highest split-luciferase signals were synthesized as peptides and characterized by circular dichroism (CD) spectroscopy. For clarity, the homotetrameric designs 1-2.2, 2-1, 2-5, and 2-8 are hereafter referred to as Homomer-1, Homomer-2, Homomer-3, and Homomer-4, respectively, while the heterotetrameric designs 3-4 and 3-13 are designated Heteromer-1 and Heteromer-2, respectively.

Far-UV CD spectra of all four homomeric designs exhibited the characteristic features of α-helical secondary structure, with estimated α-helical contents ranging from 83% to 97% (**Figure 2B,C**). Temperature-dependent CD measurements further demonstrated substantial thermal stability, with apparent melting temperatures (Tₘ) ranging from 55.2 ± 0.8 °C to 67.0 ± 1.0 °C (**Figure 2D,E**).

The two heteromeric designs exhibited different secondary-structure characteristics. For Heteromer-1, neither peptide was predominantly α-helical in isolation, and mixing the two peptides did not appreciably increase their α-helical content, which remained at 35.8% (**Figure 2B,C**). The low α-helical content precluded a reliable determination of thermal stability by CD spectroscopy. In contrast, the two constituent peptides of Heteromer-2 differed in their secondary-structure propensities, with one peptide displaying a predominantly α-helical structure in isolation and the other exhibiting only partial helicity. Mixing the two peptides increased the estimated α-helical content to 92.6% (**Figure 2B,C**), consistent with the formation of a predominantly α-helical assembly. Notably, Heteromer-2 exhibited an apparent Tₘ of 72.5 ± 1.0 °C, exceeding the values measured for all four homomeric designs (**Figure 2D,E**). This high thermal stability was consistent with the pronounced structural stability observed in the subsequent molecular dynamics (MD) simulations, which were performed for all six designs.

### Molecular dynamics simulations reveal differences in structural stability among designed coiled coils

To investigate the structural stability and conformational dynamics of the designed coiled coils, we performed three independent 1.05 µs MD simulations for each of the six coiled-coil systems at 30 °C and 40 °C. Structural stability was assessed by calculating the backbone root-mean-square deviation (RMSD) relative to the initial AF3 model and the trajectory-averaged structure (**Figures S4–S15, Tables S4** and **S5**).

The simulations revealed differences in conformational stability among the coiled-coil designs. Heteromer-2 exhibited the smallest deviations from the AF3 model, with replicate-averaged mean RMSDs of 0.98 Å at 30 °C and 1.17 Å at 40 °C (**Figures S9, S15**), followed by Heteromer-1 (1.36 and 1.61 Å; **Figures S8, S14**) and Homomer-3 (1.55 and 1.66 Å; **Figures S6, S12**). In contrast, Homomer-1 showed the largest structural deviations, with mean RMSDs of 2.54 Å at 30 °C and 3.03 Å at 40 °C, reaching maximum values of up to 4.55 Å (**Figures S4, S10**). Homomer-2 (2.06 and 2.13 Å; **Figures S5, S11**) and Homomer-4 (2.22 and 2.56 Å; **Figures S7, S13**) also exhibited larger deviations from their initial AF3 models. Increasing the simulation temperature from 30 °C to 40 °C generally increased the RMSD, particularly for Homomer-1 and Homomer-4. Representative RMSD trajectories and structural snapshots of Heteromer-2 and Homomer-4 at 30 °C illustrate these differences in conformational stability (**Figure 3**). RMSDs calculated relative to the trajectory-averaged structures were consistently lower, indicating that deviations from the initial AF3 models partly reflected conformational adjustments during the simulations. However, Homomer-1 also exhibited the largest RMSDs relative to its trajectory-averaged structures (1.76 and 1.97 Å at 30 °C and 40 °C, respectively), suggesting greater conformational flexibility.

**Figure 3.**
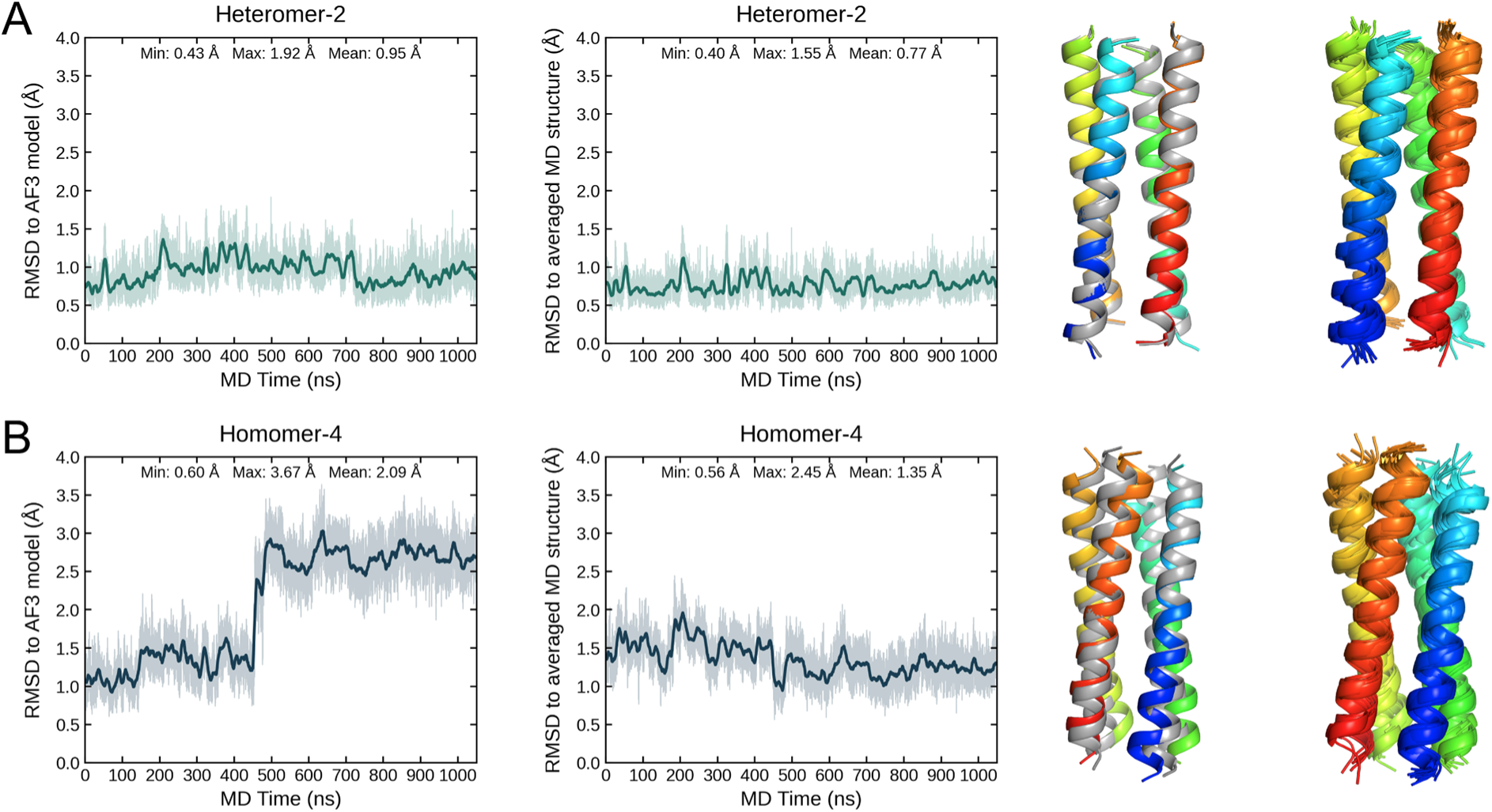
Representative molecular dynamics trajectories of Heteromer-2 and Homomer-4. Backbone RMSD over 1.05 μs of MD simulation at 30 °C for Heteromer-2 (top) and Homomer-4 (bottom). The left panels show the RMSD relative to the AF3 prediction, the right panels the RMSD relative to the trajectory-averaged MD structure. Far right: the simulated ensembles colored (rainbow from the N-to the C-terminus) and the predicted AF3 structure (grey).

These observations were broadly consistent with the experimentally determined thermal stabilities of the coiled coils. Heteromer-2 and Homomer-3, which exhibited low RMSDs during MD simulations, also showed the highest melting temperatures (Tₘ = 72.53 ± 0.98 °C and 66.96 ± 0.96 °C, respectively). In contrast, Homomer-1, Homomer-2, and Homomer-4, which showed larger deviations from their initial structures, exhibited lower melting temperatures of 61.76 ± 0.92 °C, 63.97 ± 0.94 °C, and 55.15 ± 0.83 °C, respectively. Although the RMSD values did not strictly correlate with the experimental melting temperatures, the simulations indicate greater structural rigidity of Heteromer-2 and Homomer-3 under the investigated conditions and reveal distinct differences in conformational dynamics among the designed coiled coils.

### Crystal structure of Heteromer-2 confirms the computational design model

To experimentally validate the designed coiled-coil architecture, we crystallized Heteromer-2, the most thermally stable of the six characterized designs, and determined its crystal structure at a resolution of 1.92 Å. The crystals belonged to space group P1 (triclinic; unit-cell parameters a = 25.1 Å, b = 85.4 Å, c = 91.9 Å, α = 63.3°, β = 89.5°, γ = 89.5°; **Table S6**) and contained six copies of the hetero-tetrameric coiled coil in the asymmetric unit. The experimentally determined structure closely matched the AF3 model (**Figure 4A**), with a Cα RMSD of 0.42 ± 0.02 Å and a backbone RMSD of 0.45 ± 0.01 Å over all 120 residues (mean ± SD across the six copies). This sub-ångström agreement confirms that Heteromer-2 adopts the designed right-handed, antiparallel four-helix architecture. The close structural agreement is also consistent with the MD simulations, in which Heteromer-2 exhibited the smallest deviations from its AF3 model among all six investigated coiled coils, with mean backbone RMSDs of 0.98 Å at 30 °C and 1.17 Å at 40 °C.

**Figure 4.**
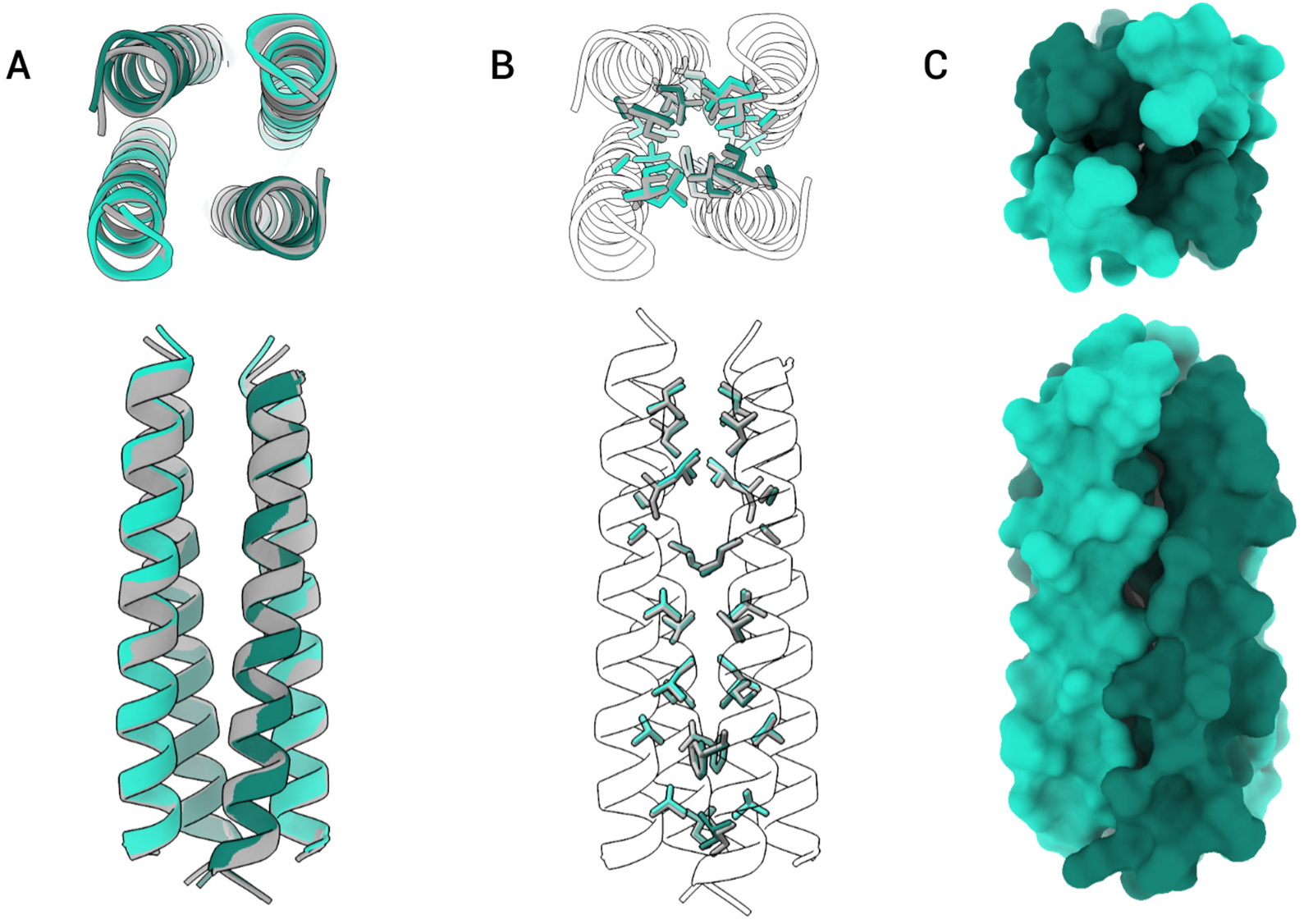
Crystal structure of Heteromer-2 and comparison with the AF3 design model. (A) Crystal structure of Heteromer-2 determined at 1.92 Å resolution, showing one of the six heterotetrameric assemblies in the asymmetric unit. The four helices are shown in cartoon representation, with the two distinct peptide chains colored separately. Superposition of the crystal structure (colored) with the AF3 design model (gray) demonstrates close agreement between the experimental and predicted structures. (B) Comparison of hydrophobic core packing in the crystal structure and AF3 model, with side chains shown in stick representation. (C) Surface representation of the Heteromer-2 crystal structure illustrating the overall coiled-coil architecture.

The crystal structure revealed a tightly packed hydrophobic core extending along nearly the entire length of the coiled coil (**Figure 4C**), consistent with its high α-helical content (92.6%) and elevated T_m_ of 72.5 ± 1.0 °C. Core packing was largely continuous, with only a small cavity at one interface layer where Ala23 (chain A) and Ala12 (chain B) face one another. The small side chains of these two alanine residues leave a local void between the helices.

The hydrophobic core packing was reproduced with remarkable accuracy in the AF3 model, with a side-chain RMSD of 0.76 ± 0.09 Å, which decreased to 0.42 ± 0.02 Å when Lys13, a residue at the core boundary that adopts a different snorkeling conformation, was excluded (**Figure 4B**). In contrast, larger deviations were observed for solvent-exposed side chains, with an RMSD of 1.97 ± 0.07 Å.

Together, these results demonstrate that the computational design accurately specifies not only the overall right-handed, antiparallel coiled-coil geometry but also the detailed side-chain packing of its hydrophobic core.

## Discussion

In this study, we demonstrate that combining parametric backbone generation with deep-learning-based sequence design provides an effective strategy for accessing right-handed, antiparallel four-helix coiled coils, an architecture that remains underrepresented among naturally occurring and de novo designed proteins. By comparing three sequence design strategies, we found that unconstrained sequence optimization increased the yield of high-confidence designs compared with approaches enforcing strict sequence periodicity. Experimental characterization identified several α-helical coiled coils with substantial thermal stability, including both homotetrameric and heterotetrameric assemblies. Most notably, the crystal structure of Heteromer-2 confirmed the intended right-handed, antiparallel architecture with sub-ångström agreement between the experimental structure and the computational model.

An important finding was that unconstrained sequence design performed substantially better than repeat-constrained design, despite the well-established relationship between sequence periodicity and coiled-coil geometry. Several factors may explain this observation. Enforcing identical hendecad or pentadecad repeats substantially reduces the accessible sequence space and prevents optimization of individual residues according to their local structural environments. Moreover, designing a repeat in the context of an alanine-containing backbone may introduce inconsistencies between the sequence environment encountered during optimization and that of the final assembly. Similarly, biasing amino acid selection toward the composition of natural coiled coils does not necessarily account for the specific packing requirements of a designed backbone. In contrast, unconstrained ProteinMPNN design allows the sequence to adapt to local differences in interhelical packing and residue interactions. Importantly, the desired superhelical geometry is already encoded in the parametrically generated backbone and therefore does not need to be imposed additionally through strict sequence repeats. For heterotetrameric coiled coils, simultaneous optimization of both peptide sequences was also more successful than iterative design, suggesting that jointly optimizing interactions between the two chain types is advantageous for maintaining the intended topology. Together, these findings support a design strategy in which the backbone defines the target geometry while sequence optimization is given sufficient flexibility to stabilize it.

The experimental and computational characterization methods provided complementary, although not always correlated, assessments of the designed coiled coils. The split-luciferase assay proved useful for identifying promising candidates, as five of the six selected designs exhibited predominantly α-helical structures by CD spectroscopy. However, luciferase activity did not directly correlate with α-helical content or thermal stability. This is expected because reporter complementation primarily reflects the proximity and productive association of the luciferase fragments and can additionally be influenced by expression levels and fusion construct geometry. It therefore does not directly report on the thermodynamic stability or oligomeric topology of the underlying coiled coil. Similarly, the MD simulations showed qualitative agreement with the experimentally determined thermal stabilities, with Heteromer-2 and Homomer-3 exhibiting both low backbone RMSDs and high melting temperatures. Nevertheless, this relationship was not universal. In particular, Heteromer-1 showed relatively small structural deviations during MD simulations despite its low α-helical content in solution. This discrepancy highlights an important limitation of MD simulations initiated from predicted structures: persistence of a folded conformation over the simulated timescale does not necessarily imply that the corresponding sequence will spontaneously assemble into that structure or that it represents the thermodynamically preferred state. Thus, the combined application of cellular screening, biophysical measurements, and MD simulations provides a more informative assessment of design success than any individual method.

The crystal structure of Heteromer-2 provides the strongest experimental validation of the computational design approach. Its backbone closely matches the AF3 model, with an RMSD of only 0.45 Å, while the hydrophobic core side-chain conformations are reproduced with similarly high accuracy. The structure confirms not only the intended right-handed superhelical geometry and antiparallel arrangement of the four helices but also the detailed packing interactions stabilizing the assembly. This degree of agreement is comparable to that reported for earlier parametrically designed helical bundles.^[6, 19]^ Importantly, Heteromer- 2 represents a structurally validated, self-assembling A_2_B_2_ heterotetramer, extending previous demonstrations of right-handed, antiparallel coiled-coil design to an assembly composed of two distinct peptide sequences. Although the precise topology of the other designed coiled coils remains to be established experimentally, the crystallographic result demonstrates that the combined parametric and AI-assisted workflow can accurately specify this noncanonical architecture at both the backbone and side-chain levels.

Together, our findings address the central objective of this study: exploring the designability of right-handed, antiparallel coiled coils beyond isolated examples. They demonstrate that geometrically defined backbones can be combined with flexible, deep-learning-based sequence optimization to generate stable coiled-coil assemblies with architectures that are uncommon in natural proteins. More broadly, this approach provides a framework for systematically expanding the accessible structural space of de novo coiled coils and developing new scaffolds with precisely defined geometries and assembly properties.

## Materials and Methods

### Parameter-space analysis of natural coiled-coil structures

Target ranges for the coiled-coil design parameters ω₀, R₀, and Δω₁ were derived from experimentally determined coiled-coil structures. The reference set comprised 171 coiled-coil assemblies, corresponding to 123 distinct PDB structures covering oligomeric states from dimer to nonamer (**Table S7**). Backbone coordinates of the reference structures were retrieved from the PDB and fitted to the Crick parameterization using the CCCP software (https://www.grigoryanlab.org/cccp/).^[4]^ The fitting procedure yielded the superhelical radius (R₀), superhelical angular frequency (ω₀), α-helical angular frequency (ω₁), and rotational phase of the α-helix about its local axis (Δω₁) for each structure. Parameter fitting was performed using cleaned coordinate files. The distributions of R₀, ω₀, ω₁, and Δω₁ obtained from the reference structures are shown in **Figure S1**.

### Backbone generation with Rosetta

Polyalanine backbones of four-helix coiled coils were generated using the BundleGridSampler mover implemented in Rosetta (version 2025.02+release.26484dedf0).^[30]^ BundleGridSampler constructs helical bundles directly from the Crick parameterization and systematically samples user-defined ranges of geometric parameters. Backbone bond lengths, bond angles, and dihedral angles were assigned by the mover, and the resulting structures were evaluated using the Rosetta ref2015 energy function with symmetry-aware scoring.^[31]^

Antiparallel four-helix bundle backbones were generated under C2 symmetry using two helix definitions. The parameter grid sampled the superhelical radius R₀ from 5 to 9 Å, the superhelical angular frequency ω₀ from 0 to 3.5°/res, the α-helical rotational phase Δω₁ from 41° to 347° in steps of 360°/7 ≈ 51°. The axial register offset Δt was fixed at 0.

Backbones of 28 residues per helix were used for the initial computational assessment of sequence design success across different backbone geometries. For the subsequent repeat-based sequence design experiments, helix lengths were selected to match the intended sequence periodicity. Hendecad-repeat designs comprised 33 residues per helix, corresponding to three 11-residue repeats, whereas pentadecad-repeat designs contained either 30 or 45 residues per helix, corresponding to two or three 15-residue repeats, respectively. For the consequent strategies 2 and 3, helices of length 30 were used.

### Sequence design with ProteinMPNN

Amino acid sequences were designed for the generated coiled-coil backbones using ProteinMPNN (commit 8907e667)^[11]^ with symmetry constraints applied as appropriate for each oligomeric architecture. Cysteine, proline, tryptophan, phenylalanine, and tyrosine were excluded from the design alphabet.

For each backbone, 20-100 sequences were generated using a sampling temperature of 0.2, different random seeds, and a batch size of one. Per-position amino acid probabilities were retained for subsequent analysis of sequence preferences. All sequences used for experimental characterization originated from the default ProteinMPNN model (v_48_020).

Three sequence design strategies were investigated, differing in the extent of sequence periodicity and interchain symmetry imposed during optimization.

In **strategy 1 (repeat-constrained homotetramer design)**, a single hendecad or pentadecad sequence repeat was optimized using ProteinMPNN. Corresponding positions in successive repeats were tied during sequence generation to enforce sequence periodicity along the helix. The resulting repeat unit was concatenated to generate the full-length sequence, comprising three hendecad repeats or two or three pentadecad repeats, depending on the target backbone length.

In **strategy 2 (unconstrained homotetramer design)**, the entire helix sequence was optimized without imposing an internal sequence repeat. Corresponding positions across all four helices were constrained to have identical amino acid identities, yielding a single sequence designed to assemble into an A_4_ homotetrameric coiled coil.

In **strategy 3 (heterotetramer design)**, two distinct peptide sequences were optimized to form an A_2_B_2_ heterotetramer, with the four helices arranged in an alternating ABAB pattern around the bundle axis. Both peptide sequences were optimized simultaneously, with corresponding positions tied within each pair of identical chains (A/A and B/B), but without sequence constraints between the A and B chains. This enabled the sequences and their mutual interfaces to be optimized in a single design step. The resulting designs were subsequently evaluated using AlphaFold3 structure prediction as described below.

### Sequence design with a position-specific scoring matrix bias

To bias ProteinMPNN sequence design toward amino acid compositions observed in naturally occurring right-handed coiled coils, a position-specific scoring matrix (PSSM) was constructed from the sequence patterns of structurally characterized coiled coils.

All coiled-coil entries with 11/3 and 15/4 periodicities were retrieved from the CCdb database generated using SamCC-Turbo^[25]^ (https://lbs.cent.uw.edu.pl/ccdb; database version: pdb_scan_20200717). Sequences were aligned according to the structural repeat register assigned by SamCC-Turbo rather than by primary sequence similarity, ensuring that structurally equivalent core and solvent-exposed positions were compared across different coiled coils. Within these alignments, hydrophobic core positions corresponding to hendecad (X..X…X…X) and pentadecad (X..X…X…X…X) repeat patterns were identified, where X represents a core position and the final X denotes the beginning of the subsequent repeat. Amino acid frequencies were calculated for each position within the respective repeat units and converted into a PSSM. Observed amino acid counts were converted directly into a probability distribution for each repeat position using a Boltzmann-weighted softmax, p(a|i) = exp(n(a,i)/T) divided by the sum of exp(n(b,i)/T) over all amino acids b, with a temperature T of 10.0. Positions with zero observed counts therefore retain a non-zero weight, because exp(0) = 1, which takes the role of a pseudocount. The PSSMs are given in **Tables S1** and **S2**.

The resulting PSSMs were incorporated into ProteinMPNN using its built-in PSSM bias functionality and applied to strategy 1 (repeat-constrained homotetramer design). The position-specific biases were repeated along the full-length sequence according to the corresponding hendecad or pentadecad periodicity. A PSSM softmax temperature of 10.0 and a PSSM mixing weight of 0.8 were used to combine the PSSM-derived amino acid preferences with the probabilities predicted by ProteinMPNN. The bias was applied as linear probability mixing (pssm_bias_flag = 1, pssm_multi = 0.8), so that the distribution sampled at each position was 0.2 times the ProteinMPNN probability plus 0.8 times the PSSM probability. For each backbone, sequences were generated both with and without the PSSM bias. The two design sets were compared in terms of amino acid composition at each repeat-register position and the proportion of sequences satisfying the structure prediction criteria.

### Structure prediction and computational selection of designs

Designed sequences were evaluated using AlphaFold3 (AF3) in the MSA-free mode.^[13]^ Designs were retained for further evaluation if the predicted structures satisfied several criteria: (i) a mean predicted local distance difference test (pLDDT) score greater than 90, (ii) an interface predicted template modeling (ipTM) score greater than 0.9, and (iii) the intended four-helix coiled-coil architecture, including right-handed superhelical geometry and antiparallel orientation of opposing helices. Visual inspection of the predicted structures was done using PyMOL (The PyMOL Molecular Graphics System, Schrödinger, LLC; version 3.1.6.1).

As an independent structure prediction approach, sequences that passed the AF3 screening criteria were also evaluated using Boltz-2 (release v2.2.1; commit cb04aecc; https://github.com/jwohlwend/boltz).^[26]^ Ten diffusion samples were generated for each sequence, using MSAs obtained from the integrated MSA server. Only designs for which both AF3 and Boltz-2 predicted the intended four-helix coiled-coil architecture and helix orientation were considered for experimental characterization.

To reduce sequence redundancy among the selected candidates, pairwise sequence similarities were assessed, and sequences were clustered using CD-HIT (version 4.8.1)^[32]^ and representatives of each cluster were chosen.

The sequence novelty of the designs was evaluated by comparison with naturally occurring coiled-coil sequences obtained from the CC+ database (http://coiledcoils.chm.bris.ac.uk/ccplus; downloaded Sep 1 2025)^[4, 5]^ and the 11/3 and 15/4 subsets of CCdb/SamCC-Turbo.^[25]^ Designed and natural sequences were compared using sliding windows matching the length of the designed repeat unit, i.e. 11 or 15 residues. Both exact and mismatch-tolerant matches were considered, allowing zero, two, or three amino acid mismatches per window. For each match, the reference database entry, sequence position, and percentage sequence identity over the aligned window were recorded. Across all designed sequences, the highest sequence identity to a natural coiled coil was 72.7% (8 of 11 residues), observed for two designs against entries of the CC+ database; no match was obtained against the 11/3 or 15/4 subsets of CCdb at any of the tested mismatch tolerances.

### Stability optimization with ThermoMPNN

The sequences of structure 1-2 and 1-7 were shown to weakly reconstitute split-Luc activity and were selected for further sequence optimization. Mutational effects on protein stability were predicted using ThermoMPNN^[29]^ (https://github.com/Kuhlman-Lab/ThermoMPNN; commit 11a1c5b), a graph neural network model trained to predict changes in folding free energy (ΔΔG) resulting from amino acid substitutions. The AF3-predicted structures of the parent designs were used as inputs for ThermoMPNN inference with the default model weights. For each design, predicted ΔΔG values were calculated for all possible single amino acid substitutions, yielding a comprehensive computational single-mutant stability landscape.

For each parent design, all substitutions with a predicted stabilizing effect were selected for further evaluation, yielding 220 variants for the first and 160 variants for the second parent design. ThermoMPNN was applied to all 33 sequence positions and all single amino acid substitutions, resulting in 726 and 759 scored substitutions for the two parent designs. Every substitution with a predicted ΔΔG below 0 kcal/mol was carried forward. Each selected substitution was introduced individually into the parent sequence, and the resulting single- point mutants were subjected to AF3 structure prediction using the settings described above. The predicted structures were compared with those of their respective parent designs in terms of mean pLDDT and ipTM. A second optimization round was subsequently performed using the most promising mutant from the first round as the new parent sequence. Two optimization rounds were performed in total for each initial design, and no further iterations were carried out. 22 variants from the first round served as parent sequences in the second round, which generated 2,794 additional variants. Where none of the tested substitutions improved the structural confidence metrics relative to the parent sequence, the original parent sequence was retained for further analysis.

### Molecular dynamics simulations

All-atom molecular dynamics (MD) simulations of the designed helical bundles were performed using AMBER 20 and the ff19SB force field.^[33, 34]^ The helical bundle structures predicted using AF3 were used as starting configurations.^[13]^ Residue pKa values were predicted and protonation states assigned using the H++ web server.^[35]^

Each helical bundle was solvated in a truncated octahedral box of optimal point charge (OPC) water molecules,^[36]^ with a minimum distance of 12 Å between any protein atom and the simulation box boundaries. Counterions were added to neutralize the net charge of the system, followed by additional NaCl to obtain a concentration of 150 mM. Bonds involving hydrogen atoms were constrained using the SHAKE algorithm.^[37]^ A cutoff of 10 Å was applied to short-range nonbonded interactions, and long-range electrostatic interactions were treated using the particle-mesh Ewald method.^[38]^

Each system was energy-minimized in three consecutive stages: 5,000 minimization steps applied to the solvent molecules and ions, 5,000 steps applied to the protein, and 10,000 steps applied to the entire system. Subsequently, the systems were heated from 0 K to either 303 K (30 °C) or 313 K (40 °C) over 500 ps in the NVT ensemble using a 1-fs time step, while the protein atoms were restrained to their energy-minimized coordinates. Temperature control during heating was achieved using an Andersen thermostat with a heat-bath coupling time constant of 1 ps.

Following heating, each system was equilibrated for 5 ns in the NPT ensemble at the respective simulation temperature and a reference pressure of 1 bar. During this stage, weak positional restraints with a force constant of 1 kcal mol⁻¹ Å⁻² were applied to the protein backbone atoms. Langevin dynamics with a collision frequency of 2 ps⁻¹ and a 2-fs integration time step were used. All positional restraints were subsequently removed, and the systems were equilibrated for an additional 5 ns.

Production simulations were performed for 1,050 ns under periodic boundary conditions in the NPT ensemble using Langevin dynamics at either 303 K or 313 K. Three independent replicates were simulated for each helical bundle at each temperature, resulting in a total simulation time of 6.3 µs per bundle system. Across the six investigated helical bundle systems, a cumulative simulation time of 37.8 µs was obtained.

### Split-luciferase constructs

Assembly of the designed coiled coils in mammalian cells was assessed using a split-firefly luciferase complementation assay, following the strategy described by Merljak et al.^[39]^ Each designed four-helix coiled coil was divided into two polypeptide chains, each comprising two helices. The optimal attachment sites for the luciferase fragments were determined in preliminary experiments using the previously characterized, left-handed coiled coil DHD15.^[28]^ Different configurations were tested by fusing the N-terminal (nLuc) and C-terminal (cLuc) fragments of firefly luciferase to either terminus of the two DHD15 chains (**Figure S3**). The highest complementation activity was obtained with nLuc fused to the N- terminus of one two-helix chain and cLuc to the C-terminus of the complementary chain. This configuration was used for all subsequent designs. Within each construct, consecutive helices and the respective luciferase fragment were connected by flexible (GGGGS)_2_ linkers. Association of the two polypeptide chains into the designed four-helix coiled coil brings the complementary luciferase fragments into proximity, enabling reconstitution of enzymatic activity. The amino acid sequences of all constructs are provided in **Table S3**.

### Gene synthesis and split-luciferase reporter plasmid preparation

Coding sequences for the designed coiled-coil chains were codon-optimized for expression in human cells and synthesized as eBlocks or gBlocks gene fragments by Integrated DNA Technologies (Coralville, IA, USA). Lyophilized gBlocks were resuspended in 50 µL nuclease-free water to a final concentration of 20 ng µL⁻¹, whereas eBlocks were supplied at a concentration of 10 ng µL⁻¹.

Two in-house generated pCDSec2 expression plasmids, derived from pcDNA3.4 (https://www.addgene.org/253816/), encoding either the nLuc or cLuc reporter fragment (**Table S8**) were linearized by restriction digestion with BamHI and KpnI (New England Biolabs) for the nLuc reporter plasmid or NotI and KpnI (New England Biolabs) for the cLuc reporter plasmid. Linearized vectors were separated by electrophoresis on a 1% (w/v) agarose gel in 1× TAE buffer containing SYBR Safe DNA stain and purified using the Monarch DNA Gel Extraction Kit (New England Biolabs), according to the manufacturer’s instructions. DNA concentrations were determined using a NanoDrop One spectrophotometer (Thermo Fisher Scientific).

For plasmid assembly, 100 ng of linearized vector and 30 ng of the corresponding synthetic gene fragment were combined using the NEBuilder HiFi DNA Assembly Master Mix (New England Biolabs), according to the manufacturer’s instructions.

The assembled plasmids were transformed into chemically competent *E. coli* NEB 5-alpha cells (New England Biolabs) by heat shock. Transformed cells were selected overnight on LB agar containing 100 µg mL⁻¹ ampicillin. Individual colonies were cultured in 4 mL LB medium supplemented with 100 µg mL⁻¹ ampicillin at 37 °C overnight with shaking. Plasmid DNA was isolated using the ZymoPURE Plasmid Miniprep Kit (Zymo Research, cat. no. D4210), according to the manufacturer’s instructions, and quantified using a NanoDrop spectrophotometer. All constructs were verified by Sanger sequencing (Azenta Life Sciences/GENEWIZ, Leipzig, Germany). Sequencing was performed with the forward primer 5′-CCAGCCTCCGGACTCTAGAG-3′, which anneals to the vector-derived region immediately upstream of the insert, and the reverse primer 5′-CCACATAGCGTAAAAGGAGCAAC-3′.

### Cell culture and plasmid transfection

HEK293T/17 cells (ATCC CRL-11268; obtained from LGC Standards GmbH, Wesel, Germany) were maintained in Dulbecco’s modified Eagle medium (DMEM, 1×, with GlutaMAX; Gibco, Thermo Fisher Scientific) supplemented with 10% fetal bovine serum (FBS; Gibco, Thermo Fisher Scientific, Waltham, MA, USA) at 37 °C in a humidified atmosphere containing 5% CO₂.

Before seeding, cells were detached from the culture flask by gentle tapping and resuspended by repeated pipetting. For cell counting, 12 µL of the cell suspension was mixed 1:1 with methylene blue, and cell numbers were determined using a Neubauer counting chamber. Cells were seeded into 24-well plates (Sarstedt, Nümbrecht, Germany) at a density of 3 × 10⁵ cells per well in 1 mL DMEM supplemented with FBS. A common cell suspension was prepared for all wells within each experiment to ensure consistent seeding densities.

Plasmid transfection was performed 24 h after cell seeding. For each well, 1 µg of each of the two split-luciferase reporter plasmids encoding the respective coiled-coil chains and 1 µg of a Renilla luciferase control plasmid (pRL-TK Vector; Promega, Madison, WI, USA) were combined in 40 µL Opti-MEM (Gibco, Thermo Fisher Scientific). Subsequently, 9 µL polyethylenimine (PEI MAX, MW 40,000; Polysciences, cat. no. 24765; supplied by VWR International) was added, and the mixture was incubated for 15 min at room temperature. The culture medium was replaced with 200 µL serum-free DMEM, and the transfection mixture was added dropwise to the cells. After 6–8 h, the transfection medium was replaced with 1 mL DMEM. Cells were harvested 48 h after transfection for luciferase activity measurements.

### Dual-luciferase reporter assay

Firefly and *Renilla* luciferase activities were measured using the Dual-Luciferase Reporter Assay System (Promega, Madison, WI, USA; cat. no. E1910), according to the manufacturer’s instructions. Firefly luciferase activity was normalized to the activity of the constitutively expressed, cotransfected Renilla luciferase control (1 µg plasmid per well; pRL-TK Vector, Promega, Madison, WI, USA) to account for differences in transfection efficiency.

At 48 h post-transfection, the culture medium was removed, and the cells were washed once with 1× phosphate-buffered saline (PBS). After complete removal of the wash buffer, cells were lysed by adding 100 µL of 1× Passive Lysis Buffer (PLB) per well and gently rocking the plate for 15 min at room temperature. Cell lysates were subsequently transferred to 1.5 mL microcentrifuge tubes.

For each measurement, 20 µL of cell lysate was mixed with 100 µL Luciferase Assay Reagent II (LAR II), and firefly luciferase activity was recorded. Subsequently, 100 µL Stop & Glo Reagent was added to the same sample to quench firefly luciferase activity and initiate the *Renilla* luciferase reaction. Luminescence was measured using a Synergy H1 multimode plate reader (BioTek Instruments, Winooski, VT, USA) without a measurement delay and with an integration time of 12 s. Reagents were pipetted manually, and data were acquired using Gen5 software version 3.12 (BioTek Instruments). Measurements were performed in triplicate.

Firefly luciferase activity was divided by the corresponding *Renilla* luciferase activity to obtain relative luminescence values. For the DHD15 controls, a ratio of ca. 4 was obtained with the optimized reporter configuration, whereas a configuration with both luciferase fragments fused N-terminally yielded a ratio of ca. 1 (**Figure S3**). The latter value was used as an empirical reference for interpreting assembly-dependent reporter activity and guiding the selection of designs for subsequent biophysical and structural characterization.

### Circular dichroism spectroscopy

Far-UV circular dichroism spectra were recorded on a Jasco J-1500 spectropolarimeter using a quartz cuvette with a path length of 10 mm. Lyophilized peptides (ordered from JPT Peptide Technologies GmbH, Berlin, Germany) were dissolved in TBS buffer and measured at a concentration of 0.2 mg mL⁻¹. Spectra were acquired from 260 to 185 nm with a data pitch of 0.5 nm in continuous scanning mode at a scanning speed of 100 nm/min, a digital integration time of 1 s and a spectral bandwidth of 1.0 nm, averaging five accumulations per spectrum. A buffer baseline was subtracted from each spectrum, and CD and high-tension voltage were recorded in parallel.

Thermal denaturation was followed at 222 nm between 20 and 100 °C using a temperature gradient of 1 °C min⁻¹ and a data interval of 1 °C, with the temperature gradient halted during each data point. In addition, full spectra were recorded at nine temperatures (20, 30, 40, 50, 60, 70, 80, 90 and 100 °C) using the same wavelength settings but three accumulations per spectrum. Melting temperatures were obtained from the 222 nm signal. Secondary-structure composition was estimated from the 20 °C spectra using BeStSel (https://bestsel.elte.hu/).

### Protein crystallization

For crystallization of Heteromer-2, the two peptides were dissolved in TBS buffer (20 mM Tris, 150 mM NaCl, pH 7.4) and mixed in equal amounts at a concentration of 8-10 mg mL^-1^.

Initial crystallization trials were performed using a high-throughput screening approach, testing approximately 400 conditions in a 96-well plate format. Crystallization drops were prepared using the sitting-drop vapor diffusion method, with 200 nL drops consisting of equal volumes of protein sample and reservoir solution (100 nL each), equilibrated against 100 μL reservoir solution. A Mosquito Xtal3 pipetting robot (SPT Labtech, Melbourn, England) was used to set up these drops. Conditions yielding initial crystals were subsequently selected and manually reproduced using the hanging-drop method, where 1 μL of protein solution was mixed with an equal volume of reservoir solution.

Heteromer-2 crystals formed under conditions containing 1 M LiCl, 30% w/v PEG 6000 and 0.1 M MES, pH 6 and grew within one week to a size of approximately 30–80 μm. Single crystals were flash-frozen in liquid nitrogen using the crystallization solution supplemented with 15% glycerol as a cryoprotectant.

### X-ray structure determination

Crystal diffraction data were collected using synchrotron radiation at beamline P14 of PETRA III (DESY Synchrotron, Hamburg, Germany). The diffraction data were indexed, integrated, and scaled using XDS (version 16 April 2026)^[40]^ and STARANISO (version 3.0.6beta) as implemented in ISPyB^[41]^ at DESY.

Structures were determined using molecular replacement (MR) with Phaser and MOLREP within CCP4i2 (CCP4 version 9.0),^[42]^ using the AF3 predicted structure as search model. Refinement was performed using jelly-body refinement in REFMAC (version 5.8.0431), and model building was completed in Coot (version 0.9.8.96).^[43]^ Final refinement was performed in Phenix (version 2.1-6048).^[44, 45]^

Data collection and refinement statistics are listed in **Table S6**.

### Funding

This work was supported by the Deutsche Forschungsgemeinschaft (DFG, German Research Foundation) through the Collaborative Research Centre TRR 386 HYP*MOL (project number 514664767; subproject B02 to G.K. and S.G. and subproject Z02 to C.L.).

G.K. acknowledges additional financial support from the German Federal Ministry of Education and Research (BMBF) and the Saxon State Ministry for Science, Culture and Tourism (SMWK) through the Center for Scalable Data Analytics and Artificial Intelligence Dresden/Leipzig (ScaDS.AI). Open-access publication was supported by the Open Access Publishing Fund of Leipzig University, funded by the DFG under the Open Access Publication Funding programme.

## Acknowledgements

The authors thank the high performance computing centers at Leipzig University and at the NHR center of TU Dresden for providing the computational resources. We further acknowledge DESY (Hamburg, Germany), a member of the Helmholtz Association HGF, and the EMBL for the provision of experimental facilities at synchrotron beamline P14. We thank the on-site scientists for their support during X-ray data collection.

