## Supporting Information for "Parametric and AI-Assisted Design of Right-Handed Four-Helix Coiled Coils"

### Supplementary Tables

**Table S1.** Position specific amino acid bias for the pentadecad-repeat. Sequences were retrieved from SamCC-Turbo database as described in Methods.

| | aa | | | | | | | | | | | | | | | | | | | |
|---|---|---|---|---|---|---|---|---|---|---|---|---|---|---|---|---|---|---|---|---|
| **position** | **A** | **C** | **D** | **E** | **F** | **G** | **H** | **I** | **K** | **L** | **M** | **N** | **P** | **Q** | **R** | **S** | **T** | **V** | **W** | **Y** |
| **1** | 2 | 0 | 1 | 0 | 0 | 4 | 0 | 1 | 0 | 0 | 3 | 0 | 0 | 1 | 1 | 0 | 2 | 1 | 0 | 0 |
| **2** | 3 | 0 | 0 | 2 | 0 | 1 | 0 | 0 | 6 | 1 | 0 | 1 | 0 | 0 | 0 | 2 | 0 | 0 | 0 | 0 |
| **3** | 0 | 0 | 0 | 2 | 1 | 0 | 0 | 1 | 4 | 1 | 0 | 3 | 0 | 2 | 0 | 0 | 2 | 0 | 0 | 0 |
| **4** | 0 | 0 | 0 | 0 | 0 | 0 | 0 | 5 | 0 | 3 | 0 | 0 | 0 | 0 | 0 | 0 | 0 | 1 | 0 | 0 |
| **5** | 0 | 0 | 2 | 2 | 0 | 0 | 0 | 2 | 2 | 0 | 3 | 2 | 0 | 0 | 0 | 1 | 0 | 2 | 0 | 0 |
| **6** | 4 | 0 | 1 | 4 | 0 | 0 | 0 | 0 | 0 | 0 | 0 | 2 | 0 | 1 | 2 | 2 | 0 | 0 | 0 | 0 |
| **7** | 1 | 0 | 0 | 7 | 0 | 0 | 0 | 1 | 2 | 0 | 0 | 0 | 0 | 2 | 0 | 0 | 0 | 0 | 0 | 3 |
| **8** | 7 | 0 | 0 | 0 | 1 | 0 | 0 | 1 | 0 | 0 | 1 | 0 | 0 | 0 | 0 | 2 | 2 | 1 | 0 | 1 |
| **9** | 1 | 0 | 2 | 4 | 0 | 0 | 0 | 0 | 0 | 0 | 0 | 4 | 0 | 2 | 2 | 0 | 0 | 1 | 0 | 0 |
| **10** | 0 | 0 | 1 | 3 | 0 | 0 | 0 | 2 | 1 | 2 | 2 | 0 | 0 | 3 | 0 | 0 | 0 | 2 | 0 | 0 |
| **11** | 2 | 0 | 0 | 5 | 0 | 0 | 0 | 0 | 4 | 1 | 0 | 0 | 0 | 0 | 0 | 0 | 0 | 0 | 0 | 2 |
| **12** | 1 | 0 | 0 | 1 | 2 | 0 | 0 | 0 | 1 | 0 | 1 | 1 | 0 | 0 | 0 | 0 | 2 | 6 | 0 | 0 |
| **13** | 0 | 0 | 2 | 3 | 0 | 1 | 0 | 0 | 1 | 2 | 0 | 2 | 0 | 0 | 1 | 2 | 3 | 0 | 0 | 0 |
| **14** | 0 | 0 | 0 | 1 | 0 | 0 | 2 | 0 | 1 | 0 | 0 | 2 | 0 | 3 | 4 | 0 | 0 | 0 | 0 | 0 |
| **15** | 2 | 0 | 0 | 3 | 0 | 0 | 0 | 2 | 2 | 2 | 0 | 0 | 0 | 1 | 0 | 2 | 0 | 0 | 0 | 2 |

**Table S2.** Position specific amino acid bias for the hendedecad-repeat. Sequences were retrieved from the SamCC-Turbo database as described in Methods.

| | aa | | | | | | | | | | | | | | | | | | | |
|---|---|---|---|---|---|---|---|---|---|---|---|---|---|---|---|---|---|---|---|---|
| **position** | **A** | **C** | **D** | **E** | **F** | **G** | **H** | **I** | **K** | **L** | **M** | **N** | **P** | **Q** | **R** | **S** | **T** | **V** | **W** | **Y** |
| **1** | 5 | 0 | 1 | 0 | 3 | 1 | 2 | 17 | 10 | 28 | 9 | 2 | 0 | 1 | 5 | 3 | 4 | 8 | 4 | 5 |
| **2** | 14 | 0 | 3 | 21 | 5 | 2 | 0 | 1 | 14 | 11 | 1 | 2 | 11 | 9 | 4 | 4 | 1 | 0 | 0 | 5 |
| **3** | 21 | 0 | 6 | 13 | 5 | 1 | 0 | 5 | 21 | 2 | 0 | 5 | 2 | 9 | 5 | 2 | 4 | 5 | 0 | 2 |
| **4** | 9 | 0 | 0 | 0 | 4 | 0 | 0 | 16 | 8 | 26 | 2 | 1 | 0 | 10 | 10 | 3 | 8 | 8 | 1 | 2 |
| **5** | 4 | 0 | 10 | 29 | 1 | 0 | 1 | 4 | 8 | 7 | 8 | 5 | 0 | 6 | 17 | 2 | 1 | 5 | 0 | 0 |
| **6** | 11 | 1 | 5 | 12 | 1 | 3 | 0 | 6 | 16 | 5 | 0 | 8 | 2 | 19 | 8 | 2 | 1 | 4 | 1 | 3 |
| **7** | 17 | 0 | 3 | 19 | 2 | 1 | 0 | 0 | 18 | 4 | 5 | 5 | 2 | 17 | 8 | 1 | 3 | 1 | 0 | 2 |
| **8** | 14 | 2 | 1 | 4 | 9 | 0 | 3 | 6 | 5 | 11 | 5 | 2 | 0 | 3 | 7 | 6 | 14 | 3 | 1 | 11 |
| **9** | 9 | 0 | 3 | 15 | 3 | 0 | 1 | 2 | 23 | 9 | 1 | 8 | 0 | 12 | 11 | 5 | 1 | 2 | 0 | 3 |
| **10** | 6 | 0 | 6 | 30 | 1 | 2 | 0 | 12 | 12 | 7 | 0 | 2 | 2 | 11 | 11 | 3 | 1 | 2 | 0 | 0 |
| **11** | 7 | 0 | 9 | 26 | 1 | 0 | 3 | 3 | 14 | 10 | 0 | 3 | 0 | 9 | 13 | 1 | 2 | 4 | 1 | 2 |

**Table S3.** Designed coiled-coil constructs from design strategies 1–3. Amino acid sequences of the two-chain constructs and corresponding DNA sequences of the synthetic gene fragments used for cloning. The peptide sequences are colored yellow.

| ID | Construct for nLuc plasmid, aa sequence | Construct for cLuc plasmid, aa sequence | Construct for nLuc plasmid, DNA sequence | Construct for cLuc plasmid, DNA sequence |
|---|---|---|---|---|
| 1-1 | HGKGSGGGSGGGSIEQLLTEVSKEIEQLLTEVSKEIEQLLTEVSKEGSGGSGGSGGSGGIEQLLTEVSKEIEQLLTEVSKEIEQLLTEVSKEGTKHHHHHH | PIQPPDSRAAATMIEQLLTEVSKEIEQLLTEVSKEIEQLLTEVSKEGGGGSGGGGSIEQLLTEVSKEIEQLLTEVSKEIEQLLTEVSKEGGGGSGGGGSTMTEKEIVDYVASQVTTAKKLRGGVVFVDEVPKGLTGKLDARKIREILIKAKKGGKIAVNSGTKHHHHHH | CATGGGAAAGGCTCTGGAGGGGGATCTGGAGGGGGATCCATTGAACAACTGCTTACAGAGGTGAGCAAGGAAATCGAGCAACTGCTTACCGAGGTGTCTAAAGAGATTGAGCAGCTCCTCACGGAGGTCTCAAAAGAAGGTTCCGGCGGCTCAGGAGGGAGTGGCGGATCTGGAGGAATTGAACAGCTCCTGACTGAAGTTAGTAAGGAGATAGAGCAGTTGCTGACGGAAGTGTCCAAAGAAATTGAGCAGCTCTTGACAGAAGTGAGTAAGGAAGGTACCAAGCACCACCATCACCATCACTAA | CCGATCCAGCCTCCGGACTCTAGAGCGGCCGCCACCATGATAGAACAGCTCCTGACAGAAGTGTCTAAGGAAATAGAGCAGCTGCTTACGGAGGTCTCCAAAGAAATTGAACAGCTCCTGACAGAGGTTTCCAAGGAGGGCGGAGGAGGCTCTGGTGGCGGTGGAAGTATAGAACAACTCCTGACCGAAGTCTCTAAAGAGATAGAGCAGTTGCTCACAGAAGTGTCAAAGGAGATTGAACAACTGCTTACTGAAGTGAGTAAGGAAGGTGGAGGGGGATCTGGCGGAGGCGGGTCAACAATGACAGAGAAAGAAATCGTTGACTACGTGGCTTCCCAGGTCACCACTGCCAAAAAACTCCGGGGCGGAGTTGTATTCGTGGACGAGGTCCCAAAAGGTTTGACCGGCAAACTGGACGCTAGAAAGATCAGAGAGATTCTGATTAAGGCCAAAAAGGGGGGGAAGATTGCCGTTAACTCTGGTACCAAGCACCACCATCACCATCACTAA |
| 1-2 | HGKGSGGGSGGGSIQGLLAEVGAQIQGLLAEVGAQIQGLLAEVGAQGSGGSGGSGGSGGIQGLLAEVGAQIQGLLAEVGAQIQGLLAEVGAQGTKHHHHHH | PIQPPDSRAAATMIQGLLAEVGAQIQGLLAEVGAQIQGLLAEVGAQGGGGSGGGGSIQGLLAEVGAQIQGLLAEVGAQIQGLLAEVGAQGGGGSGGGGSTMTEKEIVDYVASQVTTAKKLRGGVVFVDEVPKGLTGKLDARKIREILIKAKKGGKIAVNSGTKHHHHHH | CATGGGAAAGGCTCTGGAGGGGGATCTGGAGGGGGATCCATTCAGGGACTGCTCGCTGAGGTCGGCGCCCAGATTCAAGGATTGCTGGCAGAAGTTGGCGCCCAGATACAGGGACTTCTCGCGGAAGTCGGAGCTCAGGGCTCAGGTGGAAGTGGGGGATCAGGCGGTTCTGGAGGAATACAAGGCCTCCTGGCCGAGGTTGGAGCCCAAATCCAAGGGCTGCTGGCTGAGGTCGGGGCTCAGATTCAAGGGCTTCTGGCTGAAGTTGGCGCACAAGGTACCAAGCACCACCATCACCATCACTAA | CCGATCCAGCCTCCGGACTCTAGAGCGGCCGCCACCATGATCCAAGGGCTTCTCGCCGAAGTTGGGGCTCAAATTCAGGGCCTCTTGGCCGAAGTGGGAGCCCAAATCCAGGGGCTCCTGGCCGAAGTCGGGGCCCAGGGTGGGGGCGGGTCAGGGGGAGGGGGCAGTATACAGGGCCTGCTCGCCGAAGTCGGAGCTCAGATCCAGGGGCTCCTGGCGGAAGTAGGTGCCCAAATTCAGGGCCTCCTGGCAGAGGTAGGTGCACAAGGTGGCGGTGGATCTGGTGGTGGCGGTAGCACTATGACTGAGAAGGAGATTGTTGATTATGTGGCATCTCAGGTGACTACTGCGAAGAAGCTCAGGGGAGGAGTCGTCTTTGTAGATGAAGTGCCAAAGGGCCTCACCGGGAAGCTTGACGCCCGAAAAATCAGGGAGATTCTGATCAAGGCCAAAAAGGGCGGAAAGATCGCCGTTAACAGCGGTACCAAGCACCACCATCACCATCACTAA |
| 1-3 | HGKGSGGGSGGGSGEIIHSMQRLLLKSEGEIIHSMQRLLLKSEGSGGSGGSGGSGGGEIIHSMQRLLLKSEGEIIHSMQRLLLKSE | PIQPPDSRAAATMGEIIHSMQRLLLKSEGEIIHSMQRLLLKSEGGGGSGGGGSGEIIHSMQRLLLKSEGEIIHSMQRLLLKSEGGGGSGGGGSTMTEKEIVDYVASQVTTAKKLRGGVVFVDEVPKGLTGKLDARKIREILIKAKKGGKIAVNSGTKHHHHHH | CATGGGAAAGGCTCTGGAGGGGGATCTGGAGGGGGATCCGGAGAGATCATTCACAGTATGCAGAGACTGTTGCTCAAGAGCGAAGGAGAAATCATACACAGCATGCAGCGCCTCTTGCTCAAGTCCGAAGGCAGCGGCGGCTCCGGGGGGAGTGGGGGCAGTGGTGGCGGAGAAATTATTCACTCAATGCAGCGACTGCTGCTCAAATCTGAGGGCGAGATCATTCATTCCATGCAGAGGCTGCTGCTTAAATCCGAATAATAAGGTACCAAGCACCACCATCACCATCACTAATGAGAT | CCGATCCAGCCTCCGGACTCTAGAGCGGCCGCCACCATGGGCGAAATTATCCACTCTATGCAAAGATTGCTGCTCAAATCTGAGGGGGAAATAATTCATTCCATGCAGAGACTGCTGCTTAAGTCTGAAGGCGGTGGGGGCGGAGGGGGGTGGATCTGGCGAGATTATCCACTCAATGCAGAGGCTGCTCCTCAAATCCGAGGGGGAAATTATCCACAGTATGCAGAGGTTGCTGCTTAAATCTGAAGGCGGTGGAGGCTCTGGCGGTGGCGGATCTACCATGACTGAGAAGGAGATAGTGGATTACGTTGCCAGTCAGGTCACCACGGCTAAGAAGCTGCGAGGCGGGGTGGTCTTTGTTGATGAGGTTCCAAAGGGGCTGACTGGCAAACTGGACGCTAGGAAGATTAGAGAGATTCTCATCAAGGCTAAGAAGGGAGGTAAGATCGCTGTCAACTCTGGTACCAAGCACCACCATCACCATCACTAA |
| 1-4 | HGKGSGGGSGGGSLKSYMISSVDTAAKNLKSYMISSVDTAAKNGSGGSGGSGGSGGLKSYMISSVDTAAKNLKSYMISSVDTAAKN | PIQPPDSRAAATMLKSYMISSVDTAAKNLKSYMISSVDTAAKNGGGSGGGGSLKSYMISSVDTAAKNLKSYMISSVDTAAKNGGGGSGGGGSTMTEKEIVDYVASQVTTAKKLRGGVVFVDEVPKGLTGKLDARKIREILIKAKKGGKIAVNSGTKHHHHHH | CATGGGAAAGGCTCTGGAGGGGGATCTGGAGGGGGATCCCTCAAGTCCTACATGATTAGTAGCGTGGATACAGCGGCGAAGAACCTGAAGAGTTATATGATAAGCAGTGTCGATACCGCTGCAAAAAACGGATCTGGGGGAAGCGGTGGGTCCGGTGGCTCCGGGGGCCTGAAATCATATATGATTTCCAGTGTAGACACAGCCGCCAAAAATCTGAAAAGCTACATGATTTCCAGTGTGGACACCGCCGCTAAAAAATTAATAAGGTACCAAGCACCACCATCACCATCACTAATGAGAT | CCGATCCAGCCTCCGGACTCTAGAGCGGCCGCCACCATGCTGAAATCATATATGATCAGCTCAGTTGATACAGCCGCCAAGAACTTGAAGTCCTACATGATCTCCTCCGTGGATACCGCTGCCAAGAATGGAGGAGGTGGTTCAGGAGGAGGAGGGAAGTCTCAAGTCCTATATGATCTCCTCAGTCGATACTGCTGCGAAAAATTTGAAGTCTTACATGATTTCCTCTGTTGACACAGCCGCTAAGAACGGGGGAGGGGGATCTGGCGGTGGAGGAAGTACGATGACTGAGAAAGAAATTGTGGACTACGTGGCATCACAGGTGACAACAGCAAAGAAACTTAGGGGAGGCGTGGTTTTCGTGGACGAAGTACCCAAAGGGTTGACAGGCAAACTGGATGCCAGGAAGATACGAGAGATACTGATCAAAGCTAAAAAGGGCGGCAAGATCGCCGTCAATAGCGGTACCAAGCACCACCATCACCATCACTAA |
| 1-5 | HGKGSGGGSGGGSANMRNIVHFVKESEEANMRNIVHFVKESEEGSGGSGGSGGSGGANMRNIVHFVKESEEANMRNIVHFVKESEE | PIQPPDSRAAATMANMRNIVHFVKESEEANMRNIVHFVKESEEGGGSGGGGSANMRNIVHFVKESEEANMRNIVHFVKESEEGGGGSGGGGSTMTEKEIVDYVASQVTTAKKLRGGVVFVDEVPKGLTGKLDARKIREILIKAKKGGKIAVNSGTKHHHHHH | CATGGGAAAGGCTCTGGAGGGGGATCTGGAGGGGGATCCGCTAATATGAGGAATATAGTGCACTTCGTGAAGGAATCCGAGGAAGCTAACATGCGGAACATCGTGCACTTTGTTAAGGAAAGCGAAGAGGGATCTGGGGGCTCAGGTGGGAGTGGGGGGTCTGGCGGCGCAAATATGCGGAATATCGTGCACTTTGTGAAGGAAAGTGAAGAGGCTAACATGCGAAACATTGTTCATTTTGTGAAAGAGAGTGAGGAATAATAAGGTACCAAGCACCACCATCACCATCACTAATGAGAT | CCGATCCAGCCTCCGGACTCTAGAGCGGCCGCCACCATGGCTAACATGCGGAACATCGTGCATTTCGTCAAGGAGAGCGAAGAAGCAAATATGCGGAATATTGTGCATTTCGTGAAAGAAAGCGAGGAAGGAGGAGGCGGAAGCGGAGGCGGGGGCTCTGCTAATATGCGCAATATTGTTCACTTCGTGAAAGAGTCTGAAGAAGCCAACATGCGGAACATTGTGCACTTTGTGAAGGAATCCGAGGAGGGTGGGGGTGGAAGCGGGGGTGGAGGCAGCACCATGACTGAAAAAGAAATCGTCGACTACGTTGCCTCACAGGTTACAACCGCCAAAAAACTGAGAGGTGGCGTAGTCTTCGTCGACGAGGTTCCCAAGGGCCTCACAGGCAAGTTGGATGCCAGAAAGATTAGAGAGATTTTGATTAAAGCGAAAAAGGGGGGGAAAATTGCAGTGAACAGTGGTACCAAGCACCACCATCACCATCACTAA |

| | | | | |
|---|---|---|---|---|
| 1-6 | HGKGSGGGSGGGSSSRFMKETGKREMWYSSRFMKETGKREMWYGSGGSGGSGGSGGSSRFMKETGKREMWYSSRFMKETGKREMWY | PIQPPDSRAAATMSSRFMKETGKREMWYSSRFMKETGKREMWYGGGGSGGGGSSSRFMKETGKREMWYSSRFMKETGKREMWYGGGGSGGGGSTMTEKEIVDYVASQVTTAKKLRGGVVFVDEVPKGLTGKLDARKIREILIKAKKGGKIAVNSGTKHHHHHH | CATGGGAAAGGCTCTGGAGGGGGATCTGGAGGGGGATCCTCCTCACGATTTATGAAGGAAACAGGTAAAAGGGAAATGTGGTATAGCAGCCGGTTTATGAAGGAGACTGGCAAGCGCGAGATGTGGTACGGCTCAGGCGGTAGTGGCGGCTCCGGAGGAAGCGGGGGTTCCTCTAGGTTTATGAAGGAGACCGGGAAGAGAGAAATGTGGTATTCCAGCAGATTTATGAAGGAGACAGGGAAACGGGAAATGTGGTACTAATAAGGTACCAAGCACCACCATCACCATCACTAATGAGAT | CCGATCCAGCCTCCGGACTCTAGAGCGGCCGCCACCATGTCTAGCCGGTTTATGAAGGAGACAGGAAAGAGGGAGATGTGGTATTCTTCCAGATTCATGAAGGAGACAGGCAAGAGGGAGATGTGGTACGGAGGGGGAGGTAGCGGGGGCGGGGGCAGCAGTAGTAGGTTCATGAAAGAGACAGGCAAGCGGGAGATGTGGTATAGTTCTAGGTTCATGAAAGAGACAGGGAAACGAGAAATGTGGTACGGGGGTGGCGGTTCTGGAGGCGGTGGATCTACCATGACTGAGAAAGAGATTGTTGACTATGTCGCCTCCCAGGTCACCACTGCCAAAAAGTTGAGGGGTGGCGTCGTGTTCGTCGACGAAGTCCCAAAAGGGCTGACAGGCAAACTTGACGCGAGGAAGATTAGGGAAATTTTGATCAAGGCTAAAAAAGGCGGAAAGATCGCCGTGAATTCTGGTACCAAGCACCACCATCACCATCACTAA |
| 1-7 | HGKGSGGGSGGGSINEMLEKVADEINEMLEKVADEINEMLEKVADEGGGGSGGGGSINEMLEKVADEINEMLEKVADEINEMLEKVADE | PIQPPDSRAAATMINEMLEKVADEINEMLEKVADEINEMLEKVADEGGSGGGGSGGGGSINEMLEKVADEINEMLEKVADEINEMLEKVADEGGSGGGGSGGGGSTMTEKEIVDYVASQVTTAKKLRGGVVFVDEVPKGLTGKLDARKIREILIKAKKGGKIAVNSGTKHHHHHH | CATGGGAAAGGCTCTGGAGGGGGATCTGGAGGGGGATCCATCAACGAGATGTTGGAGAAGGTCGCCGACGAGATAAACGAGATGCTCGAGAAAGTGGCTGATGAGATCAACGAAATGCTGGAGAAGGTAGCGGACGAAGGCGGTGGAGGCTCTGGTGGCGGCGGCAGCATTAACGAAATGCTTGAAAAAGTCGCTGATGAGATTAATGAGATGCTGGAGAAGGTTGCTGACGAGATCAATGAAATGTTGGAGAAGGTTGCGGACGAATAATAAGGTACCAAGCACCACCATCACCATCACTAA | CCGATCCAGCCTCCGGACTCTAGAGCGGCCGCCACCATGATAAATGAAATGCTGGAAAAAGTTGCGGACGAGATAAATGAAATGCTGGAGAAAGTGGCCGACGAGATTAACGAAATGCTCGAAAAGGTCGCCGATGAGGGTGGGTCTGGAGGCGGAGGCAGCGGCGGCGGGGGCAGTATCAACGAGATGCTCGAAAAGGTCGCAGATGAGATCAACGAAATGCTGGAAAAGGTGGCCGACGAGATCAATGAAATGCTTGAGAAGGTAGCTGATGAAGGCGGGTCCGGCGGCGGCGGATCTGGAGGCGGTGGCAGCACGATGACTGAAAAGGAAATTGTGGACTACGTAGCATCCCAAGTGACCACAGCAAAGAAATTGAGAGGCGGGGTCGTCTTTGTAGACGAGGTGCCTAAAGGGCTGACAGGCAAGCTGGATGCTCGCAAAATCCGGGAAATCCTGATAAAAGCCAAAAAGGGAGGTAAGATTGCCGTAAACAGCGGTACCAAGCACCACCATCACCATCACTAA |
| 1-8 | HGKGSGGGSGGGSMRELITVAEAHMRELITVAEAHMRELITVAEAHGGGGSGGGGSMRELITVAEAHMRELITVAEAHMRELITVAEAH | PIQPPDSRAAATMMRELITVAEAHMRELITVAEAHMRELITVAEAHGGSGGGGSGGGGSMRELITVAEAHMRELITVAEAHMRELITVAEAHGGSGGGGSGGGGSTMTEKEIVDYVASQVTTAKKLRGGVVFVDEVPKGLTGKLDARKIREILIKAKKGGKIAVNSGTKHHHHHH | CATGGGAAAGGCTCTGGAGGGGGATCTGGAGGGGGATCCATGCGGGAACTGATCACAGTTGCTGAGGCTCACATGCGGGAGCTGATAACCGTCGCGGAGGCACATATGCGCGAACTGATTACCGTGGCTGAAGCGCACGGCGGAGGTGGCAGCGGCGGTGGCGGTTCAATGCGGGAGCTCATCACGGTGGCGGAAGCACACATGAGGGAGCTTATAACTGTGGCGGAAGCCCACATGAGAGAACTCATCACAGTAGCCGAAGCACACTAATAAGGTACCAAGCACCACCATCACCATCACTAA | CCGATCCAGCCTCCGGACTCTAGAGCGGCCGCCACCATGATGAGGGAACTCATTACGGTGGCCGAAGCACACATGAGAGAACTTATCACTGTTGCCGAAGCCCACATGCGGGAATTGATTACCGTCGCTGAAGCCCATGGCGGCAGCGGAGGTGGTGGGAGCGGCGGAGGTGGAAGTATGCGCGAGCTGATCACTGTTGCGGAGGCTCATATGCGAGAGCTCATCACAGTGGCTGAGGCCCATATGCGCGAACTTATAACTGTTGCCGAAGCCCATGGTGGAAGTGGCGGCGGCGGCTCCGGGGGCGGAGGATCTACCATGACCGAGAAAGAGATCGTGGACTACGTGGCTTCCCAGGTTACCACCGCGAAAAAGTTGCGGGGCGGCGTGGTGTTCGTTGATGAGGTGCCTAAAGGCCTCACTGGAAAGCTGGACGCACGGAAAATTAGGGAGATTCTGATTAAGGCTAAGAAAGGGGGTAAGATCGCGGTTAATTCCGGTACCAAGCACCACCATCACCATCACTAA |
| 1-9 | HGKGSGGGSGGGSISTLLEKVRKEISTLLEKVRKEISTLLEKVRKEGGGGSGGGGSISTLLEKVRKEISTLLEKVRKEISTLLEKVRKE | PIQPPDSRAAATMISTLLEKVRKEISTLLEKVRKEISTLLEKVRKEGGSGGGGSGGGGSISTLLEKVRKEISTLLEKVRKEISTLLEKVRKEGGSGGGGSGGGGSTMTEKEIVDYVASQVTTAKKLRGGVVFVDEVPKGLTGKLDARKIREILIKAKKGGKIAVNSGTKHHHHHH | CATGGGAAAGGCTCTGGAGGGGGATCTGGAGGGGGATCCATCTCAACTCTTCTTGAAAAGGTCAGAAAGGAAATATCTACGCTGCTGGAGAAGGTGCGGAAAGAGATTTCCACTCTCCTTGAGAAGGTGAGGAAAGAAGGAGGGGGTGGTAGCGGCGGTGGGGGCTCCATCTCAACCCTGCTTGAGAAGGTCAGGAAAGAGATCAGCACACTGCTCGAAAAAGTACGCAAAGAGATATCTACACTGCTCGAAAAAGTCCGAAAGGAGTAATAAGGTACCAAGCACCACCATCACCATCACTAA | CCGATCCAGCCTCCGGACTCTAGAGCGGCCGCCACCATGATTTCCACCCTCCTGGAGAAAGTTAGGAAAGAAATAAGCACTCTGTTGGAGAAAGTGCGGAAGGAGATCTCAACCCTTCTCGAGAAGGTTCGAAAGGAGGGAGGCAGCGGGGGAGGTGGGTCTGGCGGAGGTGGGAGTATTTCCACTCTGCTGGAGAAGGTCCGGAAGGAAATCTCCACTCTCCTTGAGAAAGTGAGGAAAGAAATCAGTACTTTGCTGGAAAAGGTAAGGAAGGAGGGGGGCTCTGGGGGCGGTGGTTCAGGTGGCGGCGGAAGTACCATGACTGAGAAGGAGATAGTCGATTACGTGGCCTCTCAGGTGACAACTGCAAAGAAGTTGCGCGGGGGTGTAGTGTTCGTTGATGAGGTTCCTAAGGGGCTTACTGGCAAGTTGGATGCCCGGAAAATCAGGGAGATCCTGATTAAGGCCAAGAAGGGAGGCAAGATTGCCGTGAACAGTGGTACCAAGCACCACCATCACCATCACTAA |
| 1-10 | EHGKGSGGGSGGGSHFEWKEVMEKFHTRMHFEWKEVMEKFHTRMGGSGGGGSGGGGSHFEWKEVMEKFHTRMHFEWKEVMEKFHTRM | PIQPPDSRAAATMHFEWKEVMEKFHTRMHFEWKEVMEKFHTRMGGSGGGGSGGGGSHFEWKEVMEKFHTRMHFEWKEVMEKFHTRMGGSGGGGSGGGGSTMTEKEIVDYVASQVTTAKKLRGGVVFVDEVPKGLTGKLDARKIREILIKAKKGGKIAVNSGTKHHHHHH | GAGCATGGGAAAGGCTCTGGAGGGGGATCTGGAGGGGGATCCCACTTTGAATGGAAAGAGGTCATGGAGAAGTTCCATACCCGGATGCACTTTGAATGGAAAGAGGTGATGGAAAAATTTCATACTAGAATGGGCGGATCAGGAGGCGGCGGGAGCGGAGGCGGAGGGTCACACTTTGAGTGGAAGGAAGTCATGGAGAAATTTCACACTAGGATGCATTTCGAATGGAAGGAGGTCATGGAGAAATTCCACACTCGGATGTGATAATAAGGTACCAAGCACCACCATCACCATCACTAATGA | CCGATCCAGCCTCCGGACTCTAGAGCGGCCGCCACCATGCACTTCGAATGGAAGGAGGTCATGGAAAAATTTCACACCAGGATGCATTTCGAGTGGAAGGAAGTCATGGAGAAATTCCATACAAGAATGGGCGGAAGCGGCGGTGGTGGTTCTGGGGGAGGAGGATCTCACTTCGAGTGGAAGGAAGTGATGGAAAAGTTCCATACTCGGATGCATTTCGAATGGAAAGAGGTCATGGAGAAATTCCACACCCGGATGGGGGGCTCCGGTGGGGGAGGGAGCGGGGGTGGCGGGTCCACCATGACCGAGAAGGAAATTGTCGACTATGTGGCTAGTCAGGTTACCACCGCCAAGAAGCTCAGGGGGGGAGTCGTATTCGTCGATGAGGTGCCAAAAGGTCTGACAGGCAAGCTGGATGCACGGAAGATAAGGGAAATCCTGATCAAAGCAAAGAAAGGTGGAAAAATTGCCGTCAATAGTGGTACCAAGCACCACCATCACCATCACTAA |

| | | | | |
|---|---|---|---|---|
| 1-11 | KGSGGGSGGGSSETWHRLQEFYWGNYSETWHRLQEFYWGNYSETWHRLQEFYWGNYGGGGSGGGGSSETWHRLQEFYWGNYSETWHRLQEFYWGNYSETWHRLQEFYWGNYGGGGSGGGGSSETWHRLQEFYWGNYSETWHRLQEFYWGNYSETWHRLQEFYWGNYGTKHHHHHH | PIQPPDSRAAATMSETWHRLQEFYWGNYSETWHRLQEFYWGNYSETWHRLQEFYWGNYGGGGSGGGGSSETWHRLQEFYWGNYSETWHRLQEFYWGNYGGGGSGGGGSSETWHRLQEFYWGNYSETWHRLQEFYWGNYSETWHRLQEFYWGNYGGGGSGGGGSTMTEKEIVDYVASQVTTAKKLRGGVVFVDEVPKGLTGKLDARKIREILIKAKKGGKIAVNSGTKHHHHHH | AAAGGCTCTGGAGGGGGATCTGGAGGGGGATCCAGTGAGACCTGGCATCGCCTCCAAGAATTTTACTGGGGGAATTATAGCGAGACGTGGCACAGACTGCAGGAGTTCTATTGGGGAAACTACTCAGAGACATGGCACCGACTGCAGGAATTTTATTGGGGTAATTATGGCGGGGGAGGTAGTGGTGGCGGTGGAAGTAGCGAAACATGGCATAGGCTGCAGGAGTTTTATTGGGGCAATTATAGCGAAACATGGCACCGCCTTCAAGAGTTCTATTGGGGCAACTACAGTGAGACCTGGCATCGCCTGCAGGAATTCTATTGGGGAAATTACGGTGGTGGTGGCAGCGGCGGCGGCGGTTCTAGCGAAACCTGGCATAGACTCCAGGAGTTCTATTGGGGCAACTACTCTGAGACTTGGCATAGGCTGCAGGAATTTTACTGGGGCAATTATTCCGAGACCTGGCACCGACTTCAGGAGTTCTATTGGGGAAACTACGGTACCAAGCACCACCATCACCATCACTAA | CCGATCCAGCCTCCGGACTCTAGAGCGGCCGCCACCATGTCTGAAACGTGGCACAGACTGCAGGAGTTCTATTGGGGTAACTACAGTGAGACTTGGCACAGGCTGCAGGAGTTCTACTGGGGGAACTATTCAGAGACCTGGCACCGGCTTCAGGAGTTCTACTGGGGCAATTACGGCGGCGGTGGTTCAGGCGGTGGAGGTTCTTCCGAAACATGGCATAGGCTCCAGGAATTCTACTGGGGGAATTACTCTGAGACATGGCATAGGCTGCAGGAGTTTTATTGGGGGAACTATAGTGAAACCTGGCATAGGTTGCAGGAATTCTACTGGGGGAATTACGGAGGTGGAGGCAGTGGCGGCGGCGGCAGCTCCGAGACCTGGCACCGCCTGCAGGAGTTTTACTGGGGAAACTACAGCGAGACTTGGCATCGCCTCCAAGAGTTCTACTGGGGGAATTATTCCGAAACTTGGCATCGGCTTCAGGAGTTCTACTGGGGAAACTATGGCGGGGGAGGCTCAGGAGGCGGCGGAAGTACCATGACGGAGAAGGAGATTGTTGATTATGTGGCATCCCAGGTCACTACCGCAAAAAAGCTGAGGGGCGGGGTGGTGTTTGTCGACGAAGTGCCCAAGGGGCTTACTGGTAAGTTGGATGCTCGAAAAATCAGGGAAATTCTGATCAAGGCCAAAAAAGGGGGGAAGATAGCGGTGAATTCCGGTACCAAGCACCACCATCACCATCACTAA |
| 1-12 | KGSGGGSGGGSGDDLDNLKRGLGDDLDNLKRGLGDDLDNLKRGLGGGGSGGGGSGDDLDNLKRGLGDDLDNLKRGLGDDLDNLKRGLGGGGSGGGGSGDDLDNLKRGLGDDLDNLKRGLGDDLDNLKRGLGTKHHHHHH | PIQPPDSRAAATMGDDLDNLKRGLGDDLDNLKRGLGDDLDNLKRGLGGGGSGGGGSGDDLDNLKRGLGDDLDNLKRGLGDDLDNLKRGLGGGGSGGGGSGDDLDNLKRGLGDDLDNLKRGLGDDLDNLKRGLGGGGSGGGGSTMTEKEIVDYVASQVTTAKKLRGGVVFVDEVPKGLTGKLDARKIREILIKAKKGGKIAVNSGTKHHHHHH | AAAGGCTCTGGAGGGGGATCTGGAGGGGGATCCGGAGATGACCTGGACAATCTGAAGCGCGGGCTCGGCGACGACCTTGATAATCTCAAAAGGGGCCTCGGTGATGACCTGGACAATCTTAAACGGGGGCTCGGCGGGGGTGGGAGTGGGGGCGGCGGCAGTGGCGATGACCTGGACAATCTGAAGCGGGGCCTCGGAGACGACTTGGACAATTTGAAAAGAGGTCTGGGTGACGACCTGGATAATTTGAAGCGGGGGCTCGGGGGTGGCGGTTCTGGCGGTGGAGGCTCAGGCGATGATCTCGACAACCTCAAGCGCGGACTCGGAGACGACCTTGATAATTTGAAGAGGGGACTTGGCGATGATCTTGATAACCTTAAGCGGGGACTGGGTACCAAGCACCACCATCACCATCACTAA | CCGATCCAGCCTCCGGACTCTAGAGCGGCCGCCACCATGGGGGACGACCTTGATAATCTTAAACGAGGACTCGGGGACGATCTGGATAACTTGAAACGCGGGCTGGGAGACGATTTGGATAACCTCAAGCGCGGGTTGGGTGGAGGGGGTTCAGGGGGCGGCGGTTCCGGCGATGATTTGGATAATCTGAAGCGAGGGCTGGGTGATGACCTTGACAACCTGAAGAGGGGACTTGGTGACGACCTGGACAATTTGAAGCGGGGACTGGGGGGTGGCGGCTCCGGCGGCGGTGGTTCTGGTGATGACCTGGATAACCTGAAACGGGGGCTCGGGGACGACCTGGACAATCTCAAAAGGGGGCTGGGGGACGACCTGGACAATCTGAAGAGGGGTCTGGGGGGAGGAGGTTCTGGCGGAGGCGGTTCCACCATGACTGAGAAAGAAATAGTCGATTATGTCGCAAGTCAGGTGACGACCGCCAAGAAGCTGAGGGGAGGGGTCGTCTTTGTGGACGAGGTGCCGAAAGGGTTGACAGGCAAACTGGATGCCCGCAAAATACGGGAAATTCTGATTAAGGCAAAGAAAGGCGGCAAAATCGCCGTAAACAGTGGTACCAAGCACCACCATCACCATCACTAA |
| 1-13 | KGSGGGSGGGSSAELGREWIEDSAELGREWIEDSAELGREWIEDGGGGSGGGGSSAELGREWIEDSAELGREWIEDGGGGSGGGGSSAELGREWIEDSAELGREWIEDSAELGREWIEDGTKHHHHHH | PIQPPDSRAAATMSAELGREWIEDSAELGREWIEDSAELGREWIEDGGGGSGGGGSSAELGREWIEDSAELGREWIEDSAELGREWIEDGGGGSGGGGSSAELGREWIEDSAELGREWIEDSAELGREWIEDGGGGSGGGGSTMTEKEIVDYVASQVTTAKKLRGGVVFVDEVPKGLTGKLDARKIREILIKAKKGGKIAVNSGTKHHHHHH | AAAGGCTCTGGAGGGGGATCTGGAGGGGGATCCAGTGCAGAGCTTGGAAGGGAGTGGATCGAGGACTCTGCAGAGCTGGGGCGCGAGTGGATTGAGGATAGCGCGGAGCTCGGTCGGGAGTGGATCGAAGATGGGGGCGGGGGCAGCGGGGGAGGCGGGTCCAGTGCAGAACTGGGCCGGGAGTGGATCGAAGACTCCGCAGAACTCGGGCGAGAATGGATCGAAGACAGCGCAGAACTCGGCCGCGAATGGATAGAGGATGGCGGCGGCGGATCTGGGGGTGGTGGAAGTAGCGCCGAGCTCGGTCGCGAGTGGATTGAGGACTCCGCTGAACTGGGACGCGAGTGGATCGAAGATAGTGCTGAGCTCGGAAGAGAGTGGATTGAGGACGGTACCAAGCACCACCATCACCATCACTAA | CCGATCCAGCCTCCGGACTCTAGAGCGGCCGCCACCATGAGCGCTGAACTGGGTAGAGAGTGGATTGAAGACTCCGCCGAACTGGGCAGAGAGTGGATCGAGGATTCCGCCGAGCTGGGCCGAGAATGGATCGAAGACGGAGGTGGCGGGTCTGGCGGAGGCGGCAGCTCAGCCGAACTCGGCCGGGAATGGATAGAAGACAGCGCTGAGCTCGGGCGAGAGTGGATTGAGGACAGTGCCGAGCTGGGGAGGGAGTGGATTGAAGATGGCGGGGGGGGCTCCGGCGGCGGCGGCTCATCCGCCGAGCTTGGCAGGGAGTGGATCGAGGATTCAGCAGAGCTTGGAAGGGAGTGGATTGAGGACTCTGCAGAATTGGGCCGCGAATGGATCGAAGACGGCGGGGGCGGTTCTGGGGGTGGAGGATCTACAATGACCGAAAAGGAGATTGTCGACTATGTCGCATCACAGGTGACAACAGCAAAAAAGCTGCGCGGGGGGGTCGTTTTTGTCGATGAGGTCCCAAAGGGGCTCACTGGGAAGCTGGACGCCAGAAAGATCCGCGAGATTCTCATCAAGGCAAAAAAAGGAGGTAAAATCGCTGTTAACTCAGGTACCAAGCACCACCATCACCATCACTAA |
| 1-2.1 | KGSGGGSGGGSIQGLLAEVGAQIQGLLAEVGAQIQGLLALVGAQGGGGSGGGGSIQGLLAEVGAQIQGLLAEVGAQIQGLLALVGAQ | PIQPPDSRAAATMIQGLLAEVGAQIQGLLAEVGAQIQGLLALVGAQGGGGSGGGGSIQGLLAEVGAQIQGLLAEVGAQIQGLLALVGAQGGGGSGGGGSTMTEKEIVDYVASQVTTAKKLRGGVVFVDEVPKGLTGKLDARKIREILIKAKKGGKIAVNSGTKHHHHHH | AAAGGCTCTGGAGGGGGATCTGGAGGGGGATCCATCCAGGGGCTTCTGGCTGAGGTCGGCGCCCAGATTCAAGGGCTGTTGGCTGAAGTGGGCGCTCAGATTCAGGGGCTGCTCGCTCTTGTTGGTGCACAGGGAGGCGGCGGCTCAGGGGGAGGCGGTTCCATACAAGGGCTTCTGGCCGAAGTGGGAGCACAGATTCAAGGCCTGCTGGCGGAAGTCGGCGCTCAGATACAAGGCTTGCTGGCCTTGGTTGGAGCTCAGTAATAAGGTACCAAGCACCACCATCACCATCACTAATGAGAT | CCGATCCAGCCTCCGGACTCTAGAGCGGCCGCCACCATGATTCAGGGTTTGCTGGCAGAGGTGGGTGCCCAGATCCAGGGACTGCTGGCCGAGGTAGGCGCTCAAATTCAAGGCCTTCTCGCCTTGGTAGGAGCACAGGGTGGGGGCGGGTCCGGGGGCGGTGGGAGCATACAGGGGCTGCTGGCAGAAGTCGGCGCGCAAATCCAGGGCCTTCTCGCCGAGGTCGGTGCACAGATTCAGGGTCTGTTGGCACTGGTAGGCGCACAAGGTGGAGGCGGGTCCGGAGGGGGAGGATCAACTATGACCGAAAAAGAGATCGTCGATTACGTGGCCAGTCAGGTTACCACAGCCAAAAAACTGCGCGGCGGGGTGGTGTTTGTAGACGAGGTGCCTAAGGGGTTGACTGGCAAACTGGACGCCAGGAAAATTCGCGAAATCCTGATCAAGGCAAAAAAGGCGGAAAGATTGCAGTCAACTCAGGTACCAAGCACCACCATCACCATCACTAA |

| | | | | |
|---|---|---|---|---|
| 1-2.2 | KGSGGGSGGGSIQGLLAR<br>VGAQIQGLLAEVGAQIQG<br>LLAEVGAQGGGGSGGGGS<br>IQGLLARVGAQIQGLLAE<br>VGAQIQGLLAEVGAQ | PIQPPDSRAAATMIQGLLARVG<br>AQIQGLLAEVGAQIQGLLAEVG<br>AQGGGGSGGGGSIQGLLARVGA<br>QIQGLLAEVGAQIQGLLAEVGA<br>QGGGGSGGGGSTMTEKEIVDYV<br>ASQVTTAKKLRGGVVFVDEVPK<br>GLTGKLDARKIREILIKAKKGG<br>KIAVNSGTKHHHHHH | AAAGGCTCTGGAGGGGGATCTGGAGGGGGATCCATC<br>CAGGGACTTTTGGCCCGGGTTGGGGCGCAGATCCAG<br>GGCCTTCTCGCGGAGGTTGGCGCTCAGATACAAGGA<br>TTGCTGGCCGAAGTGGGCGCCCAGGGTGGGGGCGGC<br>AGCGGGGGTGGCGGGAGCATACAGGGCCTTCTGGCA<br>AGGGTAGGTGCTCAGATTCAAGGGCTTCTGGCCGAG<br>GTGGGCGCTCAAATACAGGGCCTGCTCGCCGAAGTG<br>GGGGCACAATAATAAGGTACCAAGCACCACCATCAC<br>CATCACTAATGAGAT | CCGATCCAGCCTCCGGACTCTAGAGCGGCCGCCACCATGATCCAGGGTCTGTTGGCACGAGTGGGCG<br>CACAGATCCAGGGTCTCCTCGCCGAGGTGGGCGCCCAGATTCAGGGACTGCTGGCAGAAGTCGGAGC<br>TCAGGGAGGAGGGGGAAGCGGTGGTGGCGGCTCAATCCAGGGCCTGCTTGCACGAGTGGGTGCCCAG<br>ATACAGGGCCTTCTCGCTGAAGTGGGGGCTCAGATTCAAGGCCTGTTGGCGGAGGTGGGAGCCCAGG<br>GCGGAGGGGGCTCAGGCGGCGGTGGATCTACCATGACAGAGAAGGAAATTGTGGACTATGTCGCCAG<br>CCAGGTGACCACAGCGAAAAAACTTAGAGGGGGTGTTGTGTTCGTTGACGAAGTTCCCAAGGGGCTT<br>ACAGGCAAGCTGGATGCAAGGAAGATCAGAGAGATCCTCATTAAGGCGAAAAAAGGCGGCAAAATCG<br>CAGTCAACTCTGGTACCAAGCACCACCATCACCATCACTAA |
| 1-2.3 | KGSGGGSGGGSIQGLLAV<br>VGAQIQGLLAEVGAQIQG<br>LLAEVGAQGGGGSGGGGS<br>IQGLLAVVGAQIQGLLAE<br>VGAQIQGLLAEVGAQ | PIQPPDSRAAATMIQGLLAVVG<br>AQIQGLLAEVGAQIQGLLAEVG<br>AQGGGGSGGGGSIQGLLAVVGA<br>QIQGLLAEVGAQIQGLLAEVGA<br>QGGGGSGGGGSTMTEKEIVDYV<br>ASQVTTAKKLRGGVVFVDEVPK<br>GLTGKLDARKIREILIKAKKGG<br>KIAVNSGTKHHHHHH | AAAGGCTCTGGAGGGGGATCTGGAGGGGGATCCATT<br>CAGGGCCTGCTGGCAGTCGTTGGAGCACAAATCCAG<br>GGCCTCCTGGCTGAAGTGGGAGCCCAAATCCAGGGG<br>TTGCTGGCTGAAGTGGGTGCACAGGGAGGTGGGGGG<br>TCAGGAGGAGGCGGCTCCATCCAGGGTCTTCTCGCC<br>GTTGTGGGAGCTCAGATTCAAGGCCTCCTCGCAGAA<br>GTAGGCGCCCAAATCCAAGGGCTGCTTGCTGAAGTG<br>GGCGCCCAGTAATAAGGTACCAAGCACCACCATCAC<br>CATCACTAATGAGAT | CCGATCCAGCCTCCGGACTCTAGAGCGGCCGCCACCATGATCCAGGGCTTGTTGGCTGTGGTGGGCG<br>CTCAGATCCAGGGCCTTCTTGCTGAGGTTGGAGCGCAGATTCAAGGGCTCCTGGCCGAAGTTGGGGC<br>TCAGGGCGGAGGTGGATCTGGAGGAGGTGGTTCCATTCAGGGACTGTTGGCAGTTGTTGGCGCCCAG<br>ATTCAAGGGTTGCTTGCAGAAGTGGGAGCACAGATTCAGGGACTGCTTGCAGAGGTCGGCGCACAAG<br>GAGGCGGAGGATCTGGTGGTGGAGGAAGTACAATGACCGAAAAAGAAATTGTAGATTATGTTGCCTC<br>ACAGGTGACGACAGCCAAGAAACTTCGGGGCGGTGTGGTCTTCGTGGATGAGGTCCCCAAGGGTCTC<br>ACCGGGAAGCTGGACGCACGCAAGATAAGAGAGATTCTGATCAAGGCCAAAAAGGGTGGCAAGATCG<br>CCGTCAACAGCGGTACCAAGCACCACCATCACCATCACTAA |
| 1-2.4 | KGSGGGSGGGSIQGLLAE<br>VGAQIQALLAEVGAQIQG<br>LLAEVGAQGGGGSGGGGS<br>IQGLLAEVGAQIQALLAE<br>VGAQIQGLLAEVGAQ | PIQPPDSRAAATMIQGLLAEVG<br>AQIQALLAEVGAQIQGLLAEVG<br>AQGGGGSGGGGSIQGLLAEVGA<br>QIQALLAEVGAQIQGLLAEVGA<br>QGGGGSGGGGSTMTEKEIVDYV<br>ASQVTTAKKLRGGVVFVDEVPK<br>GLTGKLDARKIREILIKAKKGG<br>KIAVNSGTKHHHHHH | AAAGGCTCTGGAGGGGGATCTGGAGGGGGATCCATC<br>CAGGGGCTTCTTGCTGAGGTGGGCGCCCAGATCCAG<br>GCACTGCTGGCCGAGGTAGGAGCCCAAATACAGGGC<br>CTGCTCGCCGAGGTCGGGGCACAAGGTGGCGGCGGG<br>TCAGGAGGGGGAGGAAGCATTCAGGGTCTCCTGGCA<br>GAAGTGGGAGCTCAAATCCAGGCACTGCTCGCTGAG<br>GTTGGGGCTCAGATCCAGGGCCTCCTTGCCGAGGTT<br>GGCGCCCAATAATAAGGTACCAAGCACCACCATCAC<br>CATCACTAATGAGAT | CCGATCCAGCCTCCGGACTCTAGAGCGGCCGCCACCATGATACAGGGCCTGCTGGCAGAGGTAGGTG<br>CTCAGATCCAAGCTCTCCTTGCTGAAGTGGGGGCTCAAATCCAGGGGCTCCTGGCTGAGGTAGGAGC<br>TCAGGGTGGAGGAGGATCTGGGGGCGGAGGTTCAATTCAGGGACTCCTTGCCGAAGTGGGGGCTCAG<br>ATTCAGGCCCTGTTGGCAGAAGTTGGAGCGCAGATTCAAGGGCTGCTGGCCGAAGTTGGCGCTCAGG<br>GAGGTGGCGGCTCCGGAGGGGGTGGTAGTACTATGACAGAGAAGGAAATAGTCGATTACGTCGCTAG<br>CCAGGTCACTACCGCTAAAAAACTTAGAGGTGGAGTGGTCTTTGTGGACGAAGTGCCCAAAGGCTTG<br>ACAGGGAAGCTCGACGCCCGCAAGATTCGAGAAATCCTCATCAAGGCCAAGAAGGGGGGCAAAATTG<br>CAGTGAACTCCGGTACCAAGCACCACCATCACCATCACTAA |
| 1-2.5 | KGSGGGSGGGSIQGLLAE<br>VGAQIQGLLAEVGAQIQV<br>LLAEVGAQGGGGSGGGGS<br>IQGLLAEVGAQIQGLLAE<br>VGAQIQVLLAEVGAQ | PIQPPDSRAAATMIQGLLAEVG<br>AQIQGLLAEVGAQIQVLLAEVG<br>AQGGGGSGGGGSIQGLLAEVGA<br>QIQGLLAEVGAQIQVLLAEVGA<br>QGGGGSGGGGSTMTEKEIVDYV<br>ASQVTTAKKLRGGVVFVDEVPK<br>GLTGKLDARKIREILIKAKKGG<br>KIAVNSGTKHHHHHH | AAAGGCTCTGGAGGGGGATCTGGAGGGGGATCCATC<br>CAGGGCCTGTTGGCAGAGGTCGGCGCTCAAATACAG<br>GGGCTTCTGGCAGAGGTAGGAGCGCAGATTCAGGTC<br>CTGCTGGCTGAAGTAGGAGCTCAAGGAGGTGGGGGC<br>AGCGGAGGTGGCGGCTCTATTCAGGGCCTGTTGGCC<br>GAAGTCGGGGCACAGATTCAGGGACTTTTGGCCGAA<br>GTGGGAGCTCAGATCCAGGTTCTGCTTGCCGAGGTC<br>GGTGCCCAATAATAAGGTACCAAGCACCACCATCAC<br>CATCACTAATGAGAT | CCGATCCAGCCTCCGGACTCTAGAGCGGCCGCCACCATGATCCAGGGACTGCTGGCCGAGGTGGGCG<br>CACAGATTCAGGGGCTGCTGGCTGAGGTTGGGGCTCAGATACAGGTGCTCTTGGCTGAAGTGGGGGC<br>TCAAGGCGGAGGAGGATCAGGTGGAGGAGGCTCCATCCAAGGATTGCTGGCCGAGGTGGGAGCTCAA<br>ATTCAGGGCCTGCTTGCTGAGGTCGGGGCACAGATCCAAGTACTCCTTGCAGAGGTAGGAGCTCAGG<br>GAGGAGGGGGATCTGGAGGAGGCGGCAGTACAATGACCGAAAAGGAGATAGTGGATTATGTGGCATC<br>CCAAGTGACAACAGCTAAAAAGCTGCGAGGCGGCGTAGTGTTCGTTGACGAGGTCCCAAAGGGACTT<br>ACCGGCAAGCTGGACGCTCGCAAAATTCGCGAGATACTTATAAAGGCAAAAAAGGGCGGCAAGATTG<br>CGGTTAATTCCGGTACCAAGCACCACCATCACCATCACTAA |
| 1-2.6 | KGSGGGSGGGSIQGLLAF<br>VGAQIQGLLAEVGAQIQG<br>LLAMVGAQGGGGSGGGGS<br>IQGLLAFVGAQIQGLLAE<br>VGAQIQGLLAMVGAQ | PIQPPDSRAAATMIQGLLAFVG<br>AQIQGLLAEVGAQIQGLLAMVG<br>AQGGGGSGGGGSIQGLLAFVGA<br>QIQGLLAEVGAQIQGLLAMVGA<br>QGGGGSGGGGSTMTEKEIVDYV<br>ASQVTTAKKLRGGVVFVDEVPK<br>GLTGKLDARKIREILIKAKKGG<br>KIAVNSGTKHHHHHH | AAAGGCTCTGGAGGGGGATCTGGAGGGGGATCCATC<br>CAAGGACTGCTGGCTTTTGTGGGCGCACAAATTCAG<br>GGATTGCTGGCCGAAGTGGGCGCCCAGATCCAAGGA<br>CTGCTTGCCATGGTAGGCGCACAGGGAGGGGGAGGA<br>AGCGGAGGTGGTGGGTCTATCCAGGGGCTCCTGGCA<br>TTTGTTGGCGCTCAGATACAGGGTCTGCTTGCAGAG<br>GTTGGGGCGCAAATACAGGGCCTCCTGGCTATGGTG<br>GGAGCTCAGTAATAAGGTACCAAGCACCACCATCAC<br>CATCACTAATGAGAT | CCGATCCAGCCTCCGGACTCTAGAGCGGCCGCCACCATGATTCAGGGCTTGCTGGCTTTTGTAGGGG<br>CCCAGATCCAGGGCTTGCTGGCCGAGGTCGGCGCACAGATTCAGGGCCTTCTTGCGATGGTAGGGGC<br>ACAGGGAGGTGGGGGCTCTGGCGGAGGAGGCTCAATTCAAGGGCTCCTGGCCTTCGTTGGAGCTCAA<br>ATCCAGGGACTTCTGGCTGAAGTGGGAGCTCAGATTCAGGGCCTGTTGGCAATGGTGGGAGCTCAGG<br>GCGGCGGAGGCTCCGGTGGCGGCGGCAGCACTATGACTGAGAAGGAGATTGTTGACTATGTGGCATC<br>TCAAGTCACGACCGCTAAGAAGTTGAGAGGTGGGGTGGTCTTTGTCGACGAGGTGCCTAAGGGACTG<br>ACCGGAAAACTGGATGCTAGGAAGATTAGAGAGATTTTGATTAAGGCTAAAAAGGGGGGAAAGATCG<br>CCGTAAATTCCGGTACCAAGCACCACCATCACCATCACTAA |

| | | | | |
|---|---|---|---|---|
| 1-<br>2.7 | KGSGGGSGGGSIQHLLAE<br>VGAQIQGLLAEVGAQIQG<br>LLALVGAQGGGGSGGGGS<br>IQHLLAEVGAQIQGLLAE<br>VGAQIQGLLALVGAQ | PIQPPDSRAAATMIQHLLAEVG<br>AQIQGLLAEVGAQIQGLLALVG<br>AQGGGGSGGGGSIQHLLAEVGA<br>QIQGLLAEVGAQIQGLLALVGA<br>QGGGGSGGGGSTMTEKEIVDYV<br>ASQVTTAKKLRGGVVFVDEVPK<br>GLTGKLDARKIREILIKAKKGG<br>KIAVNSGTKHHHHHH | AAAGGCTCTGGAGGGGGATCTGGAGGGGGATCCATT<br>CAGCACTTGTTGGCAGAAGTGGGAGCTCAGATCCAA<br>GGACTCCTGGCCGAGGTCGGCGCCCAGATCCAGGGA<br>CTTCTGGCCCTGGTCGGTGCTCAAGGAGGAGGCGGT<br>TCAGGGGGAGGCGGGAGCATCCAGCATTTGCTTGCT<br>GAAGTTGGAGCACAGATCCAGGGGCTGCTCGCAGAG<br>GTGGGCGCTCAAATCCAAGGTCTGTTGGCACTCGTG<br>GGTGCCCAGTAATAAGGTACCAAGCACCACCATCAC<br>CATCACTAATGAGAT | CCGATCCAGCCTCCGGACTCTAGAGCGGCCGCCACCATGATTCAGCACCTGCTTGCCGAAGTCGGGG<br>CACAGATTCAAGGGCTTCTCGCCGAAGTCGGAGCGCAGATTCAAGGTCTTCTGGCCCTGGTCGGTGC<br>TCAGGGCGGTGGGGGCTCAGGCGGAGGAGGGAGCATCCAGCATCTCCTCGCCGAAGTAGGTGCCCAG<br>ATACAGGGCTTGCTGGCGGAGGTTGGAGCCCAGATTCAGGGACTTCTGGCCCTCGTCGGCGCGCAGG<br>GCGGAGGAGGTAGCGGAGGTGGAGGGAGCACAATGACAGAGAAGGAGATCGTGGATTACGTAGCCTC<br>ACAAGTGACTACCGCTAAGAAGCTGAGAGGGGGAGTCGTGTTCGTTGATGAAGTACCTAAGGGATTG<br>ACCGGAAAACTTGACGCTCGAAAGATAAGAGAGATTCTGATCAAAGCAAAGAAGGGGGGGAAGATTG<br>CGGTTAACAGCGGTACCAAGCACCACCATCACCATCACTAA |
| 1-<br>7.1 | KGSGGGSGGGSINEMLEK<br>VAIEINEMLEKVADEINE<br>MLEKVADEGGGGSGGGGS<br>INEMLEKVAIEINEMLEK<br>VADEINEMLEKVADE | PIQPPDSRAAATMINEMLEKVA<br>IEINEMLEKVADEINEMLEKVA<br>DEGGGGSGGGGSINEMLEKVAI<br>EINEMLEKVADEINEMLEKVAD<br>EGGGGSGGGGSTMTEKEIVDYV<br>ASQVTTAKKLRGGVVFVDEVPK<br>GLTGKLDARKIREILIKAKKGG<br>KIAVNSGTKHHHHHH | AAAGGCTCTGGAGGGGGATCTGGAGGGGGATCCATC<br>AACGAAATGCTGGAAAAGGTTGCCATTGAAATAAAC<br>GAAATGCTGGAGAAAGTCGCCGATGAAATCAATGAA<br>ATGCTTGAAAAGGTCGCAGATGAGGGCGGGGGAGGA<br>AGTGGAGGCGGAGGGAGTATCAATGAAATGCTGGAA<br>AAAGTGGCTATAGAAATTAACGAGATGCTTGAGAAA<br>GTGGCAGATGAGATCAACGAAATGTTGGAAAAGGTA<br>GCCGATGAGTAATAAGGTACCAAGCACCACCATCAC<br>CATCACTAATGAGAT | CCGATCCAGCCTCCGGACTCTAGAGCGGCCGCCACCATGATCAACGAAATGCTCGAAAAAGTGGCGA<br>TCGAAATCAATGAAATGCTTGAGAAGGTCGCTGACGAGATCAACGAAATGTTGGAAAAGGTAGCGGA<br>TGAGGGAGGTGGGGGCAGCGGCGGCGGCGGGAGTATCAACGAGATGCTGGAGAAAGTCGCCATCGAG<br>ATCAATGAAATGCTTGAAAAGGTGGCAGACGAAATCAATGAGATGCTGGAAAAAGTGGCCGACGAGG<br>GCGGCGGTGGCTCCGGAGGAGGCGGAAGTACTATGACCGAGAAAGAAATCGTGGACTATGTAGCTTC<br>TCAGGTCACCACGGCCAAGAAATTGAGAGGGGGGGTCGTCTTTGTGGATGAAGTGCCGAAAGGGCTG<br>ACCGGCAAACTCGACGCCCGGAAAATTCGGGAGATTCTGATTAAGGCCAAAAAGGGGGGAAAGATCG<br>CCGTGAATAGCGGTACCAAGCACCACCATCACCATCACTAA |
| 1-<br>7.2 | KGSGGGSGGGSINEMLEK<br>VADEINEMLEKVAIEINE<br>MLEKVADEGGGGSGGGGS<br>INEMLEKVADEINEMLEK<br>VAIEINEMLEKVADE | PIQPPDSRAAATMINEMLEKVA<br>DEINEMLEKVAIEINEMLEKVA<br>DEGGGGSGGGGSINEMLEKVAD<br>EINEMLEKVAIEINEMLEKVAD<br>EGGGGSGGGGSTMTEKEIVDYV<br>ASQVTTAKKLRGGVVFVDEVPK<br>GLTGKLDARKIREILIKAKKGG<br>KIAVNSGTKHHHHHH | AAAGGCTCTGGAGGGGGATCTGGAGGGGGATCCATT<br>AACGAGATGCTCGAAAAGGTCGCAGATGAGATTAAC<br>GAAATGTTGGAAAAAGTGGCCATCGAAATTAACGAA<br>ATGCTGGAGAAAGTAGCCGACGAGGGAGGTGGCGGC<br>TCCGGTGGGGGAGGCAGCATCAATGAGATGCTGGAG<br>AAAGTGGCCGACGAAATCAATGAGATGCTCGAGAAA<br>GTGGCTATAGAGATCAACGAAATGCTTGAGAAGGTC<br>GCTGATGAGTAATAAGGTACCAAGCACCACCATCAC<br>CATCACTAATGAGAT | CCGATCCAGCCTCCGGACTCTAGAGCGGCCGCCACCATGATTAATGAGATGCTCGAAAAAGTCGCAG<br>ATGAGATTAACGAGATGTTGGAAAAAGTCGCCATCGAAATCAATGAGATGCTCGAAAAAGTGGCTGA<br>TGAAGGAGGAGGTGGCAGTGGTGGTGGCGGTTCCATAAATGAGATGCTCGAGAAAGTGGCAGACGAG<br>ATCAATGAGATGCTGGAGAAGGTGGCTATTGAGATTAACGAAATGCTCGAGAAAGTTGCAGACGAGG<br>GAGGAGGTGGCAGTGGCGGCGGCGGGTCTACAATGACCGAAAAGGAAATTGTTGACTATGTGGCTTC<br>TCAGGTGACTACAGCGAAGAAGCTGCGCGGAGGTGTGGTGTTTGTGGACGAGGTGCCCAAAGGGCTG<br>ACCGGCAAGCTGGATGCCAGAAAGATCAGAGAAATACTCATAAAGGCTAAGAAGGGAGGAAAAATTG<br>CCGTGAACTCCGGTACCAAGCACCACCATCACCATCACTAA |
| 1-<br>7.3 | KGSGGGSGGGSINEMLEK<br>VADEINEMLEKVADEINE<br>MLEKVADAGGGGSGGGGS<br>INEMLEKVADEINEMLEK<br>VADEINEMLEKVADA | PIQPPDSRAAATMINEMLEKVA<br>DEINEMLEKVADEINEMLEKVA<br>DAGGGGSGGGGSINEMLEKVAD<br>EINEMLEKVADEINEMLEKVAD<br>AGGGGSGGGGSTMTEKEIVDYV<br>ASQVTTAKKLRGGVVFVDEVPK<br>GLTGKLDARKIREILIKAKKGG<br>KIAVNSGTKHHHHHH | AAAGGCTCTGGAGGGGGATCTGGAGGGGGATCCATC<br>AACGAGATGCTCGAAAAGGTGGCGGACGAGATTAAT<br>GAGATGCTGGAGAAAGTGGCTGACGAAATTAATGAG<br>ATGCTCGAGAAGGTCGCTGATGCCGGGGGAGGGGGC<br>TCCGGAGGCGGAGGTTCTATTAACGAGATGCTGGAA<br>AAGGTAGCCGACGAGATCAATGAGATGCTTGAAAAA<br>GTCGCAGACGAAATTAACGAAATGCTGGAAAAAGTA<br>GCCGACGCATAATAAGGTACCAAGCACCACCATCAC<br>CATCACTAATGAGAT | CCGATCCAGCCTCCGGACTCTAGAGCGGCCGCCACCATGATCAACGAGATGCTCGAGAAGGTGGCCG<br>ATGAGATCAACGAGATGTTGGAGAAAGTGGCTGACGAGATCAACGAGATGTTGGAAAAGGTCGCTGA<br>TGCCGGTGGAGGCGGCAGCGGCGGCGGAGGATCTATCAATGAAATGCTGGAGAAGGTCGCTGATGAA<br>ATAAACGAAATGCTGGAGAAGGTGGCCGATGAGATCAATGAGATGTTGGAGAAAGTAGCGGATGCTG<br>GTGGGGGTGGCTCTGGAGGTGGTGGTTCAACAATGACCGAGAAAGAAATCGTCGACTACGTTGCGAG<br>CCAAGTCACCACAGCTAAAAAACTTCGAGGTGGGGTGGTCTTCGTTGACGAAGTCCCAAAGGGACTT<br>ACAGGTAAGCTGGACGCTCGCAAAATCCGGGAGATTCTGATCAAGGCGAAGAAAGGGGGCAAGATTG<br>CCGTCAATTCTGGTACCAAGCACCACCATCACCATCACTAA |
| 1-<br>7.4 | KGSGGGSGGGSIAEMLEK<br>VADEINEMLEKVADEINE<br>MLEKVADEGGGGSGGGGS<br>IAEMLEKVADEINEMLEK<br>VADEINEMLEKVADE | PIQPPDSRAAATMIAEMLEKVA<br>DEINEMLEKVADEINEMLEKVA<br>DEGGGGSGGGGSIAEMLEKVAD<br>EINEMLEKVADEINEMLEKVAD<br>EGGGGSGGGGSTMTEKEIVDYV<br>ASQVTTAKKLRGGVVFVDEVPK<br>GLTGKLDARKIREILIKAKKGG<br>KIAVNSGTKHHHHHH | AAAGGCTCTGGAGGGGGATCTGGAGGGGGATCCATC<br>GCTGAGATGTTGGAGAAGGTGGCTGATGAGATTAAC<br>GAAATGCTCGAAAAGGTTGCTGATGAGATTAATGAG<br>ATGCTTGAGAAGGTGGCGGATGAGGGAGGGGGTGGC<br>AGCGGTGGAGGAGGAAGCATAGCAGAGATGCTCGAG<br>AAAGTCGCGGACGAGATCAATGAAATGCTGGAAAAA<br>GTGGCCGACGAGATCAACGAAATGCTTGAGAAAGTG<br>GCCGATGAGTAATAAGGTACCAAGCACCACCATCAC<br>CATCACTAATGAGAT | CCGATCCAGCCTCCGGACTCTAGAGCGGCCGCCACCATGATCGCCGAGATGTTGGAGAAAGTGGCAG<br>ACGAGATTAACGAGATGCTGGAGAAAGTGGCCGACGAGATAAATGAGATGCTGGAGAAGGTGGCTGA<br>TGAGGGCGGCGGCGGCAGCGGCGGAGGTGGCTCAATTGCAGAGATGCTGGAAAAAGTTGCCGACGAG<br>ATAAACGAGATGCTCGAGAAAGTTGCTGACGAAATAAACGAAATGCTTGAGAAAGTTGCTGACGAGG<br>GCGGCGGCGGATCAGGGGGAGGAGGATCTACAATGACTGAGAAGGAAATTGTTGACTATGTCGCGAG<br>TCAGGTCACAACTGCCAAGAAGCTGCGCGGAGGCGTGGTCTTCGTGGACGAAGTACCTAAGGGACTT<br>ACAGGTAAGCTCGATGCCAGAAAGATACGGGAGATCCTCATTAAGGCCAAGAAGGGAGGAAAAATCG<br>CTGTGAATAGCGGTACCAAGCACCACCATCACCATCACTAA |

| | | | | |
|---|---|---|---|---|
| 1-7.5 | KGSGGGSGGGSIREMLEK<br>VAIEINEMLEKVADEINE<br>MLEKVADEGGGGSGGGGS<br>IREMLEKVAIEINEMLEK<br>VADEINEMLEKVADE | PIQPPDSRAAATMIREMLEKVA<br>IEINEMLEKVADEINEMLEKVA<br>DEGGGGSGGGGSIREMLEKVAI<br>EINEMLEKVADEINEMLEKVAD<br>EGGGGSGGGGSTMTEKEIVDYV<br>ASQVTTAKKLRGGVVFVDEVPK<br>GLTGKLDARKIREILIKAKKGG<br>KIAVNSGTKHHHHHH | AAAGGCTCTGGAGGGGGATCTGGAGGGGGATCCATT<br>CGGGAGATGCTCGAAAAAGTCGCTATCGAAATTAAT<br>GAAATGCTCGAGAAAGTGGCTGACGAGATAAATGAG<br>ATGCTCGAGAAAGTGGCCGATGAGGGAGGAGGGGGA<br>AGCGGAGGAGGTGGATCAATCAGAGAGATGCTGGAG<br>AAGGTTGCCATTGAGATCAACGAAATGCTCGAGAAG<br>GTCGCAGACGAAATAAATGAGATGTTGGAAAAGGTC<br>GCGGATGAATAATAAGGTACCAAGCACCACCATCAC<br>CATCACTAATGAGAT | CCGATCCAGCCTCCGGACTCTAGAGCGGCCGCCACCATGATCAGAGAGATGCTGGAGAAGGTGGCCA<br>TAGAAATCAATGAGATGCTGGAAAAGGTTGCGGATGAAATTAACGAAATGCTTGAAAAGGTCGCCGA<br>CGAGGGCGGAGGAGGAAGTGGCGGGGGCGGCAGTATAAGAGAGATGCTGGAAAAAGTCGCCATCGAG<br>ATTAACGAGATGTTGGAAAAAGTCGCAGATGAAATCAATGAAATGCTTGAAAAAGTGGCCGATGAGG<br>GTGGTGGCGGGAGCGGGGGCGGCGGCAGCACAATGACAGAGAAGGAAATCGTGGACTATGTGGCCAG<br>CCAAGTGACAACAGCCAAGAAACTGCGAGGAGGCGTCGTCTTTGTGGATGAGGTGCCAAAAGGGCTT<br>ACAGGAAAGTTGGACGCACGCAAGATCAGAGAAATTCTCATCAAAGCCAAGAAGGGAGGTAAAATAG<br>CCGTTAATAGCGGTACCAAGCACCACCATCACCATCACTAA |
| 1-7.6 | KGSGGGSGGGSIREMLEK<br>VADEIKEMLEKVADEINE<br>MLEKVADEGGGGSGGGGS<br>IREMLEKVADEIKEMLEK<br>VADEINEMLEKVADE | PIQPPDSRAAATMIREMLEKVA<br>DEIKEMLEKVADEINEMLEKVA<br>DEGGGGSGGGGSIREMLEKVAD<br>EIKEMLEKVADEINEMLEKVAD<br>EGGGGSGGGGSTMTEKEIVDYV<br>ASQVTTAKKLRGGVVFVDEVPK<br>GLTGKLDARKIREILIKAKKGG<br>KIAVNSGTKHHHHHH | AAAGGCTCTGGAGGGGGATCTGGAGGGGGATCCATT<br>CGCGAAATGCTTGAAAAGGTGGCCGACGAGATCAAG<br>GAGATGCTGGAAAAAGTGGCCGATGAGATAAACGAA<br>ATGTTGGAGAAAGTGGCCGATGAAGGGGGCGGAGGG<br>TCTGGAGGCGGGGGCAGCATACGCGAGATGCTTGAA<br>AAGGTGGCAGATGAAATAAAAGAAATGCTGGAGAAA<br>GTGGCGGACGAAATTAATGAAATGTTGGAGAAGGTT<br>GCCGATGAGTAATAAGGTACCAAGCACCACCATCAC<br>CATCACTAATGAGAT | CCGATCCAGCCTCCGGACTCTAGAGCGGCCGCCACCATGATTCGCGAAATGCTTGAAAAAGTCGCCG<br>ACGAGATCAAGGAGATGTTGGAGAAGGTGGCCGATGAAATAAACGAGATGCTGGAAAAAGTGGCCGA<br>CGAGGGTGGGGGCGGCTCAGGTGGCGGTGGAAGCATTCGGGAGATGTTGGAAAAGGTTGCCGATGAA<br>ATAAAAGAGATGTTGGAGAAAGTGGCCGACGAAATCAACGAAATGTTGGAGAAAGTGGCCGATGAAG<br>GGGGAGGCGGCAGCGGCGGAGGTGGCTCAACCATGACAGAGAAGGAAATTGTTGATTATGTGGCCTC<br>TCAGGTTACCACCGCAAAGAAGCTCAGAGGCGGTGTCGTTTTTGTGGATGAAGTGCCCAAGGGGCTC<br>ACAGGCAAGCTCGACGCCAGGAAAATTAGGGAGATTCTCATCAAAGCCAAAAAGGGAGGGAAAATCG<br>CTGTTAACTCAGGTACCAAGCACCACCATCACCATCACTAA |
| 1-7.7 | KGSGGGSGGGSIREMLEK<br>VADEINEMLEKVADEIKE<br>MLEKVADEGGGGSGGGGS<br>IREMLEKVADEINEMLEK<br>VADEIKEMLEKVADE | PIQPPDSRAAATMIREMLEKVA<br>DEINEMLEKVADEIKEMLEKVA<br>DEGGGGSGGGGSIREMLEKVAD<br>EINEMLEKVADEIKEMLEKVAD<br>EGGGGSGGGGSTMTEKEIVDYV<br>ASQVTTAKKLRGGVVFVDEVPK<br>GLTGKLDARKIREILIKAKKGG<br>KIAVNSGTKHHHHHH | AAAGGCTCTGGAGGGGGATCTGGAGGGGGATCCATT<br>CGCGAGATGCTGGAAAAGGTGGCCGATGAGATCAAC<br>GAGATGCTTGAAAAGGTCGCCGACGAAATTAAGGAA<br>ATGCTGGAGAAAGTGGCTGATGAGGGTGGCGGCGGT<br>TCTGGTGGTGGGGGGTCTATTAGAGAAATGCTGGAA<br>AAGGTGGCAGACGAAATAAATGAAATGCTTGAAAAG<br>GTCGCAGACGAGATAAAGGAGATGCTTGAGAAGGTC<br>GCTGACGAGTAATAAGGTACCAAGCACCACCATCAC<br>CATCACTAATGAGAT | CCGATCCAGCCTCCGGACTCTAGAGCGGCCGCCACCATGATCAGGGAAATGCTGGAGAAAGTAGCTG<br>ATGAGATTAACGAGATGCTCGAAAAGGTCGCTGACGAAATTAAGGAGATGCTGGAGAAGGTTGCAGA<br>TGAAGGAGGAGGGGGCAGCGGAGGCGGGGGCTCTATACGAGAGATGCTCGAGAAGGTTGCAGATGAG<br>ATAAACGAAATGCTGGAGAAGGTGGCTGATGAAATAAAAGAAATGCTGGAAAAAGTGGCAGACGAGG<br>GGGGAGGGGGATCAGGAGGTGGCGGCTCAACGATGACCGAGAAAGAAATCGTGGACTATGTTGCCAG<br>TCAGGTCACGACCGCCAAAAAACTTAGGGGGGGCGTGGTGTTCGTTGATGAAGTGCCAAAAGGCCTG<br>ACAGGTAAGCTCGACGCCCGGAAGATTCGGGAGATACTTATCAAAGCCAAGAAAGGAGGGAAGATAG<br>CTGTCAACTCCGGTACCAAGCACCACCATCACCATCACTAA |
| 1-7.8 | KGSGGGSGGGSIREMLEK<br>VADEINEMLEKVADEIRE<br>MLEKVADEGGGGSGGGGS<br>IREMLEKVADEINEMLEK<br>VADEIREMLEKVADE | PIQPPDSRAAATMIREMLEKVA<br>DEINEMLEKVADEIREMLEKVA<br>DEGGGGSGGGGSIREMLEKVAD<br>EINEMLEKVADEIREMLEKVAD<br>EGGGGSGGGGSTMTEKEIVDYV<br>ASQVTTAKKLRGGVVFVDEVPK<br>GLTGKLDARKIREILIKAKKGG<br>KIAVNSGTKHHHHHH | AAAGGCTCTGGAGGGGGATCTGGAGGGGGATCCATC<br>CGGGAGATGCTGGAGAAAGTGGCAGACGAAATTAAT<br>GAAATGCTCGAGAAGGTCGCCGACGAGATCCGCGAG<br>ATGCTTGAGAAAGTTGCCGATGAAGGCGGGGGCGGA<br>AGTGGAGGCGGAGGATCTATACGCGAAATGCTGGAA<br>AAGGTAGCTGATGAGATCAACGAAATGCTTGAGAAA<br>GTGGCCGACGAGATTAGGGAAATGCTGGAGAAAGTC<br>GCAGATGAATAATAAGGTACCAAGCACCACCATCAC<br>CATCACTAATGAGAT | CCGATCCAGCCTCCGGACTCTAGAGCGGCCGCCACCATGATCAGAGAAATGCTCGAGAAAGTTGCCG<br>ATGAAATCAACGAGATGCTCGAGAAAGTCGCTGACGAGATACGGGAGATGCTCGAGAAAGTCGCCGA<br>TGAAGGTGGCGGCGGTTCTGGCGGAGGTGGCTCTATTCGCGAGATGTTGGAAAAGGTGGCAGACGAA<br>ATTAATGAGATGCTCGAAAAGGTCGCGGACGAAATCCGGGAGATGCTTGAGAAAGTCGCCGACGAGG<br>GCGGAGGGGGCAGTGGGGGAGGAGGCAGCACCATGACCGAAAAGGAGATCGTGGACTACGTGGCGAG<br>TCAGGTGACAACAGCCAAGAAGCTGAGGGGAGGAGTCGTCTTTGTGGATGAGGTGCCAAAGGGACTT<br>ACTGGCAAACTCGATGCAAGGAAAATCAGGGAGATTCTGATTAAGGCCAAGAAGGGGGGCAAGATCG<br>CTGTGAACAGTGGTACCAAGCACCACCATCACCATCACTAA |
| 2-1 | EHGKGSGGGSGGGSAKAE<br>EAAQALLAELTAAAKKAV<br>EALLKRLKGGGGSGGGGS<br>AKAEEAAQALLAELTAAA<br>KKAVEALLKRLK | PIQPPDSRAAATMAKAEEAAQA<br>LLAELTAAAKKAVEALLKRLKG<br>GGGSGGGGSAKAEEAAQALLAE<br>LTAAAKKAVEALLKRLKGGGGS<br>GGGGSTMTEKEIVDYVASQVTT<br>AKKLRGGVVFVDEVPKGLTGKL<br>DARKIREILIKAKKGGKIAVNS<br>GTKHHHHHH | CGAGCATGGGAAAGGCTCTGGAGGGGGATCTGGAGG<br>GGGATCCGCTAAGGCTGAAGAGGCCGCACAGGCTCT<br>GCTCGCAGAGCTTACCGCAGCGGCCAAGAAAGCTGT<br>GGAGGCCCTGCTGAAGAGGTTGAAGGGCGGTGGCGG<br>AAGCGGTGGAGGTGGGTCCGCTAAAGCCGAGGAGGC<br>AGCCCAAGCTCTCCTTGCCGAGCTCACAGCAGCAGC<br>CAAGAAGGCAGTTGAGGCACTGCTGAAGCGCCTCAA<br>GTAATAAGGTACCAAGCACCACCATCACCATCACTA<br>ATGAGATATCTC | CCGATCCAGCCTCCGGACTCTAGAGCGGCCGCCACCATGGCCAAGGCAGAGGAGGCTGCCCAAGCAC<br>TGTTGGCCGAGCTGACTGCTGCCGCCAAGAAAGCCGTTGAGGCTTTGCTCAAACGCCTTAAGGGCGG<br>AGGCGGCAGCGGCGGAGGGGGCAGTGCGAAAGCCGAGGAAGCCGCGCAAGCTCTGCTGGCCGAGCTG<br>ACCGCTGCCGCCAAAAAGGCTGTCGAGGCTCTGTTGAAACGCCTGAAGGGCGGCGGCGGCTCTGGTG<br>GGGGAGGAAGTACCATGACCGAAAAAGAGATCGTCGATTACGTTGCCTCCCAGGTTACTACAGCTAA<br>GAAGCTTCGAGGGGGTGTGGTATTTGTAGACGAGGTGCCTAAAGGCCTGACCGGTAAGCTGGATGCC<br>CGAAAGATTAGAGAGATTCTGATTAAGGCCAAGAAGGGCGGAAAAATTGCCGTTAACTCCGGTACCA<br>AGCACCACCATCACCATCACTAA |

| | | | | |
|---|---|---|---|---|
| 2-2 | EHGKGSGGGSGGGSGKAE<br>KAVEKLLKELKEAAEAAA<br>KKLLKELKGGGGSGGGGS<br>GKAEKAVEKLLKELKEAA<br>EAAAKKLLKELK | PIQPPDSRAAATMGKAEKAVEK<br>LLKELKEAAEAAAKKLLKELKG<br>GGGSGGGGSGKAEKAVEKLLKE<br>LKEAAEAAAKKLLKELKGGGGS<br>GGGGSTMTEKEIVDYVASQVTT<br>AKKLRGGVVFVDEVPKGLTGKL<br>DARKIREILIKAKKGGKIAVNS<br>GTKHHHHHH | CGAGCATGGGAAAGGCTCTGGAGGGGGATCTGGAGG<br>GGGATCCGGCAAGGCGGAGAAGGCTGTTGAGAAACT<br>GCTCAAAGAACTGAAAGAAGCCGCAGAAGCCGCAGC<br>TAAGAAGCTTTTGAAAGAGCTGAAAGGAGGTGGAGG<br>AAGTGGTGGGGGCGGCTCTGGCAAAGCCGAGAAGGC<br>AGTGGAGAAACTGCTGAAAGAACTGAAAGAGGCAGC<br>CGAAGCAGCCGCCAAAAAACTCCTCAAGGAGCTGAA<br>ATAATAAGGTACCAAGCACCACCATCACCATCACTA<br>ATGAGATATCTC | CCGATCCAGCCTCCGGACTCTAGAGCGGCCGCCACCATGGGAAAAGCTGAGAAGGCAGTGGAGAAAC<br>TGCTGAAGGAATTGAAAGAGGCCGCCGAGGCTGCCGCTAAAAAGCTCCTCAAAGAACTGAAGGGCGG<br>AGGGGGGTCAGGTGGGGGAGGGAGTGGAAAGGCAGAAAAGGCGGTGGAAAAGCTCCTGAAAGAGCTG<br>AAAGAGGCTGCGGAGGCAGCCGCAAAGAAACTCTTGAAGGAACTTAAAGGGGGAGGCGGCTCAGGGG<br>GAGGGGGCAGCACAATGACTGAAAAAGAAATCGTTGATTACGTCGCCAGCCAGGTGACAACCGCAAA<br>GAAGCTGCGGGGCGGAGTCGTGTTCGTGGACGAGGTGCCTAAGGGGCTTACCGGGAAGCTGGACGCC<br>AGAAAAATCCGGGAGATACTTATCAAAGCAAAGAAGGGTGGCAAGATCGCTGTGAACTCCGGTACCA<br>AGCACCACCATCACCATCACTAA |
| 2-3 | EHGKGSGGGSGGGSDKVK<br>KVAEKALKELTAGAKKVV<br>EKMEKELKGGGGSGGGGS<br>DKVKKVAEKALKELTAGA<br>KKVVEKMEKELK | PIQPPDSRAAATMDKVKKVAEK<br>ALKELTAGAKKVVEKMEKELKG<br>GGGSGGGGSDKVKKVAEKALKE<br>LTAGAKKVVEKMEKELKGGGGS<br>GGGGSTMTEKEIVDYVASQVTT<br>AKKLRGGVVFVDEVPKGLTGKL<br>DARKIREILIKAKKGGKIAVNS<br>GTKHHHHHH | CGAGCATGGGAAAGGCTCTGGAGGGGGATCTGGAGG<br>GGGATCCGACAAGGTCAAGAAAGTAGCTGAGAAAGC<br>TCTCAAAGAACTGACAGCCGGCGCGAAGAAAGTGGT<br>TGAAAAGATGGAAAAAGAGTTGAAAGGCGGGGGTGG<br>ATCTGGGGGCGGAGGGAGCGATAAGGTCAAGAAGGT<br>AGCCGAAAAGGCGTTGAAAGAACTGACTGCCGGGGC<br>TAAGAAAGTGGTAGAAAAAATGGAAAAAGAGTTGAA<br>GTAATAAGGTACCAAGCACCACCATCACCATCACTA<br>ATGAGATATCTC | CCGATCCAGCCTCCGGACTCTAGAGCGGCCGCCACCATGGACAAGGTAAAGAAGGTGGCTGAAAAAG<br>CACTTAAAGAACTGACAGCCGGCGCAAAGAAGGTGGTGGAGAAGATGGAGAAAGAGTTGAAGGGCGG<br>AGGGGGATCTGGAGGCGGAGGATCAGATAAGGTTAAGAAAGTGGCAGAGAAGGCGCTCAAAGAGCTT<br>ACTGCAGGGGCCAAGAAGGTGGTAGAGAAGATGGAGAAGGAGTTGAAAGGAGGTGGAGGGAGTGGGG<br>GCGGAGGGTCTACAATGACTGAAAAGGAGATAGTTGACTATGTTGCCTCTCAGGTGACAACTGCCAA<br>AAAACTCCGGGGCGGCGTTGTTTTCGTTGACGAGGTCCCAAAAGGGCTGACTGGAAAGCTGGACGCC<br>CGGAAAATACGGGAAATTCTTATCAAAGCCAAGAAAGGTGGCAAGATAGCTGTGAATAGTGGTACCA<br>AGCACCACCATCACCATCACTAA |
| 2-4 | EHGKGSGGGSGGGSDKAK<br>AAAAALEAKLTAAAKAAT<br>AELLAALQGGGGSGGGGS<br>DKAKAAAAALEAKLTAAA<br>KAATAELLAALQ | PIQPPDSRAAATMDKAKAAAAA<br>LEAKLTAAAKAATAELLAALQG<br>GGGSGGGGSDKAKAAAAALEAK<br>LTAAAKAATAELLAALQGGGGS<br>GGGGSTMTEKEIVDYVASQVTT<br>AKKLRGGVVFVDEVPKGLTGKL<br>DARKIREILIKAKKGGKIAVNS<br>GTKHHHHHH | CGAGCATGGGAAAGGCTCTGGAGGGGGATCTGGAGG<br>GGGATCCGACAAGGCCAAAGCTGCCGCCGCGGCACT<br>TGAAGCCAAATTGACCGCTGCTGCCAAAGCAGCCAC<br>TGCTGAGCTGCTCGCAGCATTGCAGGGCGGCGGCGG<br>ATCTGGGGGTGGCGGGAGCGATAAAGCCAAGGCAGC<br>CGCAGCCGCATTGGAGGCCAAGCTTACTGCTGCAGC<br>CAAAGCCGCTACAGCCGAACTTCTTGCCGCCCTGCA<br>GTAATAAGGTACCAAGCACCACCATCACCATCACTA<br>ATGAGATATCTC | CCGATCCAGCCTCCGGACTCTAGAGCGGCCGCCACCATGGACAAAGCAAAGGCCGCTGCTGCCGCCC<br>TGGAGGCCAAGTTGACCGCAGCCGCCAAAGCCGCTACTGCAGAGCTCCTGGCTGCGCTGCAGGGAGG<br>GGGAGGTAGTGGTGGGGGTGGCAGCGATAAAGCTAAAGCAGCAGCCGCGGCTCTTGAAGCAAAGCTG<br>ACGGCCGCAGCCAAAGCCGCAACCGCCGAATTGCTCGCGGCCCTGCAAGGAGGCGGGGGATCTGGCG<br>GCGGCGGGAGTACTATGACCGAGAAAGAGATCGTAGATTACGTTGCCTCACAGGTCACTACAGCGAA<br>GAAACTGAGAGGCGGGGTGGTCTTTGTTGACGAAGTCCCAAAGGGACTGACAGGTAAACTGGATGCC<br>AGAAAGATCAGAGAGATTTTGATAAAAGCCAAAAAAGGGGGAAAGATTGCAGTGAACTCCGGTACCA<br>AGCACCACCATCACCATCACTAA |
| 2-5 | EHGKGSGGGSGGGSEAAE<br>AAVKAMEEALVKAVKAAA<br>EALLKALKGGGGSGGGGS<br>EAAEAAVKAMEEALVKAV<br>KAAAEALLKALK | PIQPPDSRAAATMEAAEAAVKA<br>MEEALVKAVKAAAEALLKALKG<br>GGGSGGGGSEAAEAAVKAMEEA<br>LVKAVKAAAEALLKALKGGGGS<br>GGGGSTMTEKEIVDYVASQVTT<br>AKKLRGGVVFVDEVPKGLTGKL<br>DARKIREILIKAKKGGKIAVNS<br>GTKHHHHHH | CGAGCATGGGAAAGGCTCTGGAGGGGGATCTGGAGG<br>GGGATCCGAAGCTGCGGAAGCCGCCGTAAAAGCGAT<br>GGAGGAGGCCCTTGTCAAAGCTGTCAAGGCTGCCGC<br>AGAAGCCCTGCTGAAGGCTCTTAAGGGCGGTGGAGG<br>ATCTGGAGGCGGAGGAAGTGAGGCAGCCGAGGCGGC<br>CGTGAAGGCCATGGAGGAGGCACTCGTTAAAGCAGT<br>TAAAGCCGCGGCTGAAGCACTGCTGAAAGCACTCAA<br>GTAATAAGGTACCAAGCACCACCATCACCATCACTA<br>ATGAGATATCTC | CCGATCCAGCCTCCGGACTCTAGAGCGGCCGCCACCATGGAAGCAGCTGAGGCGGCTGTGAAAGCCA<br>TGGAGGAGGCCTTGGTGAAAGCTGTCAAAGCCGCTGCCGAAGCTCTCCTTAAGGCACTGAAGGGCGG<br>CGGCGGCAGTGGAGGCGGCGGCTCAGAGGCAGCTGAGGCAGCCGTCAAGGCTATGGAGGAAGCACTG<br>GTCAAAGCAGTCAAAGCTGCCGCGGAAGCCCTTTTGAAAGCACTCAAGGGAGGCGGTGGGTCTGGAG<br>GGGGCGGCTCTACCATGACGGAGAAGGAGATCGTGGATTACGTTGCCTCCCAAGTTACTACCGCGAA<br>GAAACTTCGAGGAGGGGTCGTTTTCGTGGACGAGGTGCCCAAGGGTCTGACTGGTAAACTCGATGCT<br>AGGAAAATCCGCGAGATTCTGATCAAAGCCAAAAAAGGAGGGAAGATAGCCGTCAACAGCGGTACCA<br>AGCACCACCATCACCATCACTAA |
| 2-6 | EHGKGSGGGSGGGSEKAA<br>EAVKKAEKKLEEAVKKVA<br>EELKKELEGGGGSGGGGS<br>EKAAEAVKKAEKKLEEAV<br>KKVAEELKKELE | PIQPPDSRAAATMEKAAEAVKK<br>AEKKLEEAVKKVAEELKKELEG<br>GGGSGGGGSEKAAEAVKKAEKK<br>LEEAVKKVAEELKKELEGGGGS<br>GGGGSTMTEKEIVDYVASQVTT<br>AKKLRGGVVFVDEVPKGLTGKL<br>DARKIREILIKAKKGGKIAVNS<br>GTKHHHHHH | CGAGCATGGGAAAGGCTCTGGAGGGGGATCTGGAGG<br>GGGATCCGAAAAAGCAGCCGAAGCTGTGAAAAAAGC<br>CGAGAAAAAACTCGAAGAAGCCGTGAAGAAAGTGGC<br>AGAGGAGCTGAAGAAAGAGCTGGAAGGGGGAGGCGG<br>CTCTGGCGGTGGAGGTTCCGAGAAAGCGGCCGAGGC<br>CGTAAAGAAAGCAGAGAAGAAACTGGAGGAAGCCGT<br>GAAGAAGGTGGCAGAGGAGTTGAAGAAGGAGCTGGA<br>ATAATAAGGTACCAAGCACCACCATCACCATCACTA<br>ATGAGATATCTC | CCGATCCAGCCTCCGGACTCTAGAGCGGCCGCCACCATGGAGAAAGCAGCAGAGGCCGTGAAGAAGG<br>CCGAAAAGAAACTGGAGGAGGCCGTGAAAAAAGTCGCTGAGGAACTGAAGAAAGAGCTGGAAGGTGG<br>GGGAGGGTCTGGAGGCGGGGGGTCAGAGAAGGCCGCAGAGGCCGTTAAAAAAGCCGAGAAAAAACTC<br>GAGGAAGCCGTAAAAAAAGTGGCAGAAGAACTGAAGAAAGAGTTGGAAGGAGGAGGAGGCTCTGGAG<br>GAGGAGGATCAACCATGACCGAAAAAGAAATAGTGGATTACGTGGCCAGCCAGGTGACAACAGCTAA<br>GAAGTTGAGGGGCGGGGTCGTGTTCGTAGATGAGGTTCCTAAAGGTCTGACAGGAAAGCTGGACGCA<br>CGCAAAATTAGAGAGATCCTGATAAAAGCCAAGAAGGGCGGGAAGATCGCCGTCAACAGTGGTACCA<br>AGCACCACCATCACCATCACTAA |

| | | | | |
|---|---|---|---|---|
| 2-7 | EHGKGSGGGSGGGSGRVE<br>RVAEELLAKLKEAAEKAV<br>KEALEKLKGGGGSGGGGS<br>GRVERVAEELLAKLKEAA<br>EKAVKEALEKLK | PIQPPDSRAAATMGRVERVAEE<br>LLAKLKEAAEKAVKEALEKLKG<br>GGGSGGGGSGRVERVAEELLAK<br>LKEAAEKAVKEALEKLKGGGGS<br>GGGGSTMTEKEIVDYVASQVTT<br>AKKLRGGVVFVDEVPKGLTGKL<br>DARKIREILIKAKKGGKIAVNS<br>GTKHHHHHH | CGAGCATGGGAAAGGCTCTGGAGGGGGATCTGGAGG<br>GGGATCCGGAAGGGTCGAGAGAGTTGCAGAGGAATT<br>GCTGGCTAAGCTCAAAGAGGCTGCAGAAAAGGCAGT<br>CAAGGAAGCCCTGGAGAAGCTCAAGGGCGGTGGGGG<br>CTCCGGTGGTGGCGGCTCAGGGCGGGTAGAGCGCGT<br>CGCAGAGGAGTTGCTGGCAAAACTCAAGGAAGCAGC<br>AGAAAAAGCAGTCAAAGAGGCTCTGGAAAAGCTGAA<br>ATAATAAGGTACCAAGCACCACCATCACCATCACTA<br>ATGAGATATCTC | CCGATCCAGCCTCCGGACTCTAGAGCGGCCGCCACCATGGGTAGGGTTGAGCGGGTGGCTGAGGAGC<br>TCCTCGCGAAGCTGAAGGAGGCCGCTGAGAAAGCCGTGAAAGAAGCACTCGAGAAACTGAAGGGCGG<br>AGGAGGATCTGGCGGCGGCGGCTCCGGACGGGTAGAACGCGTCGCCGAAGAACTGCTGGCGAAACTT<br>AAAGAGGCAGCCGAAAAAGCCGTTAAGGAGGCTCTGGAAAAACTCAAGGGAGGGGGCGGAGCGGGG<br>GCGGCGGTAGCACCATGACCGAAAAGGAGATTGTTGACTATGTGGCCTCTCAAGTGACCACTGCTAA<br>GAAGTTGCGCGGCGGGGTTGTGTTTGTGGATGAGGTGCCAAAGGGGCTTACAGGAAAGCTGGATGCA<br>CGGAAGATCAGGGAGATTCTCATCAAGGCTAAAAAAGGAGGAAAGATTGCCGTGAATTCCGGTACCA<br>AGCACCACCATCACCATCACTAA |
| 2-8 | EHGKGSGGGSGGGSGKVE<br>EAVKELEKRLVEGVKKAA<br>EELLKRLKGGGGSGGGGS<br>GKVEEAVKELEKRLVEGV<br>KKAAEELLKRLK | PIQPPDSRAAATMGKVEEAVKE<br>LEKRLVEGVKKAAEELLKRLKG<br>GGGSGGGGSGKVEEAVKELEKR<br>LVEGVKKAAEELLKRLKGGGGS<br>GGGGSTMTEKEIVDYVASQVTT<br>AKKLRGGVVFVDEVPKGLTGKL<br>DARKIREILIKAKKGGKIAVNS<br>GTKHHHHHH | CGAGCATGGGAAAGGCTCTGGAGGGGGATCTGGAGG<br>GGGATCCGGCAAAGTGGAGGAGGCCGTTAAAGAGCT<br>TGAAAAGAGACTTGTGGAAGGGGTGAAAAAGGCAGC<br>GGAAGAACTGCTGAAACGGCTGAAGGGCGGCGGGGG<br>CAGTGGGGGAGGGGGCAGCGGGAAGGTAGAGGAAGC<br>AGTCAAAGAATTGGAGAAACGCCTGGTGGAGGGCGT<br>GAAAAAGGCGGCCGAGGAGCTGCTGAAGCGCCTGAA<br>GTAATAAGGTACCAAGCACCACCATCACCATCACTA<br>ATGAGATATCTC | CCGATCCAGCCTCCGGACTCTAGAGCGGCCGCCACCATGGGGAAAGTCGAGGAAGCGGTGAAAGAAC<br>TTGAAAAGCGGCTGGTTGAGGGAGTGAAAAAAGCCGCCGAAGAGCTGCTGAAGCGACTGAAAGGAGG<br>CGGGGGGTCCGGCGGCGGGGGATCTGGAAAAGTCGAGGAGGCAGTGAAGGAGCTGGAGAAGCGCCTG<br>GTCGAGGGGGTGAAAAAAGCCGCGGAAGAGCTGCTTAAACGCCTCAAGGGAGGGGGAGGATCTGGAG<br>GGGGCGGAAGCACAATGACTGAAAAGGAGATCGTGGATTACGTGGCCTCACAGGTCACCACAGCAAA<br>GAAACTGCGCGGCGGTGTCGTCTTCGTTGACGAAGTGCCGAAGGGACTTACAGGTAAGCTGGATGCT<br>CGCAAAATAAGGGAGATACTGATAAAGGCCAAGAAAGGAGGTAAAATCGCAGTGAACAGCGGTACCA<br>AGCACCACCATCACCATCACTAA |
| 2-9 | EHGKGSGGGSGGGSERLR<br>ALLEQLRAELEEELARLL<br>ARLEEELAGGGGSGGGGS<br>ERLRALLEQLRAELEEEL<br>ARLLARLEEELA | PIQPPDSRAAATMERLRALLEQ<br>LRAELEEELARLLARLEEELAG<br>GGGSGGGGSERLRALLEQLRAE<br>LEEELARLLARLEEELAGGGGS<br>GGGGSTMTEKEIVDYVASQVTT<br>AKKLRGGVVFVDEVPKGLTGKL<br>DARKIREILIKAKKGGKIAVNS<br>GTKHHHHHH | CGAGCATGGGAAAGGCTCTGGAGGGGGATCTGGAGG<br>GGGATCCGAAAGATTGCGAGCCCTGCTGGAGCAGCT<br>GCGGGCCGAGTTGGAAGAGGAACTGGCTCGACTGCT<br>GGCCAGACTTGAAGAAGAGCTGGCGGGAGGCGGGGG<br>CTCCGGTGGAGGTGGTAGTGAAAGGCTGAGGGCCCT<br>CCTGGAACAGCTGCGGGCTGAACTGGAAGAAGAGCT<br>GGCTAGGCTGCTTGCCAGACTCGAAGAAGAGCTGGC<br>ATAATAAGGTACCAAGCACCACCATCACCATCACTA<br>ATGAGATATCTC | CCGATCCAGCCTCCGGACTCTAGAGCGGCCGCCACCATGGAACGACTGAGGGCGCTGCTGGAACAGC<br>TGAGGGCGGAATTGGAGGAAGAGCTTGCCAGACTGTTGGCCAGACTTGAGGAGGAACTCGCTGGAGG<br>GGGTGGGAGCGGTGGAGGAGGGTCTGAAAGGCTGCGGGCACTCCTGGAGCAGCTGCGGGCCGAGCTG<br>GAAGAGGAGCTGGCAAGGCTGCTTGCAAGGCTGGAAGAGGAATTGGCCGGAGGAGGCGGGTCTGGGG<br>GAGGTGGGTCCACCATGACCGAAAAGGAAATTGTTGACTACGTGGCAAGTCAGGTGACAACTGCTAA<br>AAAGCTTCGGGGCGGTGTGGTTTTCGTCGACGAGGTGCCCAAGGGACTGACCGGAAAGCTGGATGCC<br>CGAAAAATTCGGGAGATTCTGATCAAGGCCAAAAAGGGCGGCAAAATCGCCGTAAATAGCGGTACCA<br>AGCACCACCATCACCATCACTAA |
| 2-<br>10 | EHGKGSGGGSGGGSGKVE<br>EAVKKAEKALVAAAKAVA<br>AKLLKELKGGGGSGGGGS<br>GKVEEAVKKAEKALVAAA<br>KAVAAKLLKELK | PIQPPDSRAAATMGKVEEAVKK<br>AEKALVAAAKAVAAKLLKELKG<br>GGGSGGGGSGKVEEAVKKAEKA<br>LVAAAKAVAAKLLKELKGGGGS<br>GGGGSTMTEKEIVDYVASQVTT<br>AKKLRGGVVFVDEVPKGLTGKL<br>DARKIREILIKAKKGGKIAVNS<br>GTKHHHHHH | CGAGCATGGGAAAGGCTCTGGAGGGGGATCTGGAGG<br>GGGATCCGGGAAAGTAGAAGAGGCAGTGAAAAAAGC<br>TGAGAAAGCACTCGTAGCCGCCGCAAAAGCCGTTGC<br>GGCCAAGTTGCTCAAGGAACTCAAGGGAGGAGGCGG<br>ATCTGGAGGCGGAGGCAGCGGTAAGGTTGAGGAGGC<br>TGTGAAAAAAGCCGAGAAGGCCCTGGTGGCAGCGGC<br>AAAGGCTGTCGCCGCCAAGCTGTTGAAGGAGCTGAA<br>GTAATAAGGTACCAAGCACCACCATCACCATCACTA<br>ATGAGATATCTC | CCGATCCAGCCTCCGGACTCTAGAGCGGCCGCCACCATGGGGAAAGTGGAAGAAGCCGTCAAAAAGG<br>CGGAAAAAGCGTTGGTCGCTGCCGCAAAAGCTGTAGCGGCAAAGTTGCTTAAAGAGCTGAAGGGTGG<br>GGGTGGGAGTGGTGGGGGCGGTAGTGGAAAGGTGGAGGAGGCAGTCAAAAAGGCAGAAAAAGCACTG<br>GTCGCAGCCGCTAAGGCCGTGGCTGCAAAGCTGTTGAAAGAGTTGAAAGGTGGCGGCGGCTCCGGCG<br>GCGGGGGATCTACCATGACAGAAAAAGAGATCGTCGATTACGTTGCGAGCCAGGTGACCACAGCCAA<br>AAAACTGAGAGGCGGGGTCGTGTTTGTTGATGAGGTGCCTAAGGGCCTCACCGGGAAGCTCGATGCC<br>AGGAAGATCCGGGAAATCCTTATCAAAGCAAAGAAGGGAGGGAAGATTGCTGTCAATTCCGGTACCA<br>AGCACCACCATCACCATCACTAA |
| 2-<br>11 | EHGKGSGGGSGGGSGKVK<br>EAAKELEKELVKGNKEAV<br>KKAKEKLEGGGGSGGGGS<br>GKVKEAAKELEKELVKGN<br>KEAVKKAKEKLE | PIQPPDSRAAATMGKVKEAAKE<br>LEKELVKGNKEAVKKAKEKLEG<br>GGGSGGGGSGKVKEAAKELEKE<br>LVKGNKEAVKKAKEKLEGGGGS<br>GGGGSTMTEKEIVDYVASQVTT<br>AKKLRGGVVFVDEVPKGLTGKL<br>DARKIREILIKAKKGGKIAVNS<br>GTKHHHHHH | CGAGCATGGGAAAGGCTCTGGAGGGGGATCTGGAGG<br>GGGATCCGGGAAGGTGAAGGAAGCTGCCAAGGAACT<br>CGAAAAGGAACTGGTCAAAGGCAATAAGGAAGCCGT<br>GAAGAAAGCCAAGGAAAAGCTCGAAGGAGGCGGGGG<br>CTCCGGGGGCGGGGGTAGCGGTAAGGTTAAGGAGGC<br>AGCTAAGGAACTGGAGAAAGAACTCGTGAAAGGGAA<br>TAAAGAAGCTGTGAAAAAAGCGAAAGAGAAACTGGA<br>ATAATAAGGTACCAAGCACCACCATCACCATCACTA<br>ATGAGATATCTC | CCGATCCAGCCTCCGGACTCTAGAGCGGCCGCCACCATGGGCAAGGTGAAAGAGGCAGCTAAGGAGC<br>TCGAAAAAGAGCTTGTGAAGGGGAACAAAGAGGCCGTGAAAAAGGCCAAGGAAAAACTGGAGGGTGG<br>CGGCGGCAGCGGGGGCGGCGGTTCCGGTAAGGTCAAAGAAGCAGCCAAAGAACTGGAGAAAGAGCTT<br>GTAAAAGGTAATAAAGAGGCAGTGAAGAAGGCGAAAGAGAAACTCGAAGGAGGAGGTGGTAGCGGGG<br>GCGGAGGCTCTACAATGACTGAGAAAGAGATTGTGGACTACGTTGCAAGCCAGGTCACTACAGCGAA<br>AAAGCTGAGGGGAGGTGTCGTCTTTGTGGATGAGGTGCCTAAGGGCCTGACAGGTAAACTCGATGCC<br>CGCAAGATACGCGAGATCCTGATCAAGGCCAAAAAGGGCGGTAAGATTGCTGTGAATTCCGGTACCA<br>AGCACCACCATCACCATCACTAA |

| | | | | |
|---|---|---|---|---|
| 2-12 | EHGKGSGGGSGGGSEKVKEAVKKLEEALKKAVEKAAKELEKKLKGGGGSGGGGSEKVKEAVKKLEEALKKAVEKAAKELEKKLK | PIQPPDSRAAATMEKVKEAVKKLEEALKKAVEKAAKELEKKLKGGGGSGGGGSEKVKEAVKKLEEALKKAVEKAAKELEKKLKGGGGSGGGGSTMTEKEIVDYVASQVTTAKKLRGGVVFVDEVPKGLTGKLDARKIREILIKAKKGGKIAVNSGTKHHHHHH | CGAGCATGGGAAAGGCTCTGGAGGGGGATCTGGAGGGGGATCCGAGAAAGTGAAAGAGGCAGTGAAGAAACTTGAAGAGGCACTGAAAAAGGCCGTCGAAAAAGCAGCGAAGGAGCTGGAGAAGAAGCTGAAGGGTGGTGGGGGTCAGGGGGAGGAGGGAGCGAGAAGGTGAAAGAGGCTGTCAAGAAGCTCGAAGAGGCATTGAAAAAAGCGGTCGAGAAGGCTGCGAAGGAATTGGAAAAGAAGCTGAAATAATAAGGTACCAAGCACCACCATCACCATCACTAATGAGATATCTC | CCGATCCAGCCTCCGGACTCTAGAGCGGCCGCCACCATGGAGAAGGTCAAGGAGGCCGTAAAAAAGCTCGAGGAGGCACTCAAAAAAGCTGTGGAGAAGGCCGCCAAAGAACTCGAAAAAAAACTGAAGGCGGGGGCGGCTCCGGAGGCGGAGGTTCTGAGAAAGTCAAGGAAGCCGTCAAGAAACTCGAGGAAGCCCTGAAAAAAGCTGTAGAGAAGGCAGCAAAGGAGCTCGAAAAGAAGCTGAAGGGTGGCGGAGGCTCCGGCGGAGGTGGATCTACTATGACCGAGAAAGAAATAGTGGATTATGTCGCTTCTCAGGTGACGACCGCCAAGAAGCTGCGCGGGGGAGTCGTATTCGTGGATGAGGTGCCAAAGGGCCTCACTGGAAAGCTCGATGCAAGAAAAATCAGGGAAATCCTTATCAAGGCCAAGAAGGGCGGTAAGATTGCGGTAAACTCAGGTACCAAGCACCACCATCACCATCACTAA |
| 2-13 | EHGKGSGGGSGGGSAKAAAAVAATEAALKAAVAAAAKALAAALAGGGGSGGGGSAKAAAAVAATEAALKAAVAAAAKALAAALA | PIQPPDSRAAATMAKAAAAVAATEAALKAAVAAAAKALAAALAGGGGSGGGGSAKAAAAVAATEAALKAAVAAAAKALAAALAGGGGSGGGGSTMTEKEIVDYVASQVTTAKKLRGGVVFVDEVPKGLTGKLDARKIREILIKAKKGGKIAVNSGTKHHHHHH | CGAGCATGGGAAAGGCTCTGGAGGGGGATCTGGAGGGGGATCCGCTAAAGCCGCCGCCGCCGTAGCTGCTACCGAAGCCGCTTTGAAAGCTGCAGTAGCTGCCGCCGCCAAAGCCCTTGCTGCTGCTCTGGCCGGTGGCGGCGGATCAGGAGGCGGCGGCAGTGCAAAGGCCGCTGCCGCCGTGGCGGCTACAGAGGCAGCACTGAAAGCCGCCGTGGCTGCCGCGGCTAAAGCTCTGGCTGCAGCCCTGGCTTAATAAGGTACCAAGCACCACCATCACCATCACTAATGAGATATCTC | CCGATCCAGCCTCCGGACTCTAGAGCGGCCGCCACCATGGCTAAAGCTGCCGCCGCCGTGGCGGCCACCGAGGCCGCACTTAAAGCCGCCGTAGCGGCTGCAGCCAAGGCCTTGGCTGCTGCCTTGGCCGGCGGAGGGGGCTCCGGTGGCGGAGGCAGTGCTAAAGCAGCCGCAGCGGTCGCAGCCACAGAGGCCGCGTTGAAGGCAGCCGTGGCTGCTGCGGCCAAAGCCCTGGCTGCTGCCCTTGCCGGGGGTGGCGGGAGCGGAGGTGGAGGGAGTACTATGACCGAGAAAGAGATTGTCGACTACGTCGCGTCCCAGGTCACCACTGCCAAAAAGCTGAGGGGAGGCGTCGTGTTTGTGGACGAAGTTCCGAAAGGGTTGACTGGCAAACTCGATGCAAGGAAAATCCGGGAGATACTGATAAAAGCGAAAAAGGGTGGAAAAATAGCTGTGAACTCTGGTACCAAGCACCACCATCACCATCACTAA |
| 2-14 | EHGKGSGGGSGGGSGKAVLAAVVGTIIGIIGAIIAGIVAGLIALGGGGSGGGGSGKAVLAAVVGTIIGIIGAIIAGIVAGLIAL | PIQPPDSRAAATMGKAVLAAVVGTIIGIIGAIIAGIVAGLIALGGGGSGGGGSGKAVLAAVVGTIIGIIGAIIAGIVAGLIALGGGGSGGGGSTMTEKEIVDYVASQVTTAKKLRGGVVFVDEVPKGLTGKLDARKIREILIKAKKGGKIAVNSGTKHHHHHH | CGAGCATGGGAAAGGCTCTGGAGGGGGATCTGGAGGGGGATCCGGTAAAGCAGTGCTGGCTGCCGTGGTCGGAACTATTATCGGCATCATCGGCGCAATTATCGCCGGTATCGTTGCGGGGCTTATAGCTCTGGGCGGCGGAGGCTCTGGGGGAGGCGGGTCTGGGAAGGCCGTCCTCGCTGCAGTTGTCGGCACAATCATTGGGATTATAGGGGCCATAATTGCTGGCATCGTCGCTGGACTGATCGCACTGTAATAAGGTACCAAGCACCACCATCACCATCACTAATGAGATATCTC | CCGATCCAGCCTCCGGACTCTAGAGCGGCCGCCACCATGGGCAAAGCCGTTCTGGCTGCCGTAGTCGGTACAATTATTGGCATTATTGGCGCAATTATTGCAGGTATTGTTGCAGGGCTGATAGCCCTGGGTGGAGGAGGATCTGGTGGCGGTGGGTCTGGGAAGGCCGTCCTGGCCGCAGTCGTTGGCACGATAATCGGTATTATCGGAGCCATCATTGCCGGAATAGTCGCGGGTCTGATCGCTCTCGGGGGCGGCGGCAGTGGAGGGGGCGGGTCCACGATGACGGAAAAGGAGATCGTCGATTACGTCGCAAGTCAGGTGACCACAGCTAAAAAGCTGAGGGGCGGCGTCGTTTTTGTGGACGAGGTCCCTAAGGGGCTCACAGGAAAGCTCGATGCACGGAAGATCCGAGAGATCCTTATTAAAGCCAAGAAAGGAGGGAAGATTGCCGTGAACAGTGGTACCAGCACCACCATCACCATCACTAA |
| 2-15 | EHGKGSGGGSGGGSDRLDALLAALRAKAEAALAALLEELRAALAGGGGSGGGGSDRLDALLAALRAKAEAALAALLEELRAALA | PIQPPDSRAAATMDRLDALLAALRAKAEAALAALLEELRAALAGGGGSGGGGSDRLDALLAALRAKAEAALAALLEELRAALAGGGGSGGGGSTMTEKEIVDYVASQVTTAKKLRGGVVFVDEVPKGLTGKLDARKIREILIKAKKGGKIAVNSGTKHHHHHH | CGAGCATGGGAAAGGCTCTGGAGGGGGATCTGGAGGGGGATCCGATCGGCTCGATGCCCTGCTGGCAGCCCTCAGGGCAAAAGCTGAAGCTGCACTTGCTGCCCTGCTCGAGGAACTCCGAGCTGCACTGGCCGGAGGTGGTGGGAGTGGTGGTGGGGGGAGCGATAGGCTTGACGCCCTTCTGGCTGCCCTTCGCGCCAAGGCAGAGGCCGCACTCGCAGCACTGCTTGAAGAATTGCGAGCAGCACTGGCATAATAAGGTACCAAGCACCACCATCACCATCACTAATGAGATATCTC | CCGATCCAGCCTCCGGACTCTAGAGCGGCCGCCACCATGGATAGGTTGGATGCTCTCTTGGCAGCCCTGCGGGCAAAAGCAGAAGCAGCTCTCGCAGCTCTTTTGGAGGAGCTGCGCGCAGCTCTGGCCGGCGGAGGCGGTTCCGGTGGTGGGGGAAGTGACCGGCTTGATGCGCTTTTGGCCGCCCTGCGCGCCAAGGCGGAGGCTGCTCTCGCCGCACTTCTGGAAGAACTGCGCGCAGCTCTGGCGGGTGGTGGTGGCTCCGGTGGCGGCGGAAGCACTATGACCGAGAAGGAGATCGTTGATTACGTCGCCTCACAGGTGACAACCGCGAAAAAGTTGAGAGGCGGGGTGGTCTTTGTAGATGAGGTGCCAAAAGGCCTGACTGGTAAGCTTGATGCTCGGAAGATACGAGAAATACTGATCAAGGCCAAAAAGGGTGGGAAGATTGCCGTTAACTCCGGTACCAAGCACCACCATCACCATCACTAA |
| 2-16 | EHGKGSGGGSGGGSEALEALLAAAEARLRAALAELAAELEAALAGGGGSGGGGSEALEALLAAAEARLRAALAELAAELEAALA | PIQPPDSRAAATMEALEALLAAAEARLRAALAELAAELEAALAGGGGSGGGGSEALEALLAAAEARLRAALAELAAELEAALAGGGGSGGGGSTMTEKEIVDYVASQVTTAKKLRGGVVFVDEVPKGLTGKLDARKIREILIKAKKGGKIAVNSGTKHHHHHH | CGAGCATGGGAAAGGCTCTGGAGGGGGATCTGGAGGGGGATCCGAAGCTCTTGAGGCGTTGCTCGCGGCTGCTGAAGCTAGGTTGCGAGCAGCTCTGGCTGAACTGGCCGCCGAGCTGGAGGCCGCATTGGCTGGAGGAGGCGGTAGCGGGGGCGGAGGGTCTGAGGCCCTGGAGGCCCTGCTGGCCGCTGCCGAGGCAAGATTGAGAGCGGCACTGGCAGAGCTGGCCGCTGAACTCGAGGCCGCATTGGCCTAATAAGGTACCAAGCACCACCATCACCATCACTAATGAGATATCTC | CCGATCCAGCCTCCGGACTCTAGAGCGGCCGCCACCATGGAAGCCCTTGAGGCGCTTCTGGCTGCAGCAGAGGCCAGGCTCCGGGCAGCATTGGCAGAGCTTGCTGCCGAGCTCGAGGCTGCCCTGGCTGGAGGCGGCGGATCTGGAGGAGGAGGTTCCGAAGCACTGGAGGCCCTGTTGGCCGCTGCGGAAGCGCGACTGCGAGCTGCTCTGGCTGAGCTCGCTGCTGAGTTGGAAGCAGCGCTGGCCGGCGGGGGAGGGAGCGGAGGCGGTGGAAGTACAATGACCGAAAAAGAGATTGTCGATTATGTCGCCTCTCAAGTGACCACTGCTAAGAAGTTGAGGGGAGGGGTCGTGTTCGTCGATGAGGTGCCCAAGGGACTGACGGGCAAGCTGGACGCGAGGAAGATCCGAGAGATCTTGATTAAAGCAAAAAAGGGCGGCAAGATAGCAGTGAACTCCGGTACCAAGCACCACCATCACCATCACTAA |

| | | | | |
|---|---|---|---|---|
| 3-1 | EHGKGSGGGSGGGSEEAA<br>KFVKEEGKKAVEYVKKEG<br>QEAVKFVKGGGGSGGGGS<br>EEAAQFVEKQNKKAVEYV<br>KEEYEKAVKFVK | PIQPPDSRAAATMEEAAKFVKE<br>EGKKAVEYVKKEGQEAVKFVKG<br>GGGSGGGGSEEAAQFVEKQNKK<br>AVEYVKEEYEKAVKFVKGGGGS<br>GGGGSTMTEKEIVDYVASQVTT<br>AKKLRGGVVFVDEVPKGLTGKL<br>DARKIREILIKAKKGGKIAVNS<br>GTKHHHHHH | CGAGCATGGGAAAGGCTCTGGAGGGGGATCTGGAGG<br>GGGATCCGAGGAAGCCGCTAAATTCGTCAAAGAGGA<br>GGGAAAAAAGGCCGTGGAATACGTTAAGAAGGAAGG<br>CCAGGAAGCCGTCAAATTTGTCAAAGGAGGAGGAGG<br>ATCTGGCGGCGGAGGGAGCGAGGAAGCTGCACAGTT<br>CGTCGAGAAGCAAAACAAGAAAGCTGTGGAGTACGT<br>GAAGGAGGAATACGAGAAAGCGGTGAAGTTTGTGAA<br>ATAATAAGGTACCAAGCACCACCATCACCATCACTA<br>ATGAGATATCTC | CCGATCCAGCCTCCGGACTCTAGAGCGGCCGCCACCATGGAGGAGGCTGCCAAGTTCGTAAAAGAGG<br>AGGGCAAGAAGGCAGTAGAGTACGTTAAGAAAGAGGGCCAGGAAGCCGTGAAATTCGTGAAGGGAGG<br>AGGGGGCAGTGGCGGGGGAGGCAGCGAGGAGGCCGCTCAATTTGTGGAGAAGCAAAATAAAAAAGCC<br>GTGGAGTACGTCAAGGAAGAGTATGAAAAAGCCGTCAAATTCGTTAAAGGAGGTGGGGTAGCGGCG<br>GTGGAGGTAGCACCATGACCGAGAAGGAAATTGTCGATTATGTGGCCTCCCAGGTTACAACCGCTAA<br>GAAGCTGCGGGGCGGCGTCGTGTTCGTAGATGAGGTGCCAAAAGGTCTTACTGGGAAACTCGACGCA<br>CGAAAAATCCGCGAGATTCTGATCAAAGCAAAGAAGGGAGGTAAGATCGCAGTCAACTCAGGTACCA<br>AGCACCACCATCACCATCACTAA |
| 3-2 | EHGKGSGGGSGGGSGKAV<br>QAVKQATQEAVKAVKEKG<br>QRAVKAVKGGGGSGGGGS<br>EKAAKAVKQEYQEAVKAV<br>KEQGWKAAQAVA | PIQPPDSRAAATMGKAVQAVKQ<br>ATQEAVKAVKEKGQRAVKAVKG<br>GGGSGGGGSEKAAKAVKQEYQE<br>AVKAVKEQGWKAAQAVAGGGGS<br>GGGGSTMTEKEIVDYVASQVTT<br>AKKLRGGVVFVDEVPKGLTGKL<br>DARKIREILIKAKKGGKIAVNS<br>GTKHHHHHH | CGAGCATGGGAAAGGCTCTGGAGGGGGATCTGGAGG<br>GGGATCCGGGAAGGCGGTGCAAGCTGTAAAGCAGGC<br>CACACAGGAGGCCGTCAAAGCCGTCAAGGAAAAGGG<br>CCAGCGAGCCGTGAAGGCAGTCAAGGGAGGGGGAGG<br>ATCTGGGGGCGGAGGTAGCGAGAAAGCTGCCAAGGC<br>TGTGAAGCAGGAGTACCAGGAGGCTGTGAAGGCCGT<br>GAAAGAGCAGGGCTGGAAAGCTGCTCAGGCAGTCGC<br>ATAATAAGGTACCAAGCACCACCATCACCATCACTA<br>ATGAGATATCTC | CCGATCCAGCCTCCGGACTCTAGAGCGGCCGCCACCATGGGTAAAGCCGTTCAGGCCGTCAAGCAAG<br>CCACCCAGGAGGCCGTCAAAGCAGTGAAGGAAAAGGGTCAAAGAGCCGTGAAAGCCGTGAAGGGTGG<br>AGGCGGGTCCGGGGGCGGCGGCTCTGAGAAGGCTGCAAAGGCCGTAAAACAGGAATATCAAGAAGCC<br>GTCAAAGCCGTAAAAGAGCAAGGATGGAAAGCCGCTCAGGCTGTCGCAGGGGGAGGCGGGTCCGGGG<br>GAGGCGGGTCAACCATGACCGAGAAGGAGATCGTAGACTACGTCGCCTCTCAAGTTACTACGGCAAA<br>GAAACTGAGGGGCGGCGTCGTGTTTGTTGACGAGGTGCCCAAAGGGCTGACCGGAAAGCTGGATGCT<br>AGGAAGATTAGAGAGATATTGATTAAAGCAAAAAAAGGGGGTAAAATCGCAGTCAACAGCGGTACCA<br>AGCACCACCATCACCATCACTAA |
| 3-3 | EHGKGSGGGSGGGSAVEA<br>ARARLLAAIEKAKEELLA<br>VVEALAKAGGGGSGGGGS<br>AVAKAKATLKAAIEKAKE<br>ELVALVEELAAA | PIQPPDSRAAATMAVEAARARL<br>LAAIEKAKEELLAVVEALAKAG<br>GGGSGGGGSAVAKAKATLKAAI<br>EKAKEELVALVEELAAAGGGGS<br>GGGGSTMTEKEIVDYVASQVTT<br>AKKLRGGVVFVDEVPKGLTGKL<br>DARKIREILIKAKKGGKIAVNS<br>GTKHHHHHH | CGAGCATGGGAAAGGCTCTGGAGGGGGATCTGGAGG<br>GGGATCCGCTGTGGAGGCTGCAAGAGCCAGGCTGCT<br>GGCGGCCATTGAAAAGGCCAAAGAGGAACTTCTTGC<br>CGTCGTCGAAGCCCTTGCAAAAGCCGGCGGAGGGGG<br>CAGCGGAGGGGGAGGGTCCGCAGTGGCCAAAGCAAA<br>GGCTACACTGAAAGCAGCTATTGAGAAGGCTAAGGA<br>AGAGCTCGTGGCTCTTGTGGAGGAGCTCGCCGCTGC<br>GTAATAAGGTACCAAGCACCACCATCACCATCACTA<br>ATGAGATATCTC | CCGATCCAGCCTCCGGACTCTAGAGCGGCCGCCACCATGGCCGTTGAAGCAGCCAGAGCTAGACTGC<br>TTGCCGCTATCGAGAAGGCAAAAGAGGAACTGTTGGCTGTCGTGGAGGCTCTGGCAAAAGCGGGTGG<br>CGGGGGAAGTGGAGGCGGCGGCAGCGCCGTCGCCAAGGCCAAGGCTACGCTGAAAGCCGCAATTGAG<br>AAAGCTAAGGAGGAATTGGTGGCTCTGGTGGAAGAGCTTGCAGCAGCTGGGGGAGGCGGGAGCGGCG<br>GCGGAGGCAGTACCATGACAGAGAAAGAGATCGTCGATTACGTCGCATCTCAGGTTACTACAGCCAA<br>GAAATTGCGCGGCGGGGTAGTATTTGTGGACGAGGTGCCAAAGGGTCTCACGGGTAAGCTCGATGCA<br>AGGAAAATCAGAGAGATACTGATTAAGGCAAAGAAGGGAGGCAAGATCGCAGTGAATTCCGGTACCA<br>AGCACCACCATCACCATCACTAA |
| 3-4 | EHGKGSGGGSGGGSKLKE<br>VAEQAKKAIKEAAEEVIK<br>FAKKLLEEGGGGSGGGGS<br>KLKAIAEAIAKRIKEAAE<br>EVNKKIEELLKE | PIQPPDSRAAATMKLKEVAEQA<br>KKAIKEAAEEVIKFAKKLLEEG<br>GGGSGGGGSKLKAIAEAIAKRI<br>KEAAEEVNKKIEELLKEGGGGS<br>GGGGSTMTEKEIVDYVASQVTT<br>AKKLRGGVVFVDEVPKGLTGKL<br>DARKIREILIKAKKGGKIAVNS<br>GTKHHHHHH | CGAGCATGGGAAAGGCTCTGGAGGGGGATCTGGAGG<br>GGGATCCAAACTGAAAGAAGTGGCTGAGCAAGCTAA<br>AAAGGCCATTAAGGAGGCCGCAGAGGAGGTGATTAA<br>GTTTGCTAAGAAGCTGCTTGAGGAGGGTGGCGGCGG<br>CAGTGGGGGTGGAGGGAGCAAACTGAAGGCCATTGC<br>TGAAGCTATTGCTAAGCGAATCAAGGAGGCTGCAGA<br>GGAGGTGAACAAGAAGATCGAGGAGCTGCTGAAGGA<br>ATAATAAGGTACCAAGCACCACCATCACCATCACTA<br>ATGAGATATCTC | CCGATCCAGCCTCCGGACTCTAGAGCGGCCGCCACCATGAAGCTGAAGGAGGTGGCTGAACAGGCAA<br>AAAAGGCTATAAAAGAAGCCGCTGAAGAGGTAATAAAATTTGCCAAGAAGCTCTTGGAAGAAGGAGG<br>CGGCGGATCTGGCGGGGGTGGCTCTAAGTTGAAGGCGATTGCTGAAGCTATCGCAAAGCGAATTAAG<br>GAAGCCGCCGAAGAGGTAAATAAGAAGATCGAGGAGCTCCTCAAGGAAGGAGGGGGAGGCTCCGGCG<br>GTGGGGGGAGTACCATGACAGAGAAAGAAATCGTTGATTACGTTGCGTCACAGGTCACAACCGCCAA<br>GAAGCTGAGGGGAGGAGTGGTGTTCGTCGATGAAGTGCCTAAGGGCTTGACTGGAAAGCTGGATGCG<br>CGCAAAATACGAGAGATACTGATCAAAGCAAAAAAGGGAGGGAAGATTGCCGTTAACTCCGGTACCA<br>AGCACCACCATCACCATCACTAA |
| 3-5 | EHGKGSGGGSGGGSWKEA<br>ARAEMEAATAEVAKLMEK<br>AIEEIVKLGGGGSGGGGS<br>LKKKVEEIGRKATAEVAK<br>LLKKAAEAIVAA | PIQPPDSRAAATMWKEAARAEM<br>EAATAEVAKLMEKAIEEIVKLG<br>GGGSGGGGSLKKKVEEIGRKAT<br>AEVAKLLKKAAEAIVAAGGGGS<br>GGGGSTMTEKEIVDYVASQVTT<br>AKKLRGGVVFVDEVPKGLTGKL<br>DARKIREILIKAKKGGKIAVNS<br>GTKHHHHHH | CGAGCATGGGAAAGGCTCTGGAGGGGGATCTGGAGG<br>GGGATCCTGGAAAGAGGCAGCACGGGCGGAGATGGA<br>GGCCGCTACCGCTGAAGTCGCCAAGCTCATGGAAAA<br>AGCTATTGAGGAAATCGTTAAGCTCGGCGGCGGTGG<br>GTCCGGCGGCGGAGGCAGTCTCAAAAAAAAGGTCGA<br>GGAAATTGGTAGGAAAGCAACCGCCGAGGTCGCCAA<br>GCTGCTGAAAAAGGCCGCAGAAGCCATCGTGGCCGC<br>ATAATAAGGTACCAAGCACCACCATCACCATCACTA<br>ATGAGATATCTC | CCGATCCAGCCTCCGGACTCTAGAGCGGCCGCCACCATGTGGAAAGAAGCCGCACGAGCCGAGATGG<br>AGGCCGCAACCGCTGAGGTGGCTAAACTGATGGAGAAAGCAATAGAGGAGATTGTGAAGCTCGGAGG<br>CGGAGGTTCCGGAGGTGGTGGGAGCCTGAAGAAAAAGGTTGAAGAGATCGGTAGAAAGGCGACTGCA<br>GAAGTGGCAAAACTTCTGAAGAAGGCCGCGGAAGCCATTGTGGCCGCTGGCGGAGGTGGTAGCGGCG<br>GGGGCGGCTCAACAATGACCGAAAAGGAGATCGTCGACTATGTTGCAAGCCAGGTAACCACGGCCAA<br>AAAACTGCGCGGCGGCGTAGTCTTTGTTGATGAAGTCCCAAAAGGCCTGACAGGAAAGTTGGATGCC<br>AGGAAAATCAGGGAAATACTGATTAAGGCTAAGAAAGGCGGGAAAATAGCTGTCAATTCCGGTACCA<br>AGCACCACCATCACCATCACTAA |

| | | | | |
|---|---|---|---|---|
| 3-6 | EHGKGSGGGSGGGSLKETIKKILEEAKKKVQEIIDETERRVLEAGGGGSGGGGSLKEKIKKEFEQLKKKVQEILEKAGKEIIEA | PIQPPDSRAAATMLKETIKKILEEAKKKVQEIIDETERRVLEAGGGGSGGGGSLKEKIKKEFEQLKKKVQEILEKAGKEIIEAGGGGSGGGGSTMTEKEIVDYVASQVTTAKKLRGGVVFVDEVPKGLTGKLDARKIREILIKAKKGGKIAVNSGTKHHHHHH | CGAGCATGGGAAAGGCTCTGGAGGGGGATCTGGAGGGGGATCCCTCAAGGAGACAATTAAAAAAATTCTGGAAGAGGCCAAGAAGAAAGTGCAAGAAATCATTGATGAAACAGAGAGGAGGGTTCTGGAAGCAGGGGGCGGCGGGTCTGGCGGCGGTGGGTCCCTCAAGGAAAAGATCAAGAAAGAATTTGAACAGCTCAAAAAGAAGGTTCAGGAAATCCTGGAGAAGGCTGGAAAGGAAATCATTGAAGCGTAATAAGGTACCAAGCACCACCATCACCATCACTAATGAGATATCTC | CCGATCCAGCCTCCGGACTCTAGAGCGGCCGCCACCATGCTGAAGGAGACTATTAAAAAAATTCTCGAAGAGGCTAAAAAGAAGGTGCAGGAGATCATCGATGAGACCGAGAGGCGAGTGTTGGAAGCTGGAGGAGGGGGGTCCGGGGGTGGCGGCTCCCTCAAGGAGAAGATCAAAAAGGAGTTTGAGCAGCTTAAGAAGAAGGTTCAGGAGATTCTCGAAAAAGCCGGCAAGGAGATCATTGAGGCTGGCGGTGGCGGATCTGGGGGCGGGGGGAGCACTATGACTGAGAAGGAGATAGTCGATTACGTGGCCTCTCAGGTGACAACCGCCAAGAAGCTGCGCGGCGGAGTGGTGTTTGTAGATGAGGTTCCAAAAGGGTTGACCGGCAAATTGGACGCAAGAAAAATCCGAGAGATTCTGATAAAAGCCAAGAAGGGTGGCAAGATCGCCGTCAATAGCGGTACCAAGCACCACCATCACCATCACTAA |
| 3-7 | EHGKGSGGGSGGGSAEAAKEIDKAEKATIAAVKAEWQAAEQAIAGGGGSGGGGSKEAAQTIKAAGKATIAAVKAEYKALKAKIA | PIQPPDSRAAATMAEAAKEIDKAEKATIAAVKAEWQAAEQAIAGGGGSGGGGSKEAAQTIKAAGKATIAAVKAEYKALKAKIAGGGGSGGGGSTMTEKEIVDYVASQVTTAKKLRGGVVFVDEVPKGLTGKLDARKIREILIKAKKGGKIAVNSGTKHHHHHH | CGAGCATGGGAAAGGCTCTGGAGGGGGATCTGGAGGGGGATCCGCCGAGGCAGCAAAAGAAATAGACAAAGCTGAAAAAGCCACCATCGCCGCCGTTAAGGCTGAGTGGCAGGCAGCCGAACAGGCTATCGCTGGTGGCGGAGGTAGTGGTGGAGGGGGAAGCAAAGAAGCGGCCCAGACAATCAAAGCAGCGGGCAAAGCCACAATCGCAGCAGTAAAGGCCGAGTACAAGGCCCTTAAAGCCAAGATCGCCTAATAAGGTACCAAGCACCACCATCACCATCACTAATGAGATATCTC | CCGATCCAGCCTCCGGACTCTAGAGCGGCCGCCACCATGGCCGAAGCTGCTAAAGAGATTGATAAAGCCGAGAAAGCCACTATTGCTGCCGTGAAAGCCGAGTGGCAGGCCGCAGAGCAAGCCATCGCAGGTGGCGGAGGCTCCGGAGGCGGGGGAAGCAAGGAAGCCGCACAAACCATAAAGGCGGCAGGTAAAGCTACTATAGCCGCCGTGAAAGCTGAATACAAAGCACTGAAAGCCAAGATCGCCGGAGGAGGGGGGTCCGGAGGGGGTGGCTCAACCATGACAGAGAAAGAGATTGTTGACTATGTGGCAAGTCAAGTGACGACCGCCAAGAAATTGCGAGGGGGGAGTGGTTTTTGTGGATGAAGTCCCCAAAGGCCTGACTGGCAAGCTGGACGCCAGAAAGATCCGGGAAATACTGATCAAGGCTAAGAAGGGGGGGAAAATTGCGGTGAACAGCGGTACCAAGCACCACCATCACCATCACTAA |
| 3-8 | EHGKGSGGGSGGGSKKEEILKKIKQKIEKAKEELLEKVEALLKAGGGGSGGGGSKKEEILKAIKEEIEKAKEELEKEIEKLLKE | PIQPPDSRAAATMKKEEILKKIKQKIEKAKEELLEKVEALLKAGGGGSGGGGSKKEEILKAIKEEIEKAKEELEKEIEKLLKEGGGGSGGGGSTMTEKEIVDYVASQVTTAKKLRGGVVFVDEVPKGLTGKLDARKIREILIKAKKGGKIAVNSGTKHHHHHH | CGAGCATGGGAAAGGCTCTGGAGGGGGATCTGGAGGGGGATCCAAAAAGGAAGAGATTCTCAAAAAAATTAAACAAAAAATAGAGAAAGCCAAAGAGGAACTTCTGGAGAAAGTGGAGGCACTTCTGAAAGCTGGTGGGGGCGGCAGCGGGGGCGGGGGCTCTAAAAAGGAGGAGATCCTCAAAGCCATCAAAGAGGAGATTGAAAAGGCAAAGGAAGAACTCGAAAAGGAAATCGAGAAACTCCTGAAAGAGTAATAAGGTACCAAGCACCACCATCACCATCACTAATGAGATATCTC | CCGATCCAGCCTCCGGACTCTAGAGCGGCCGCCACCATGAAGAAAGAGGAGATCCTTAAGAAGATCAAACAGAAAATCGAAAAGGCTAAGGAGGAGTTGCTGGAAAAGGTCGAAGCGCTTCTGAAAGCAGGCGGAGGGGGAAGCGGCGGCGGCGGATCTAAGAAGGAGGAAATCCTGAAGGCGATTAAAGAGGAAATCGAGAAGGCAAAGGAGGAGCTGGAGAAGGAAATTGAGAAATTGCTGAAGGAGGGCGGCGGCGGGTCAGGCGGTGGAGGCTCTACTATGACCGAGAAGGAAATCGTAGATTACGTCGCGTCCCAGGTCACTACAGCGAAGAAGTTGAGGGGAGGCGTTGTGTTTGTGGACGAGGTTCCGAAGGGATTGACAGGCAAGCTGGACGCTCGCAAGATCCGCGAGATTCTTATCAAAGCAAAAAAGGGCGGGAAGATTGCAGTCAACAGCGGTACCAAGCACCACCATCACCATCACTAA |
| 3-9 | EHGKGSGGGSGGGSWEEVVRKLFAELKAELEALVAATRALLAAGGGGSGGGGSLAAAIDAAFAALKAELEALVERAEKKLLEA | PIQPPDSRAAATMWEEVVRKLFAELKAELEALVAATRALLAAGGGGSGGGGSLAAAIDAAFAALKAELEALVERAEKKLLEAGGGGSGGGGSTMTEKEIVDYVASQVTTAKKLRGGVVFVDEVPKGLTGKLDARKIREILIKAKKGGKIAVNSGTKHHHHHH | CGAGCATGGGAAAGGCTCTGGAGGGGGATCTGGAGGGGGATCCTGGGAAGAAGTTGTGCGGAAACTGTTCGCCGAGTTGAAGGCTGAACTCGAAGCCCTGGTGGCTGCCACAAGAGCCGCTCTGTTGGCCGCAGGCGGCGGAGGATCTGGAGGTGGAGGGAGTCTCGCCGCTGCCATCGACGCCGCTTTTGCAGCTCTCAAAGCTGAGCTGGAGGCTCTCGTTGAACGAGCCGAGAAAAAGCTTCTGGAAGCATAATAAGGTACCAAGCACCACCATCACCATCACTAATGAGATATCTC | CCGATCCAGCCTCCGGACTCTAGAGCGGCCGCCACCATGTGGGAGGAAGTCGTCCGCAAACTCTTCGCTGAACTCAAAGCTGAACTGGAGGCTCTCGTGGCGGCCACAAGGGCGGCACTGCTTGCCGCAGGAGGCGGTGGATCAGGCGGAGGAGGTTCTCTTGCCGCGGCCATAGACGCCGCTTTTGCCGCCCTGAAGGCCGAGTTGGAAGCTTTGGTTGAGAGAGCCGAGAAAAAATTGCTCGAAGCAGGGGGTGGGGGTTCTGGGGGCGGCGGCTCTACAATGACGGAAAAGGAGATAGTAGACTATGTGGCAAGTCAGGTGACTACCGCTAAGAAATTGCGCGGTGGCGTCGTGTTCGTGGACGAGGTCCCAAAGGGTCTGACTGGCAAGCTTGATGCGCGCAAGATTCGGGAGATCCTCATTAAAGCTAAAAAAGGAGGTAAAATTGCAGTGAACAGTGGTACCAAGCACCACCATCACCATCACTAA |
| 3-10 | EHGKGSGGGSGGGSEEKKEKLLAAAKRIAEEALKAMEAKLKAAAGGGGSGGGGSAAKVAELKAAAKAIAEEALAAMKKEVEAVF | PIQPPDSRAAATMEEKKEKLLAAAKRIAEEALKAMEAKLKAAAGGGGSGGGGSAAKVAELKAAAKAIAEEALAAMKKEVEAVFGGGGSGGGGSTMTEKEIVDYVASQVTTAKKLRGGVVFVDEVPKGLTGKLDARKIREILIKAKKGGKIAVNSGTKHHHHHH | CGAGCATGGGAAAGGCTCTGGAGGGGGATCTGGAGGGGGATCCGAGGAGAAAAAAGAGAAACTGTTGGCTGCAGCCAAACGGATTGCCGAAGAGGCCCTTAAAGCTATGGAGGCGAAGCTGAAAGCCGCTGCCGGAGGCGGGGGATCTGGCGGGGGCGGTTCCGCCGCCAAGGTTGCTGAGTTGAAAGCCGCCGCTAAGGCTATTGCCGAGGAAGCTCTTGCAGCCATGAAGAAGGAAGTGGAAGCTGTTTTTTAATAAGGTACCAAGCACCACCATCACCATCACTAATGAGATATCTC | CCGATCCAGCCTCCGGACTCTAGAGCGGCCGCCACCATGGAAGAGAAGAAAGAGAAGCTCCTGGCTGCAGCCAAACGGATTGCTGAGGAAGCTCTGAAAGCTATGGAAGCAAAATTGAAGGCTGCGGCAGGAGGAGGGGGCTCAGGCGGCGGCGGCAGTGCCGCTAAGGTCGCTGAACTTAAAGCCGCAGCGAAAGCCATTGCAGAGGAGGCCCTCGCCGCAATGAAAAAGGAGGTGGAGGCTGTATTCGGGGGCGGAGGGTCTGGAGGGGGCGGCAGCACCATGACTGAAAAAGAAATCGTTGACTACGTCGCATCACAAGTGACTACTGCCAAGAAGCTCAGAGGTGGCGTGGTCTTCGTGGATGAGGTCCCTAAAGGACTGACAGGTAAGCTCGATGCCAGGAAAATTCGCGAAATTTTGATAAAAGCTAAGAAAGGAGGCAAAATTGCTGTAAATAGCGGTACCAAGCACCACCATCACCATCACTAA |

| | | | | |
|---|---|---|---|---|
| 3-11 | EHGKGSGGGSGGGSAKITALVEKAEEELKKEVEAIFAELLAKLKGGGGSGGGGSEEIAALVAAAAEELKKEVEALFAKLKAELA | PIQPPDSRAAATMAKITALVEKAEEELKKEVEAIFAELLAKLKGGGGSGGGGSEEIAALVAAAAEELKKEVEALFAKLKAELAGGGGSGGGGSTMTEKEIVDYVASQVTTAKKLRGGVVFVDEVPKGLTGKLDARKIREILIKAKKGGKIAVNSGTKHHHHHH | CGAGCATGGGAAAGGCTCTGGAGGGGGATCTGGAGGGGGATCCGCCAAAATCACCGCTCTTGTCGAAAAGGCTGAAGAAGAGCTTAAAAAGGAGGTGGAAGCAATTTTCGCCGAGCTGCTGGCAAAGCTGAAGGGCGGAGGTGGAAGCGGAGGTGGTGGCTCAGAGGAAATAGCCGCACTGGTTGCCGCAGCAGCAGAGGAGCTCAAGAAAGAAGTTGAGGCCCTGTTTGCCAAGCTGAAAGCAGAGCTGGCCTAATAAGGTACCAAGCACCACCATCACCATCACTAATGAGATATCTC | CCGATCCAGCCTCCGGACTCTAGAGCGGCCGCCACCATGGCAAAAATTACCGCCCTGGTGGAGAAAGCTGAGGAGGAACTGAAAAAGGAGGTCGAGGCGATTTTTGCGGAGCTCCTCGCAAAACTCAAAGGTGGGGGCGGCTCCGGCGGCGGGGGCTCTGAGGAGATAGCAGCCTTGGTGGCTGCGGCTGCGGAAGAATTGAAAAAGGAGGTGGAAGCCCTGTTTGCTAAGCTGAAGGCAGAATTGGCTGGGGGCGGAGGTTCCGGCGGCGGCGGATCTACTATGACCGAGAAAGAGATCGTCGATTACGTCGCTAGCCAGGTGACAACTGCGAAAAAGTTGCGGGGAGGCGTCGTCTTTGTGGACGAGGTGCCTAAAGGACTCACAGGGAAATTGGACGCCAGAAAAATCAGAGAGATTCTCATCAAGGCCAAAAAAGGGGGGAAGATAGCGGTCAACAGTGGTACCAAGCACCACCATCACCATCACTAA |
| 3-12 | EHGKGSGGGSGGGSFKEVVLKEAEEAAEEAKKIIEKYAKKAAKAGGGGSGGGGSVEEEILKEGKKAAEEAKKIIKEKGKKAAQA | PIQPPDSRAAATMFKEVVLKEAEEAAEEAKKIIEKYAKKAAKAGGGGSGGGGSVEEEILKEGKKAAEEAKKIIKEKGKKAAQAGGGGSGGGGSTMTEKEIVDYVASQVTTAKKLRGGVVFVDEVPKGLTGKLDARKIREILIKAKKGGKIAVNSGTKHHHHHH | CGAGCATGGGAAAGGCTCTGGAGGGGGATCTGGAGGGGGATCCTTCAAAGAGGTTGTTCTCAAAGAGGCAGAAGAGGCCGCTGAGGAGGCTAAGAAAATTATCGAGAAGTACGCTAAAAAAGCTGCCAAGGCAGGAGGGGGCGGGAGTGGAGGCGGTGGGAGCGTGGAGGAGGAAATTTTGAAGGAAGGGAAGAAAGCCGCCGAGGAGGCTAAAAAGATAATCAAGGAAAAGGGTAAGAAAGCGGCACAGGCATAATAAGGTACCAAGCACCACCATCACCATCACTAATGAGATATCTC | CCGATCCAGCCTCCGGACTCTAGAGCGGCCGCCACCATGTTCAAGGAAGTGGTGCTGAAGGAAGCTGAGGAGGCTGCAGAGGAGGCTAAGAAAATAATTGAGAAATATGCCAAGAAGGCTGCAAAAGCAGGCGGGGGTGGGAGTGGCGGAGGTGGCTCTGTGGAGGAAGAGATCCTGAAGGAAGGGAAAAAAGCCGCTGAGGAGGCTAAAAAGATAATAAAAGAGAAGGGGAAGAAGGCTGCCCAGGCCGGCGGAGGAGGATCTGGGGGCGGAGGGTCCACCATGACGGAAAAGGAGATCGTTGACTACGTGGCCAGCCAGGTCACGACCGCAAAGAAGTTGAGAGGAGGGGTCGTGTTTGTGGATGAAGTGCCCAAGGGGCTGACCGGAAAACTGGACGCAAGAAAAATCCGGGAGATTCTCATTAAGGCCAAAAAAGGTGGCAAGATCGCCGTCAACAGCGGTACCAAGCACCACCATCACCATCACTAA |
| 3-13 | EHGKGSGGGSGGGSEKIKKVIEKAKEKLKKAVEEIAKETLAKLKGGGGSGGGGSAKIDAVIAEALEKLKKAVEEIMEEMLKELE | PIQPPDSRAAATMEKIKKVIEKAKEKLKKAVEEIAKETLAKLKGGGGSGGGGSAKIDAVIAEALEKLKKAVEEIMEEMLKELEGGGGSGGGGSTMTEKEIVDYVASQVTTAKKLRGGVVFVDEVPKGLTGKLDARKIREILIKAKKGGKIAVNSGTKHHHHHH | CGAGCATGGGAAAGGCTCTGGAGGGGGATCTGGAGGGGGATCCGAAAAAATCAAGAAGGTGATCGAGAAGGCCAAGGAAAAACTGAAAAAGGCCGTGGAAGAAATAGCAAAAGAAACTCTCGCAAAGCTGAAAGGAGGCGGCGGATCTGGCGGTGGTGGCTCAGCCAAGATTGACGCCGTTATCGCAGAGGCTCTCGAAAAATTGAAAAAAGCGGTGGAAGAAATCATGGAAGAAATGCTGAAGGAGCTGGAATAATAAGGTACCAAGCACCACCATCACCATCACTAATGAGATATCTC | CCGATCCAGCCTCCGGACTCTAGAGCGGCCGCCACCATGGAGAAGATAAAGAAAGTAATTGAAAAGGCCAAGGAGAAGCTGAAAAAGGCCGTGGAAGAAATCGCCAAGGAAACCCTTGCAAAGTTGAAGGGAGGCGGCGGAAGCGGCGGTGGCGGTTCTGCGAAAATCGATGCAGTGATAGCTGAGGCACTGGAGAAGTTGAAGAAGGCCGTGGAAGAGATTATGGAAGAAATGCTGAAGGAGTTGGAGGGCGGTGGTGGCAGCGGGGGTGGTGGATCTACGATGACCGAAAAAGAGATTGTGGATTATGTGGCATCCCAAGTCACAACGGCCAAGAAGCTGCGGGGTGGCGTCGTGTTCGTGGACGAGGTGCCCAAAGGCCTGACTGGAAAACTGGATGCACGAAAAATAAGGGAGATATTGATTAAAGCTAAAAAGGGAGGAAAGATAGCCGTCAACTCCGGTACCAAGCACCACCATCACCATCACTAA |
| 3-14 | EHGKGSGGGSGGGSAAALAAAKAATKAELAKLEAAISKEIKKLYGGGGSGGGGSEEKVKQLEAFIKAELAKLEAALAATLAALA | PIQPPDSRAAATMAAALAAAKAATKAELAKLEAAISKEIKKLYGGGGSGGGGSEEKVKQLEAFIKAELAKLEAALAATLAALAGGGGSGGGGSTMTEKEIVDYVASQVTTAKKLRGGVVFVDEVPKGLTGKLDARKIREILIKAKKGGKIAVNSGTKHHHHHH | CGAGCATGGGAAAGGCTCTGGAGGGGGATCTGGAGGGGGATCCGCAGCAGCTTTGGCGGCAGCCAAGGCTGCCACCAAGGCTGAACTGGCCAAACTCGAGGCAGCTATTAGCAAAGAAATCAAAAAGTTGTATGGCGGGGGCGGCAGCGGGGGCGGCGGAAGTGAAGAAAAGGTGAAGCAACTTGAGGCTTTCATCAAGGCAGAGCTGGCTAAATTGGAAGCGGCTCTGGCTGCCACACTTGCTGCGTTGGCCTAATAAGGTACCAAGCACCACCATCACCATCACTAATGAGATATCTC | CCGATCCAGCCTCCGGACTCTAGAGCGGCCGCCACCATGGCTGCAGCTTTGGCAGCAGCGAAGGCAGCCACCAAAGCGGAGCTCGCTAAACTGGAAGCCGCTATAAGTAAGGAGATTAAGAAATTGTACGGGGGAGGTGGCTCCGGCGGTGGCGGAAGTGAAGAGAAGGTGAAGCAGCTGGAGGCTTTCATCAAGGCGGAGCTGGCTAAGCTGGAGGCCGCTCTGGCTGCAACTCTTGCAGCTTTGGCCGGAGGGGGTGGGTCTGGAGGAGGAGGTAGCACTATGACAGAGAAAGAGATTGTTGACTATGTGGCAAGCCAGGTGACTACAGCTAAAAAACTGAGGGGGGGCGTCGTCTTTGTGGATGAAGTGCCTAAGGGCCTGACAGGAAAACTTGATGCCAGAAAGATAAGAGAAATTCTCATCAAAGCGAAGAAAGGCGGGAAGATCGCCGTAAACTCTGGTACCAAGCACCACCATCACCATCACTAA |
| 3-15 | EHGKGSGGGSGGGSKAKAITDAAKKAAQKAAAQVEKAVAAAAQAGGGGSGGGGSRAEAIKKAGKKAAQKAAAQVVKAVAEAAKK | PIQPPDSRAAATMKAKAITDAAKKAAQKAAAQVEKAVAAAAQAGGGGSGGGGSRAEAIKKAGKKAAQKAAAQVVKAVAEAAKKGGGGSGGGGSTMTEKEIVDYVASQVTTAKKLRGGVVFVDEVPKGLTGKLDARKIREILIKAKKGGKIAVNSGTKHHHHHH | CGAGCATGGGAAAGGCTCTGGAGGGGGATCTGGAGGGGGATCCAAAGCCAAGGCAATCACAGACGCAGCCAAGAAAGCAGCACAGAAGGCTGCAGCTCAAGTGGAGAAGGCCGTCGCTGCCGCAGCACAGGCTGGAGGCGGTGGCTCAGGGGGTGGAGGGTCCAGAGCAGAGGCCATTAAGAAGGCTGGAAAAAAAGCGGCTCAGAAAGCCGCCGCACAGGTGGTTAAGGCCGTTGCTGAAGCAGCTAAAAAATAATAAGGTACCAAGCACCACCATCACCATCACTAATGAGATATCTC | CCGATCCAGCCTCCGGACTCTAGAGCGGCCGCCACCATGAAGGCAAAGGCTATTACGGACGCCGCCAAAAAAGCTGCTCAGAAAGCTGCCGCGCAAGTGGAAAAAGCGGTGGCAGCCGCCGCACAGGCTGGAGGCGGCGGTTCCGGCGGGGGTGGTAGCCGGGCTGAGGCTATCAAGAAAGCAGGCAAGAAAGCCGCACAGAAGGCTGCGGCCCAAGTGGTCAAGGCCGTCGCTGAAGCTGCTAAAAAAGGGGGAGGCGGGTCAGGCGGTGGAGGCTCAACCATGACCGAGAAAGAGATCGTAGACTATGTGGCTTCTCAGGTGACAACCGCGAAGAAACTGCGCGGAGGAGTAGTATTCGTGGATGAGGTGCCCAAGGGCCTGACTGGCAAGCTCGACGCCAGAAAGATCCGGGAGATCCTCATTAAAGCTAAAAAGGGCGGAAAAATAGCCGTGAACTCTGGTACCAAGCACCACCATCACCATCACTAA |

| | | | | |
|---|---|---|---|---|
| 3-16 | EHGKGSGGGSGGGSAKIKAVFAKLEKELKEAVKAVMKKAEQEAKGGGGSGGGGSGKIEEVFKKAEKELKEAVKAVVAEAEKELL | PIQPPDSRAAATMAKIKAVFAKLEKELKEAVKAVMKKAEQEAKGGGGSGGGGSGKIEEVFKKAEKELKEAVKAVVAEAEKELLGGGGSGGGGSTMTEKEIVDYVASQVTTAKKLRGGVVFVDEVPKGLTGKLDARKIREILIKAKKGGKIAVNSGTKHHHHHH | CGAGCATGGGAAAGGCTCTGGAGGGGGATCTGGAGGGGGATCCGCCAAAATAAAAGCCGTGTTTGCAAAGCTGGAAAAGGAGCTGAAGGAAGCAGTCAAAGCCGTGATGAAAAAGGCAGAGCAGGAAGCCAAGGGCGGTGGCGGTAGCGGCGGGGGTGGTTCTGGTAAGATAGAAGAGGTTTTTAAGAAAGCAGAGAAGGAGTTGAAAGAAGCAGTAAAGGCCGTGGTCGCCGAGGCTGAGAAGGAACTGCTTTAATAAGGTACCAAGCACCACCATCACCATCACTAATGAGATATCTC | CCGATCCAGCCTCCGGACTCTAGAGCGGCCGCCACCATGGCTAAAATAAAGGCCGTGTTCGCGAAGCTGGAGAAGGAGCTCAAGGAAGCTGTGAAAGCTGTGATGAAAAAGGCTGAACAAGAAGCTAAGGGCGGCGGTGGCAGCGGGGGAGGTGGCAGCGGGAAAATAGAGGAAGTCTTCAAGAAGGCCGAGAAGGAACTCAAAGAAGCTGTGAAGGCCGTGGTTGCCGAGGCTGAAAAGGAACTCCTCGGCGGGGGTGGGAGCGGAGGAGGCGGGTCCACAATGACCGAAAAGGAGATCGTCGATTATGTGGCTTCTCAGGTGACCACCGCTAAAAAGCTGCGCGGAGGGGTGGTGTTTGTGGACGAGGTGCCTAAAGGCTTGACTGGAAAGTTGGATGCCCGGAAAATAAGAGAAATACTCATCAAAGCCAAAAAGGGAGGCAAAATCGCAGTCAACAGCGGTACCAAGCACCACCATCACCATCACTAA |
| 3-17 | EHGKGSGGGSGGGSALQAALDAVAEAVTAQALALLKEALRKVIAGGGGSGGGGSELLAKLEAVAEEVTAQALALLKAALAAVAA | PIQPPDSRAAATMALQAALDAVAEAVTAQALALLKEALRKVIAGGGGSGGGGSELLAKLEAVAEEVTAQALALLKAALAAVAAGGGGSGGGGSTMTEKEIVDYVASQVTTAKKLRGGVVFVDEVPKGLTGKLDARKIREILIKAKKGGKIAVNSGTKHHHHHH | CGAGCATGGGAAAGGCTCTGGAGGGGGATCTGGAGGGGGATCCGCGTTGCAGGCCGCTCTCGACGCGGTAGCCGAAGCGGTCACCGCACAAGCTCTGGCCCTGCTGAAAGAGGCACTTCGGAAGGTCATTGCTGGAGGTGGCGGGAGCGGGGGTGGCGGTAGTGAACTTCTCGCCAAGCTTGAGGCAGTTGCCGAAGAGGTAACGGCACAGGCTCTTGCACTCCTGAAGGCAGCCCTCGCTGCCGTTGCTGCATAATAAGGTACCAAGCACCACCATCACCATCACTAATGAGATATCTC | CCGATCCAGCCTCCGGACTCTAGAGCGGCCGCCACCATGGCTCTCCAGGCTGCTCTCGACGCGGTTGCAGAGGCAGTAACCGCACAGGCTCTCGCCCTGCTTAAAGAGGCTTTGAGGAAGGTAATAGCCGGCGGAGGCGGAAGTGGCGGTGGAGGAAGCGAACTGCTCGCCAAGCTTGAAGCTGTTGCAGAGGAAGTGACAGCACAAGCCCTGGCACTGCTTAAGGCAGCTCTGGCCGCAGTCGCAGCAGGAGGCGGGGGTAGCGGTGGCGGCGGTAGTACAATGACCGAGAAAGAGATCGTCGATTACGTGGCATCACAGGTGACTACCGCCAAAAAGCTGCGCGGCGGCGTGGTGTTCGTGGATGAGGTGCCCAAAGGGCTGACCGGGAAGCTTGATGCCCGCAAAATTCGCGAGATTCTCATCAAAGCCAAGAAAGGTGGCAAAATAGCGGTCAACAGCGGTACCAAGCACCACCATCACCATCACTAA |

**Table S4.** Backbone RMSD statistics of the six designed helical bundles during MD simulations at 30 °C. Mean and maximum backbone RMSD values relative to the initial AlphaFold3 (AF3) model and the trajectory-averaged structure (Avg) are reported in Å. Values are given separately for each of the three independent MD simulations, in the order replicate 1 / replicate 2 / replicate 3.

| | Mean RMSD AF3 (Å) | Max RMSD AF3 (Å) | Mean RMSD Avg (Å) | Max RMSD Avg (Å) |
|---|---|---|---|---|
| Homomer-1 | 2.47 / 2.69 / 2.46 | 3.56 / 4.04 / 3.71 | 1.84 / 1.83 / 1.61 | 2.75 / 3.37 2.88 |
| Homomer-2 | 1.94 / 2.03 / 2.20 | 3.39 / 3.25 / 3.53 | 1.05 / 1.13 / 1.14 | 2.17 / 2.23 / 2.78 |
| Homomer-3 | 1.61 / 1.50 / 1.55 | 2.15 / 2.25 / 2.18 | 0.79 / 1.02 / 0.93 | 2.23 / 2.01 / 2.13 |
| Homomer-4 | 2.09 / 2.66 / 1.90 | 3.67 / 3.59 / 3.52 | 1.35 / 0.93 / 1.42 | 2.45 / 2.78 / 2.26 |
| Heteromer-1 | 1.37 / 1.35 / 1.35 | 2.31 / 2.16 / 2.20 | 1.05 / 1.07 / 1.03 | 1.87 / 1.75 / 1.91 |
| Heteromer-2 | 0.95 / 1.04 / 0.95 | 1.92 / 2.30 / 1.86 | 0.77 / 0.90 / 0.78 | 1.55 / 2.10 / 1.63 |

**Table S5.** Backbone RMSD statistics of the six designed helical bundles during MD simulations at 40 °C. Mean and maximum backbone RMSD values relative to the initial AlphaFold3 (AF3) model and the trajectory-averaged structure (Avg) are reported in Å. Values are given separately for each of the three independent MD simulations, in the order replicate 1 / replicate 2 / replicate 3.

| | Mean RMSD AF3 (Å) | Max RMSD AF3 (Å) | Mean RMSD Avg (Å) | Max RMSD Avg (Å) |
|---|---|---|---|---|
| Homomer-1 | 3.02 / 3.24 / 2.83 | 4.55 / 4.33 / 3.93 | 1.93 / 2.24 / 1.75 | 3.53 / 3.21 / 3.52 |
| Homomer-2 | 2.14 / 2.03 / 2.21 | 3.68 / 3.54 / 3.62 | 1.20 / 1.10 / 1.19 | 2.51 / 2.47 / 2.59 |
| Homomer-3 | 1.55 / 1.90 / 1.54 | 2.17 / 3.03 / 2.23 | 0.96 / 1.26 / 0.95 | 2.02 / 3.12 / 1.92 |
| Homomer-4 | 2.32 / 2.65 / 2.70 | 3.61 / 3.65 / 3.62 | 1.19 / 0.95 / 1.00 | 2.61 / 2.69 / 2.95 |
| Heteromer-1 | 1.44 / 1.89 / 1.50 | 2.44 / 3.46 / 2.47 | 1.08 / 1.58 / 1.09 | 1.84 / 2.97 / 2.28 |
| Heteromer-2 | 1.00 / 1.02 / 1.50 | 2.06 / 2.09 / 2.43 | 0.87 / 0.86 / 0.98 | 1.84 / 1.92 / 1.91 |

**Table S7.** PDB identifiers of coiled-coil structures from the SamCC-Turbo database used for the analysis of coiled-coil geometric parameters.

| |
|---|
| 1BB1, 3R3K, 3R46, 3R47, 3R48, 3R4A, 3R4H, 3RA3, 4DZK, 4DZL, 4DZM, 4DZN, 4H7R, 4H8F, 4H8G, 4H8L, 4H8M, 4H8O, 4KVT, 4KVU, 4KVV, 4PN8, 4PN9, 4PNA, 4PNB, 4PND, 4TQL, 4UOS, 4UOT, 5EZ8, 5EZ9, 5EZA, 5EZC, 5EZE, 5F2Y, 5J0K, 5J10, 5J2L, 5J73, 5U59, 5U5A, 5U5B, 5U5C, 6B87, 6DKM, 6DLC, 6DLM, 6DM9, 6DMA, 6DMP, 6EIK, 6EIZ, 6G65, 6G66, 6G67, 6G68, 6G69, 6G6A, 6G6B, 6G6C, 6G6D, 6G6E, 6G6F, 6G6G, 6G6H, 6NX2, 6NXM, 6NY8, 6NZ3, 6OLO, 6OWD, 6Q5H, 6Q5I, 6Q5J, 6Q5K, 6Q5L, 6Q5M, 6Q5N, 6Q5O, 6Q5P, 6Q5Q, 6Q5R, 6Q5S, 6XXZ, 6XY0, 6XY1, 6YAZ, 6YB0, 6YB2, 6ZT1, 7A1T, 7BAS, 7BAT, 7BAU, 7BAV, 7BAW, 7BIM, 7BO8, 7BO9, 7BOA, 7L33, 7NFF, 7NFG, 7NFH, 7NFI, 7NFJ, 7NFK, 7NFL, 7NFM, 7NFN, 7NFO, 7NFP, 7Q1Q, 7Q1R, 7Q1S, 7Q1T, 7QDK, 8A09, 8A3G, 8A3I, 8A3J, 8A3K, 8BCS |

**Table S6.** X-ray Data collection and refinement statistics for Heteromer-2

| | |
|---|---|
| ***Data collection*** | |
| Source | DESY PETRA III P14 |
| Wavelength (Å) | 0.9762 |
| Resolution (Å) | 82.13-1.92 (2.44-1.92) |
| Resolution iso (Å) | |
| Resolution aniso (Å) | 2.46, 3.25, 2.02 |
| Space group | P 1 |
| Unit cell dimensions (Å;°) | 25.09, 85.43, 91.91; 63.33, 89.45, 89.52 |
| Unique reflections | 16503 (825) |
| Multiplicity | 2.7 (2.2) |
| Completeness (%)* spherical/ellipsoidal | 31.6 (3.1) / 71.6 (30.3) |
| Mean I/σ(I) | 3.3 (1.6) |
| R-meas | 0.214 (0.631) |
| R-merge | 0.172 (0.495) |
| R-pim | 0.124 (0.387) |
| $CC_{1/2}$ | 0.991 (0.811) |
| Wilson B ($Å^2$) | 5.17 |
| ***Refinement*** | |
| Resolution (Å) | 76.34-1.92 (2.04-1.92) |
| R-work | 0.3111 (0.3962) |
| R-free | 0.3640 (1000000000.00) |
| Number of non-hydrogen atoms, average B-value ($Å^2$) | |
| Protein | 5941, 3.84 |
| Heterogen | 0, 0.00 |
| Solvent | 236, 3.45 |
| Rmsd bonds (Å), angles (°) | 0.002, 0.355 |
| Ramachandran favored, allowed, outliers (%) | 99.8, 0.1, 0.0 |
| Rotamer outliers (%) | 0.0 |
| MolProbity clashscore | 3.1 |

*If values for ellipsoidal completeness are specified, anisotropic truncation has been used. The first line refers to the spherical and the second line to the ellipsoidal completeness.

**Table S8.** DNA sequences of the pcDSec2 plasmid with and without the N-terminal split-Luciferase (nLuc). The NotI restriction site (GCGGCCGC) is colored yellow, the KpnI restriction site (GGTACC) is colored green, the BamHI restriction site (GGATCC) is colored red. nLuc is colored blue.

| ID | DNA |
|---|---|
| pcDSec2 | GTTAGGCGTTTTGCGCTGCTTCGCGATGTACGGGCCAGATATACGCGTTGACATTGATTATTGACTAGTTA<br>TTAATAGTAATCAATTACGGGGTCATTAGTTCATAGCCCATATATGGAGTTCCGCGTTACATAACTTACGG<br>TAAATGGCCCGCCTGGCTGACCGCCCAACGACCCCCGCCCATTGACGTCAATAATGACGTATGTTCCCATA<br>GTAACGCCAATAGGGACTTTCCATTGACGTCAATGGGTGGAGTATTTACGGTAAACTGCCCACTTGGCAGT<br>ACATCAAGTGTATCATATGCCAAGTACGCCCCCTATTGACGTCAATGACGGTAAATGGCCCGCCTGGCATT<br>ATGCCCAGTACATGACCTTATGGGACTTTCCTACTTGGCAGTACATCTACGTATTAGTCATCGCTATTACC<br>ATGGTGATGCGGTTTTGGCAGTACATCAATGGGCGTGGATAGCGGTTTGACTCACGGGGATTTCCAAGTCT<br>CCACCCCATTGACGTCAATGGGAGTTTGTTTTGGCACCAAAATCAACGGGACTTTCCAAAATGTCGTAACA<br>ACTCCGCCCCATTGACGCAAATGGGCGGTAGGCGTGTACGGTGGGAGGTCTATATAAGCAGAGCTCGTTTA<br>GTGAACCGTCAGATCGCCTGGAGACGCCATCCACGCTGTTTTGACCTCCATAGAAGACACCGGGACCGATC<br>CAGCCTCCGGACTCTAGAGCGGCCGCCACCATGGATGCAATGAAGAGAGGGCTCTGCTGTGTGCTGCTGCT<br>GTGTGGAGCAGTCTTCGTTTCGCCCAGCCAGGAAATCCATGCCCGATTCAGAAGAGGGGCCCGTGAAACCG<br>GTTGATAAGGTACCAAGCACCACCATCACCATCACTAATGAGATATCTCGACAATCAACCTCTGGATTACA<br>AAATTTGTGAAAGATTGACTGGTATTCTTAACTATGTTGCTCCTTTTACGCTATGTGGATACGCTGCTTTA<br>ATGCCTTTGTATCATGCTATTGCTTCCCGTATGGCTTTCATTTTCTCCTCCTTGTATAAATCCTGGTTGCT<br>GTCTCTTTATGAGGAGTTGTGGCCCGTTGTCAGGCAACGTGGCGTGGTGTGCACTGTGTTTGCTGACGCAA<br>CCCCCACTGGTTGGGGCATTGCCACCACCTGTCAGCTCCTTTCCGGGACTTTCGCTTTCCCCCTCCCTATT<br>GCCACGGCGGAACTCATCGCCGCCTGCCTTGCCCGCTGCTGGACAGGGGCTCGGCTGTTGGGCACTGACAA<br>TTCCGTGGTGTTGTCGGGGAAGCTGACGTCCTTTCCATGGCTGCTCGCCTGTGTTGCCACCTGGATTCTGC<br>GCGGGACGTCCTTCTGCTACGTCCCTTCGGCCCTCAATCCAGCGGACCTTCCTTCCCGCGGCCTGCTGCCG<br>GCTCTGCGGCCTCTTCCGCGTCTTCGCCTTCGCCCTCAGACGAGTCGGATCTCCCTTTGGGCCGCCTCCCC<br>GCCTGGAAACGGGGGAGGCTAACTGAAACACGGAAGGAGACAATACCGGAAGGAACCCGCGCTATGACGGC<br>AATAAAAAGACAGAATAAAACGCACGGGTGTTGGGTCGTTTGTTCATAAACGCGGGGTTCGGTCCCAGGGC<br>TGGCACTCTGTCGATACCCCACCGAGACCCCATTGGGGCCAATACGCCCGCGTTTCTTCCTTTTCCCCACC<br>CCACCCCCCAAGTTCGGGTGAAGGCCCAGGGCTCGCAGCCAACGTCGGGGCGGCAGGCCCTGCCATAGCAG<br>ATCTGCGCAGCTGGGGCTCTAGGGGGTATCCCCTATACCGTCGACCTCTAGCTAGAGCTTGGCGTAATCAT<br>GGTCATAGCTGTTTCCTGTGTGAAATTGTTATCCGCTCACAATTCCACACAACATACGAGCCGGAAGCATA<br>AAGTGTAAAGCCTGGGGTGCCTAATGAGTGAGCTAACTCACATTAATTGCGTTGCGCTCACTGCCCGCTTT<br>CCAGTCGGGAAACCTGTCGTGCCAGCTGCATTAATGAATCGGCCAACGCGCGGGGAGAGGCGGTTTGCGTA<br>TTGGGCGCTCTTCCGCTTCCTCGCTCACTGACTCGCTGCGCTCGGTCGTTCGGCTGCGGCGAGCGGTATCA<br>GCTCACTCAAAGGCGGTAATACGGTTATCCACAGAATCAGGGGATAACGCAGGAAAGAACATGTGAGCAAA<br>AGGCCAGCAAAAGGCCAGGAACCGTAAAAAGGCCGCGTTGCTGGCGTTTTTCCATAGGCTCCGCCCCCCTG<br>ACGAGCATCACAAAAATCGACGCTCAAGTCAGAGGTGGCGAAACCCGACAGGACTATAAAGATACCAGGCG<br>TTTCCCCCTGGAAGCTCCCTCGTGCGCTCTCCTGTTCCGACCCTGCCGCTTACCGGATACCTGTCCGCCTT<br>TCTCCCTTCGGGAAGCGTGGCGCTTTCTCATAGCTCACGCTGTAGGTATCTCAGTTCGGTGTAGGTCGTTC<br>GCTCCAAGCTGGGCTGTGTGCACGAACCCCCCGTTCAGCCCGACCGCTGCGCCTTATCCGGTAACTATCGT<br>CTTGAGTCCAACCCGGTAAGACACGACTTATCGCCACTGGCAGCAGCCACTGGTAACAGGATTAGCAGAGC<br>GAGGTATGTAGGCGGTGCTACAGAGTTCTTGAAGTGGTGGCCTAACTACGGCTACACTAGAAGAACAGTAT<br>TTGGTATCTGCGCTCTGCTGAAGCCAGTTACCTTCGGAAAAAGAGTTGGTAGCTCTTGATCCGGCAAACAA<br>ACCACCGCTGGTAGCGGTGGTTTTTTTGTTTGCAAGCAGCAGATTACGCGCAGAAAAAAAGGATCTCAAGA<br>AGATCCTTTGATCTTTTCTACGGGGTCTGACGCTCAGTGGAACGAAAACTCACGTTAAGGGATTTTGGTCA<br>TGAGATTATCAAAAAGGATCTTCACCTAGATCCTTTTAAATTAAAAATGAAGTTTTAAATCAATCTAAAGT<br>ATATATGAGTAAACTTGGTCTGACAGTTACCAATGCTTAATCAGTGAGGCACCTATCTCAGCGATCTGTCT<br>ATTTCGTTCATCCATAGTTGCCTGACTCCCCGTCGTGTAGATAACTACGATACGGGAGGGCTTACCATCTG<br>GCCCCAGTGCTGCAATGATACCGCGAGACCCACGCTCACCGGCTCCAGATTTATCAGCAATAAACCAGCCA<br>GCCGGAAGGGCCGAGCGCAGAAGTGGTCCTGCAACTTTATCCGCCTCCATCCAGTCTATTAATTGTTGCCG<br>GGAAGCTAGAGTAAGTAGTTCGCCAGTTAATAGTTTGCGCAACGTTGTTGCCATTGCTACAGGCATCGTGG<br>TGTCACGCTCGTCGTTTGGTATGGCTTCATTCAGCTCCGGTTCCCAACGATCAAGGCGAGTTACATGATCC<br>CCCATGTTGTGCAAAAAAGCGGTTAGCTCCTTCGGTCCTCCGATCGTTGTCAGAAGTAAGTTGGCCGCAGT<br>GTTATCACTCATGGTTATGGCAGCACTGCATAATTCTCTTACTGTCATGCCATCCGTAAGATGCTTTTCTG<br>TGACTGGTGAGTACTCAACCAAGTCATTCTGAGAATAGTGTATGCGGCGACCGAGTTGCTCTTGCCCGGCG<br>TCAATACGGGATAATACCGCGCCACATAGCAGAACTTTAAAAGTGCTCATCATTGGAAAACGTTCTTCGGG<br>GCGAAAACTCTCAAGGATCTTACCGCTGTTGAGATCCAGTTCGATGTAACCCACTCGTGCACCCAACTGAT<br>CTTCAGCATCTTTTACTTTCACCAGCGTTTCTGGGTGAGCAAAAACAGGAAGGCAAAATGCCGCAAAAAAG<br>GGAATAAGGGCGACACGGAAATGTTGAATACTCATACTCTTCCTTTTTCAATATTATTGAAGCATTTATCA<br>GGGTTATTGTCTCATGAGCGGATACATATTTGAATGTATTTAGAAAAATAAACAAATAGGGGTTCCGCGCA<br>CATTTCCCCGAAAAGTGCCACCTGACGTCGACGGATCGGGAGATCTCCCGATCCCCTATGGTCGACTCTCA<br>GTACAATCTGCTCTGATGCCGCATAGTTAAGCCAGTATCTGCTCCCTGCTTGTGTGTTGGAGGTCGCTGAG<br>TAGTGCGCGAGCAAAATTTAAGCTACAACAAGGCAAGGCTTGACCGACAATTGCATGAAGAATCTGCTTAG<br>G |
| pcDSec2-nLuc | GTTAGGCGTTTTGCGCTGCTTCGCGATGTACGGGCCAGATATACGCGTTGACATTGATTATTGACTAGTTA<br>TTAATAGTAATCAATTACGGGGTCATTAGTTCATAGCCCATATATGGAGTTCCGCGTTACATAACTTACGG<br>TAAATGGCCCGCCTGGCTGACCGCCCAACGACCCCCGCCCATTGACGTCAATAATGACGTATGTTCCCATA<br>GTAACGCCAATAGGGACTTTCCATTGACGTCAATGGGTGGAGTATTTACGGTAAACTGCCCACTTGGCAGT<br>ACATCAAGTGTATCATATGCCAAGTACGCCCCCTATTGACGTCAATGACGGTAAATGGCCCGCCTGGCATT<br>ATGCCCAGTACATGACCTTATGGGACTTTCCTACTTGGCAGTACATCTACGTATTAGTCATCGCTATTACC<br>ATGGTGATGCGGTTTTGGCAGTACATCAATGGGCGTGGATAGCGGTTTGACTCACGGGGATTTCCAAGTCT<br>CCACCCCATTGACGTCAATGGGAGTTTGTTTTGGCACCAAAATCAACGGGACTTTCCAAAATGTCGTAACA<br>ACTCCGCCCCATTGACGCAAATGGGCGGTAGGCGTGTACGGTGGGAGGTCTATATAAGCAGAGCTCGTTTA |

GTGAACCGTCAGATCGCCTGGAGACGCCATCCACGCTGTTTTGACCTCCATAGAAGACACCGGGACCGATC
CAGCCTCCGGACTCTAGAGCGGCCGCCACCATGCATCACCACCATCACCATGAAGACGCTAAAAATATTAA
AAAAGGTCCCGCCCCTTTCTACCCTTTGGAAGATGGTACTGCCGGAGAGCAGCTGCATAAAGCAATGAAGA
GATATGCATTGGTTCCCGGTACAATAGCTTTTACGGACGCACATATCGAGGTTGACATTACATATGCCGAG
TACTTCGAGATGTCTGTACGACTGGCAGAAGCGATGAAACGATATGGCCTTAACACCAACCATAGAATAGT
CGTATGCAGTGAGAACTCTCTTCAGTTTTTCATGCCTGTGCTTGGTGCCTTGTTTATTGGGGTTGCTGTGG
CTCCGGCTAATGATATTTATAACGAACGGGAACTTCTGAACAGTATGGGCATATCTCAACCAACAGTCGTG
TTTGTCAGTAAAAAGGGTCTGCAAAAAATCTTGAACGTACAAAAAAAACTGCCGATTATACAGAAAATAAT
CATTATGGACTCAAAGACTGACTATCAAGGCTTTCAATCAATGTACACCTTCGTAACAAGCCACCTTCCCC
CCGGCTTTAATGAATATGACTTTGTACCAGAATCCTTCGACAGGGATAAAACGATTGCATTGATCATGAAC
AGCTCTGGGTCAACTGGGTTGCCAAAAGGGGTCGCGCTCCCCCATAGGACCGCTTGTGTACGATTTAGTCA
TGCGAGAGATCCTATATTCGGAAATCAAATTATTCCTGACACTGCGATTCTGAGCGTCGTACCCTTTCACC
ATGGATTCGGAATGTTTACAACCTTGGGGTATTTGATTTGTGGATTTCGGGTGGTTCTGATGTATAGATTC
GAAGAGGAGTTGTTTCTGAGATCACTCCAAGATTACAAGATACAGTCCGCCCTCCTTGTTCCTACGTTGTT
CAGTTTCTTCGCAAAATCTACACTTATCGACAAATATGACTTGTCTAACTTGCACGAGATTGCCAGCGGAG
GGGCGCCTCTTTCTAAAGAGGTAGGGGAAGCGGTAGCTAAGAGGTTCCATTTGCCCGGTATCAGGCAGGGC
TACGGTTTGACCGAGACTACCTCTGCAATCCTCATAACACCAGAAGGCGATGATAAGCCGGGCGCGGTGGG
CAAGGTGGTTCCGTTTTTCGAGGCTAAAGTGGTCGATCTCGACACTGGCAAAACTCTCGGCGTTAATCAAA
GAGGGGAACTCTGTGTGCGGGGTCCAATGATCATGAGTGGTTACGTCAACAATCCCGAGGCCACAAACGCA
CTCATAGACAAAGATGGGTGGTTGCATTCCGGGGACATCGCGTATTGGGACGAGGATGAACACTTCTTTAT
CGTAGACCGACTCAAGTCTCTTATAAAGTACAAGGGTTACCAAGTGGCCCCAGCAGAGTTGGAGAGCATTT
TGCTCCAGCATCCAAATATTTTTGATGCCGGTGTGGCAGGGTTGCCCGACGACGATGCGGGTGAACTTCCC
GCAGCCGTTGTCGTCCTCGAGCATGGGAAAGGCTCTGGAGGGGGATCTGGAGGGGGATCCTAATAAGGTAC
CAAGCACCACCATCACCATCACTAATGAGATATCTCGACAATCAACCTCTGGATTACAAAATTTGTGAAAG
ATTGACTGGTATTCTTAACTATGTTGCTCCTTTTACGCTATGTGGATACGCTGCTTTAATGCCTTTGTATC
ATGCTATTGCTTCCCGTATGGCTTTCATTTTCTCCTCCTTGTATAAATCCTGGTTGCTGTCTCTTTATGAG
GAGTTGTGGCCCGTTGTCAGGCAACGTGGCGTGGTGTGCACTGTGTTTGCTGACGCAACCCCCACTGGTTG
GGGCATTGCCACCACCTGTCAGCTCCTTTCCGGGACTTTCGCTTTCCCCCTCCCTATTGCCACGGCGGAAC
TCATCGCCGCCTGCCTTGCCCGCTGCTGGACAGGGGCTCGGCTGTTGGGCACTGACAATTCCGTGGTGTTG
TCGGGGAAGCTGACGTCCTTTCCATGGCTGCTCGCCTGTGTTGCCACCTGGATTCTGCGCGGGACGTCCTT
CTGCTACGTCCCTTCGGCCCTCAATCCAGCGGACCTTCCTTCCCGCGGCCTGCTGCCGGCTCTGCGGCCTC
TTCCGCGTCTTCGCCTTCGCCCTCAGACGAGTCGGATCTCCCTTTGGGCCGCCTCCCCGCCTGGAAACGGG
GGAGGCTAACTGAAACACGGAAGGAGACAATACCGGAAGGAACCCGCGCTATGACGGCAATAAAAAGACAG
AATAAAACGCACGGGTGTTGGGTCGTTTGTTCATAAACGCGGGGTTCGGTCCCAGGGCTGGCACTCTGTCG
ATACCCCACCGAGACCCCATTGGGGCCAATACGCCCGCGTTTCTTCCTTTTCCCCACCCCACCCCCCAAGT
TCGGGTGAAGGCCCAGGGCTCGCAGCCAACGTCGGGGCGGCAGGCCCTGCCATAGCAGATCTGCGCAGCTG
GGGCTCTAGGGGGTATCCCCTATACCGTCGACCTCTAGCTAGAGCTTGGCGTAATCATGGTCATAGCTGTT
TCCTGTGTGAAATTGTTATCCGCTCACAATTCCACACAACATACGAGCCGGAAGCATAAAGTGTAAAGCCT
GGGGTGCCTAATGAGTGAGCTAACTCACATTAATTGCGTTGCGCTCACTGCCCGCTTTCCAGTCGGGAAAC
CTGTCGTGCCAGCTGCATTAATGAATCGGCCAACGCGCGGGGAGAGGCGGTTTGCGTATTGGGCGCTCTTC
CGCTTCCTCGCTCACTGACTCGCTGCGCTCGGTCGTTCGGCTGCGGCGAGCGGTATCAGCTCACTCAAAGG
CGGTAATACGGTTATCCACAGAATCAGGGGATAACGCAGGAAAGAACATGTGAGCAAAAGGCCAGCAAAAG
GCCAGGAACCGTAAAAAGGCCGCGTTGCTGGCGTTTTTCCATAGGCTCCGCCCCCCTGACGAGCATCACAA
AAATCGACGCTCAAGTCAGAGGTGGCGAAACCCGACAGGACTATAAAGATACCAGGCGTTTCCCCCTGGAA
GCTCCCTCGTGCGCTCTCCTGTTCCGACCCTGCCGCTTACCGGATACCTGTCCGCCTTTCTCCCTTCGGGA
AGCGTGGCGCTTTCTCATAGCTCACGCTGTAGGTATCTCAGTTCGGTGTAGGTCGTTCGCTCCAAGCTGGG
CTGTGTGCACGAACCCCCCGTTCAGCCCGACCGCTGCGCCTTATCCGGTAACTATCGTCTTGAGTCCAACC
CGGTAAGACACGACTTATCGCCACTGGCAGCAGCCACTGGTAACAGGATTAGCAGAGCGAGGTATGTAGGC
GGTGCTACAGAGTTCTTGAAGTGGTGGCCTAACTACGGCTACACTAGAAGAACAGTATTTGGTATCTGCGC
TCTGCTGAAGCCAGTTACCTTCGGAAAAAGAGTTGGTAGCTCTTGATCCGGCAAACAAACCACCGCTGGTA
GCGGTGGTTTTTTTGTTTGCAAGCAGCAGATTACGCGCAGAAAAAAAGGATCTCAAGAAGATCCTTTGATC
TTTTCTACGGGGTCTGACGCTCAGTGGAACGAAAACTCACGTTAAGGGATTTTGGTCATGAGATTATCAAA
AAGGATCTTCACCTAGATCCTTTTAAATTAAAAATGAAGTTTTAAATCAATCTAAAGTATATATGAGTAAA
CTTGGTCTGACAGTTACCAATGCTTAATCAGTGAGGCACCTATCTCAGCGATCTGTCTATTTCGTTCATCC
ATAGTTGCCTGACTCCCCGTCGTGTAGATAACTACGATACGGGAGGGCTTACCATCTGGCCCCAGTGCTGC
AATGATACCGCGAGACCCACGCTCACCGGCTCCAGATTTATCAGCAATAAACCAGCCAGCCGGAAGGGCCG
AGCGCAGAAGTGGTCCTGCAACTTTATCCGCCTCCATCCAGTCTATTAATTGTTGCCGGGAAGCTAGAGTA
AGTAGTTCGCCAGTTAATAGTTTGCGCAACGTTGTTGCCATTGCTACAGGCATCGTGGTGTCACGCTCGTC
GTTTGGTATGGCTTCATTCAGCTCCGGTTCCCAACGATCAAGGCGAGTTACATGATCCCCCATGTTGTGCA
AAAAAGCGGTTAGCTCCTTCGGTCCTCCGATCGTTGTCAGAAGTAAGTTGGCCGCAGTGTTATCACTCATG
GTTATGGCAGCACTGCATAATTCTCTTACTGTCATGCCATCCGTAAGATGCTTTTCTGTGACTGGTGAGTA
CTCAACCAAGTCATTCTGAGAATAGTGTATGCGGCGACCGAGTTGCTCTTGCCCGGCGTCAATACGGGATA
ATACCGCGCCACATAGCAGAACTTTAAAAGTGCTCATCATTGGAAAACGTTCTTCGGGGCGAAAACTCTCA
AGGATCTTACCGCTGTTGAGATCCAGTTCGATGTAACCCACTCGTGCACCCAACTGATCTTCAGCATCTTT
TACTTTCACCAGCGTTTCTGGGTGAGCAAAAACAGGAAGGCAAAATGCCGCAAAAAAGGGAATAAGGGCGA
CACGGAAATGTTGAATACTCATACTCTTCCTTTTTCAATATTATTGAAGCATTTATCAGGGTTATTGTCTC
ATGAGCGGATACATATTTGAATGTATTTAGAAAAATAAACAAATAGGGGTTCCGCGCACATTTCCCCGAAA
AGTGCCACCTGACGTCGACGGATCGGGAGATCTCCCGATCCCCTATGGTCGACTCTCAGTACAATCTGCTC
TGATGCCGCATAGTTAAGCCAGTATCTGCTCCCTGCTTGTGTGTTGGAGGTCGCTGAGTAGTGCGCGAGCA
AAATTTAAGCTACAACAAGGCAAGGCTTGACCGACAATTGCATGAAGAATCTGCTTAGG

### Supplementary Figures

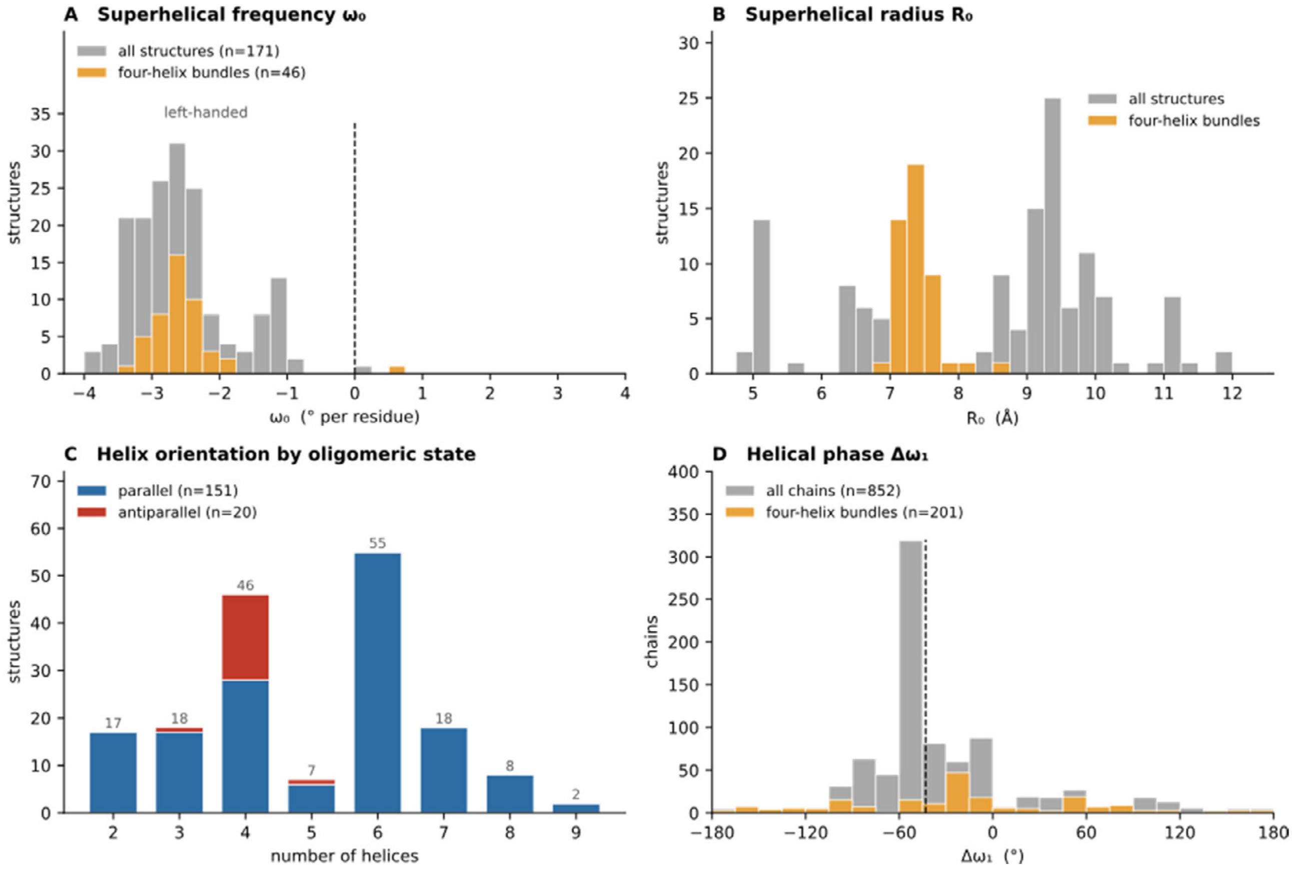


**Figure S1. Distribution of Crick parameters in the reference set of coiled-coil structures.** Crick parameters were determined by fitting 171 coiled-coil structures using CCCP. The reference set comprised 123 PDB entries, some contributing multiple biological assemblies or residue ranges, and four additional de novo designed heterodimers. Gray indicates the complete reference set, and orange highlights the subset of 46 four-helix coiled coils. (A) Superhelical angular frequency ($\omega_0$), with negative and positive values indicating left-handed and right-handed supercoiling, respectively. (B) Superhelical radius ($R_0$), which generally increases with oligomeric state. (C) Distribution of parallel and antiparallel helix orientations across oligomeric states. (D) Rotational phase of the individual helices ($\Delta\omega_1$), calculated for each of the 852 constituent chains. The dashed line indicates the circular mean of −43°, corresponding to the canonical a position of the coiled-coil heptad repeat.

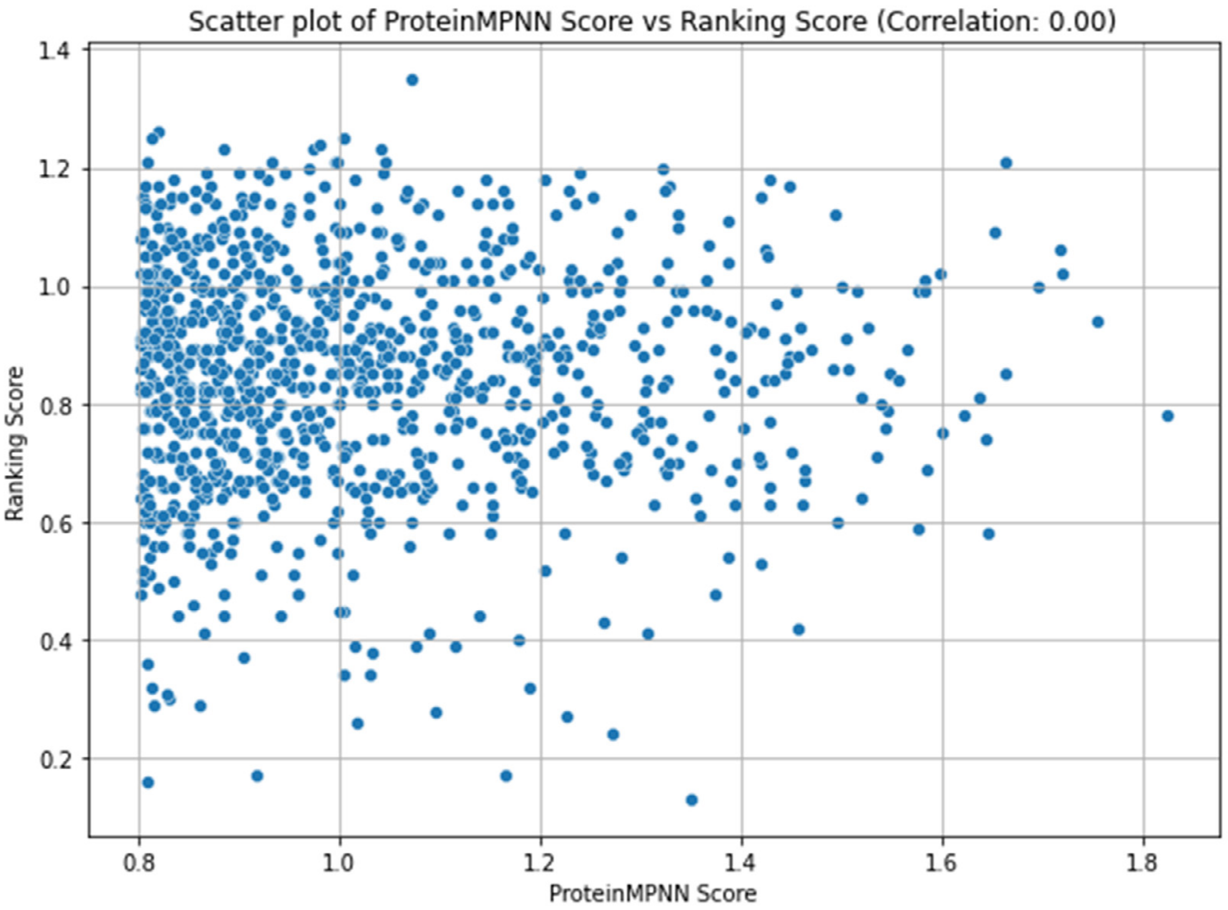


**Figure S2.** Scatter plot of ProteinMPNN sequence scores versus AlphaFold3 (AF3) predicted local distance difference test (pLDDT) scores for sequences generated using design strategy 1 (repeat-constrained homotetramer design).

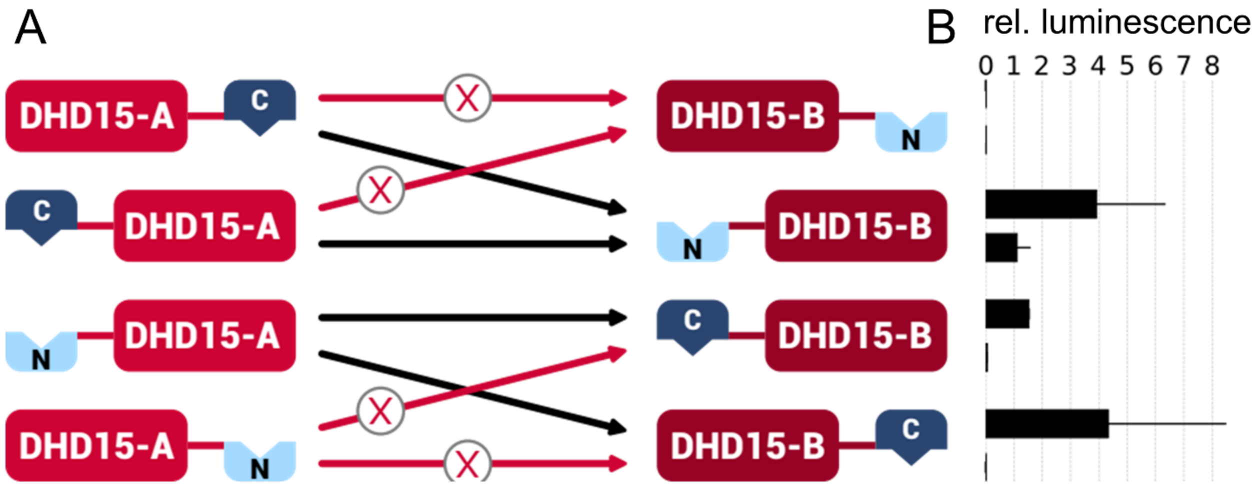


**Figure S3. Optimization of split-firefly luciferase fusion sites using the designed coiled coil DHD15.** (A) Schematic representation of the tested reporter configurations, in which the N-terminal (nLuc) and C-terminal (cLuc) fragments of firefly luciferase were fused to different termini of the two helical hairpin chains of DHD15. (B) Relative luminescence ratios measured in HEK293T cells expressing the indicated constructs. The highest complementation activity, with a luminescence ratio of approximately 4, was obtained when nLuc was fused to the N-terminus of one two-helix chain and cLuc to the C-terminus of the complementary chain. A configuration with both luciferase fragments fused N-terminally also exhibited substantial complementation activity, with a firefly/renilla luminescence ratio of ca. 1. The latter value served as an empirical reference for evaluating assembly-dependent reporter activity and guiding the selection of coiled-coil designs for further characterization. Data are presented as mean ± SD of three independent biological replicates.

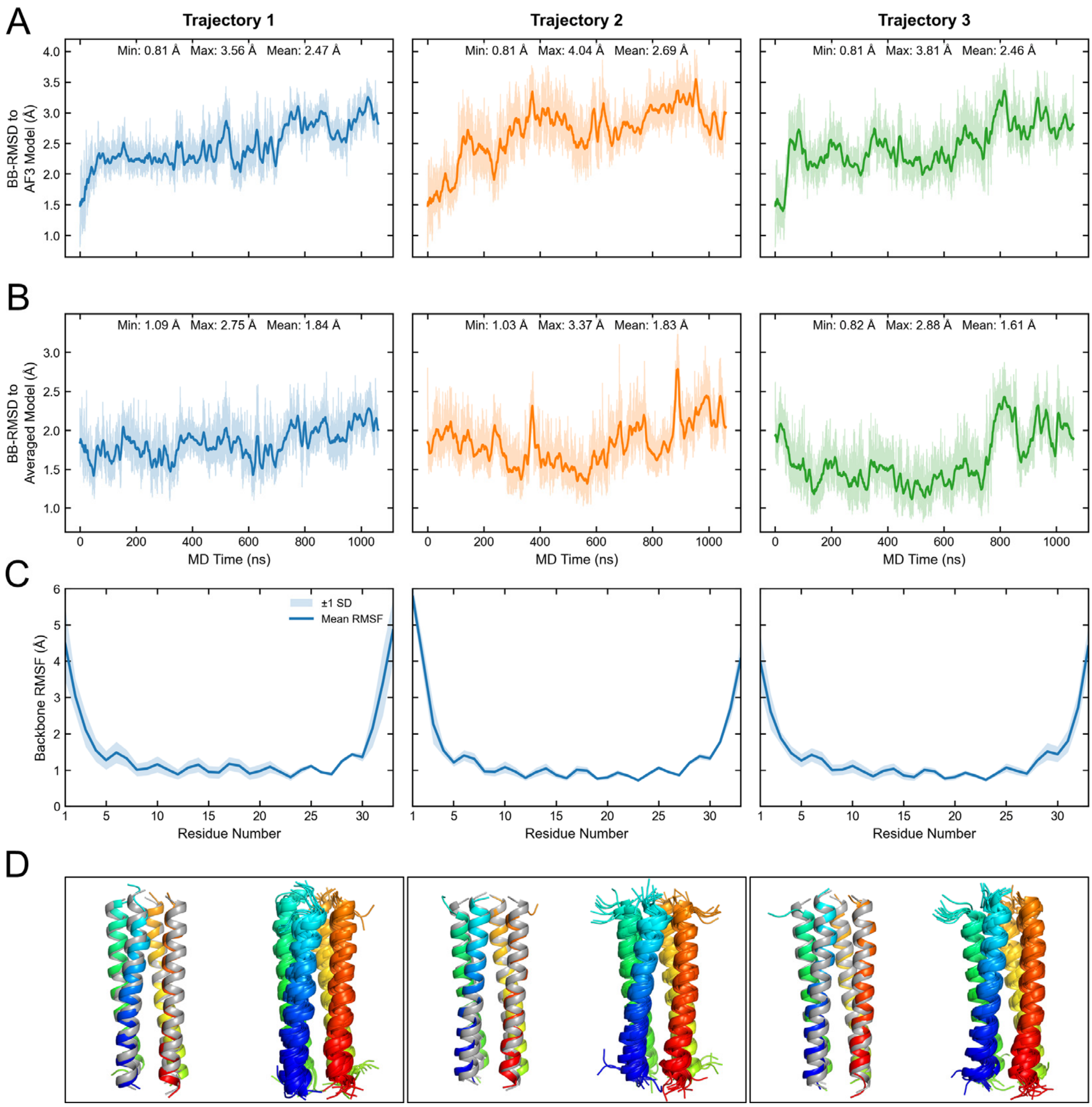


**Figure S4. Structural stability of Homomer-1 during MD simulations at 30 °C.** (A, B) Time evolution of the backbone RMSD relative to (A) the AlphaFold3 input model and (B) the corresponding trajectory-averaged structure for three independent 1.06 μs MD simulations. Transparent lines represent the original RMSD values, while solid lines show the 10 ns running average. Minimum, maximum, and mean RMSD values are indicated in each panel. (C) Backbone RMSF averaged over the four peptide chains for each MD replicate. Solid lines indicate the mean RMSF, and shaded areas represent ±1 standard deviation. (D) Structural comparison and conformational sampling during the MD simulations. Left: Superposition of the MD structure with the lowest Cα RMSD to the trajectory-averaged structure (rainbow coloring) and the AlphaFold3 input model (grey). Right: Superposition of ten structural snapshots sampled at 100 ns intervals from the 1.06 μs MD trajectory.

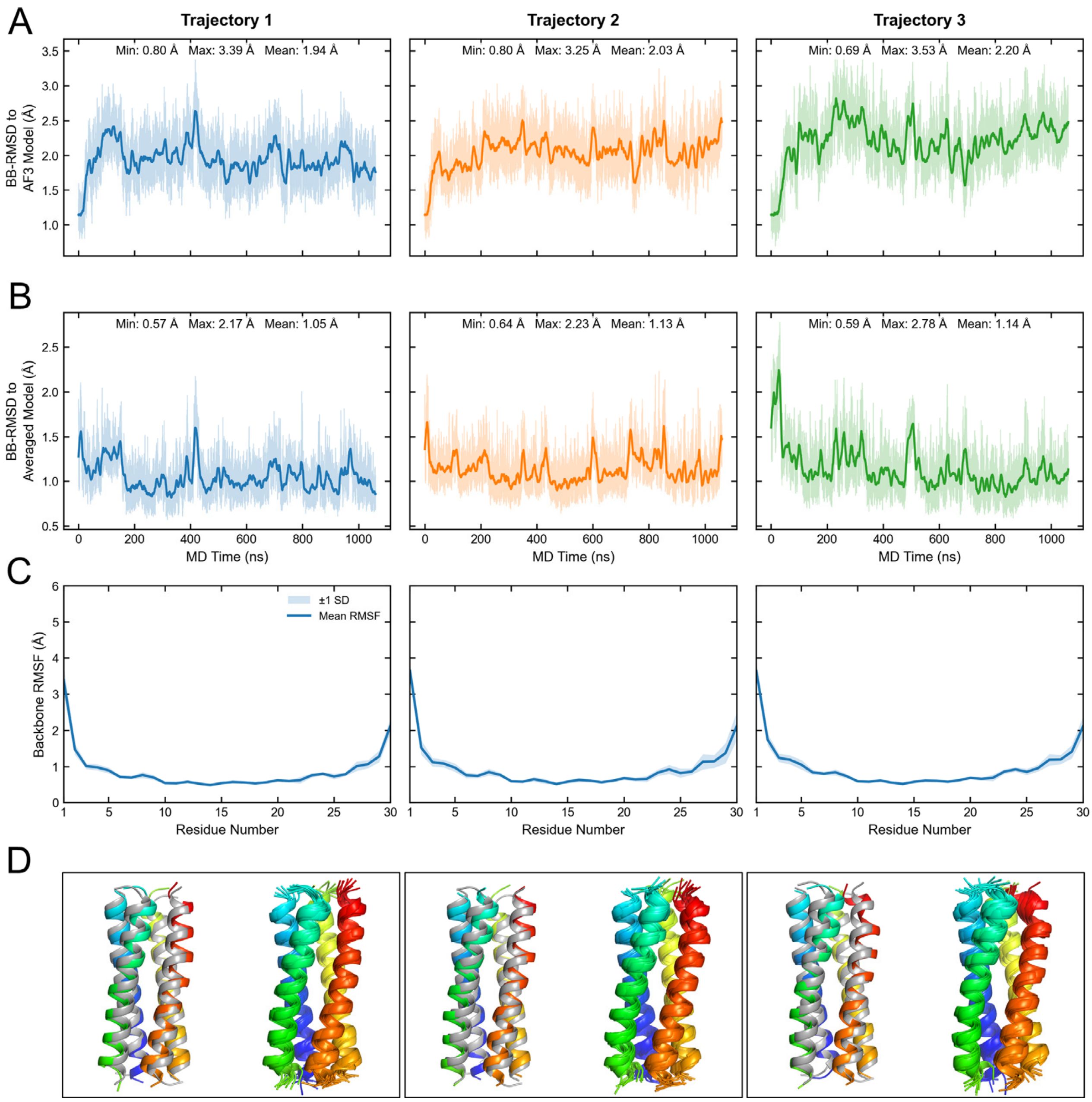


**Figure S5. Structural stability of Homomer-2 during MD simulations at 30 °C.** (A, B) Time evolution of the backbone RMSD relative to (A) the AlphaFold3 input model and (B) the corresponding trajectory-averaged structure for three independent 1.06 μs MD simulations. Transparent lines represent the original RMSD values, while solid lines show the 10 ns running average. Minimum, maximum, and mean RMSD values are indicated in each panel. (C) Backbone RMSF averaged over the four peptide chains for each MD replicate. Solid lines indicate the mean RMSF, and shaded areas represent ±1 standard deviation. (D) Structural comparison and conformational sampling during the MD simulations. Left: Superposition of the MD structure with the lowest Cα RMSD to the trajectory-averaged structure (rainbow coloring) and the AlphaFold3 input model (grey). Right: Superposition of ten structural snapshots sampled at 100 ns intervals from the 1.06 μs MD trajectory.

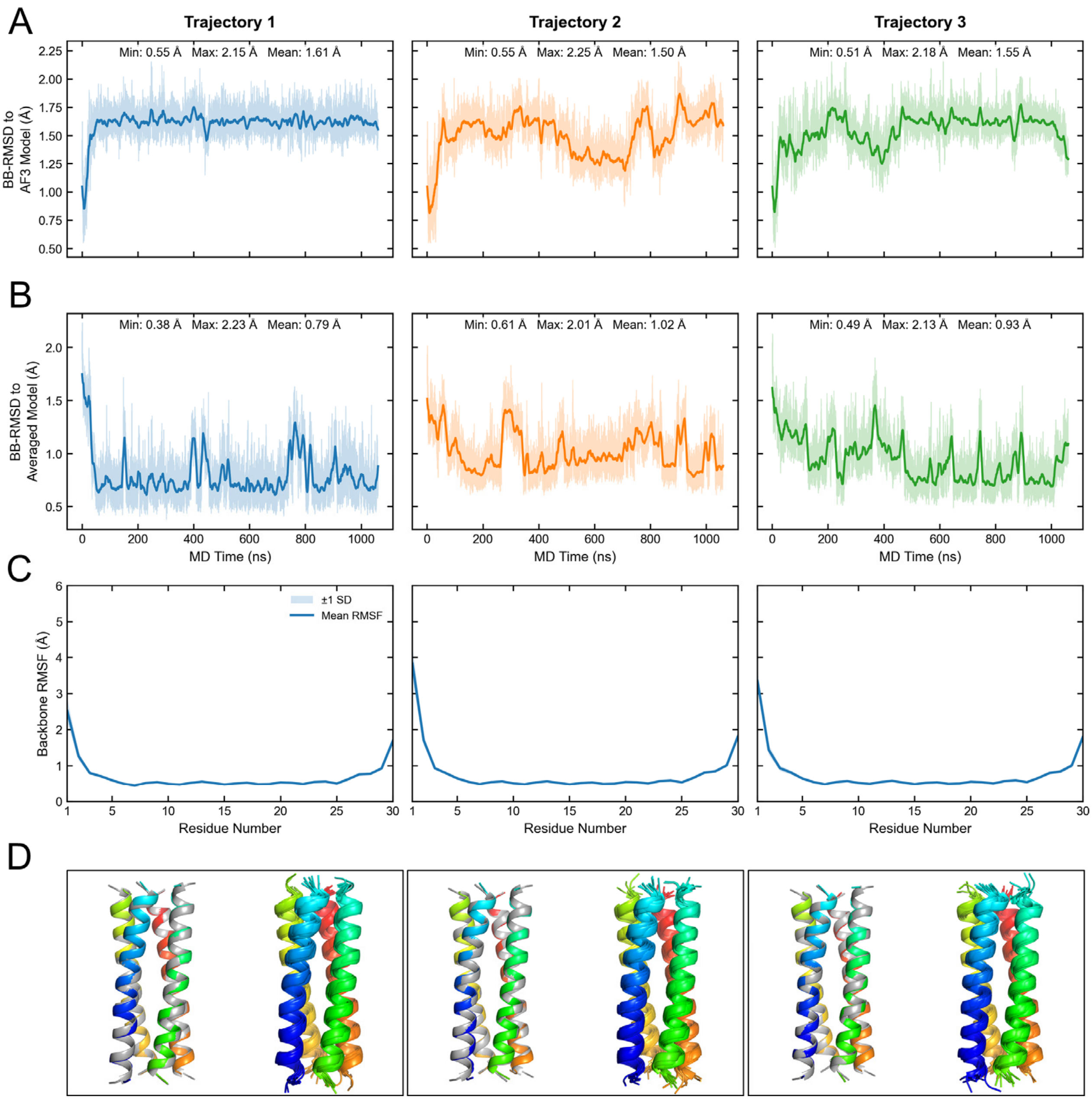


**Figure S6. Structural stability of Homomer-3 during MD simulations at 30 °C.** (A, B) Time evolution of the backbone RMSD relative to (A) the AlphaFold3 input model and (B) the corresponding trajectory-averaged structure for three independent 1.06 μs MD simulations. Transparent lines represent the original RMSD values, while solid lines show the 10 ns running average. Minimum, maximum, and mean RMSD values are indicated in each panel. (C) Backbone RMSF averaged over the four peptide chains for each MD replicate. Solid lines indicate the mean RMSF, and shaded areas represent ±1 standard deviation. (D) Structural comparison and conformational sampling during the MD simulations. Left: Superposition of the MD structure with the lowest Cα RMSD to the trajectory-averaged structure (rainbow coloring) and the AlphaFold3 input model (grey). Right: Superposition of ten structural snapshots sampled at 100 ns intervals from the 1.06 μs MD trajectory.

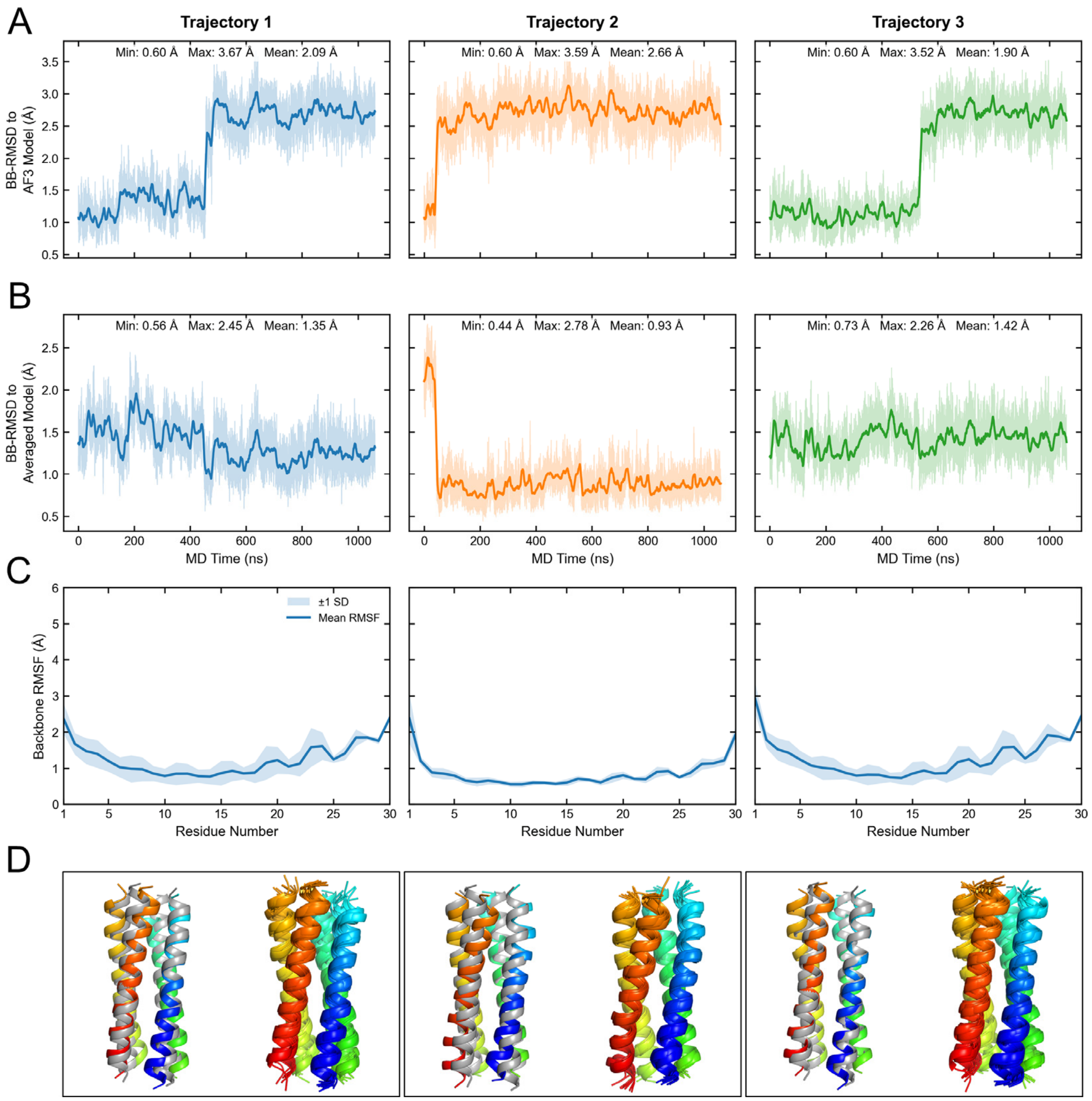


**Figure S7. Structural stability of Homomer-4 during MD simulations at 30 °C.** (A, B) Time evolution of the backbone RMSD relative to (A) the AlphaFold3 input model and (B) the corresponding trajectory-averaged structure for three independent 1.06 μs MD simulations. Transparent lines represent the original RMSD values, while solid lines show the 10 ns running average. Minimum, maximum, and mean RMSD values are indicated in each panel. (C) Backbone RMSF averaged over the four peptide chains for each MD replicate. Solid lines indicate the mean RMSF, and shaded areas represent ±1 standard deviation. (D) Structural comparison and conformational sampling during the MD simulations. Left: Superposition of the MD structure with the lowest Cα RMSD to the trajectory-averaged structure (rainbow coloring) and the AlphaFold3 input model (grey). Right: Superposition of ten structural snapshots sampled at 100 ns intervals from the 1.06 μs MD trajectory.

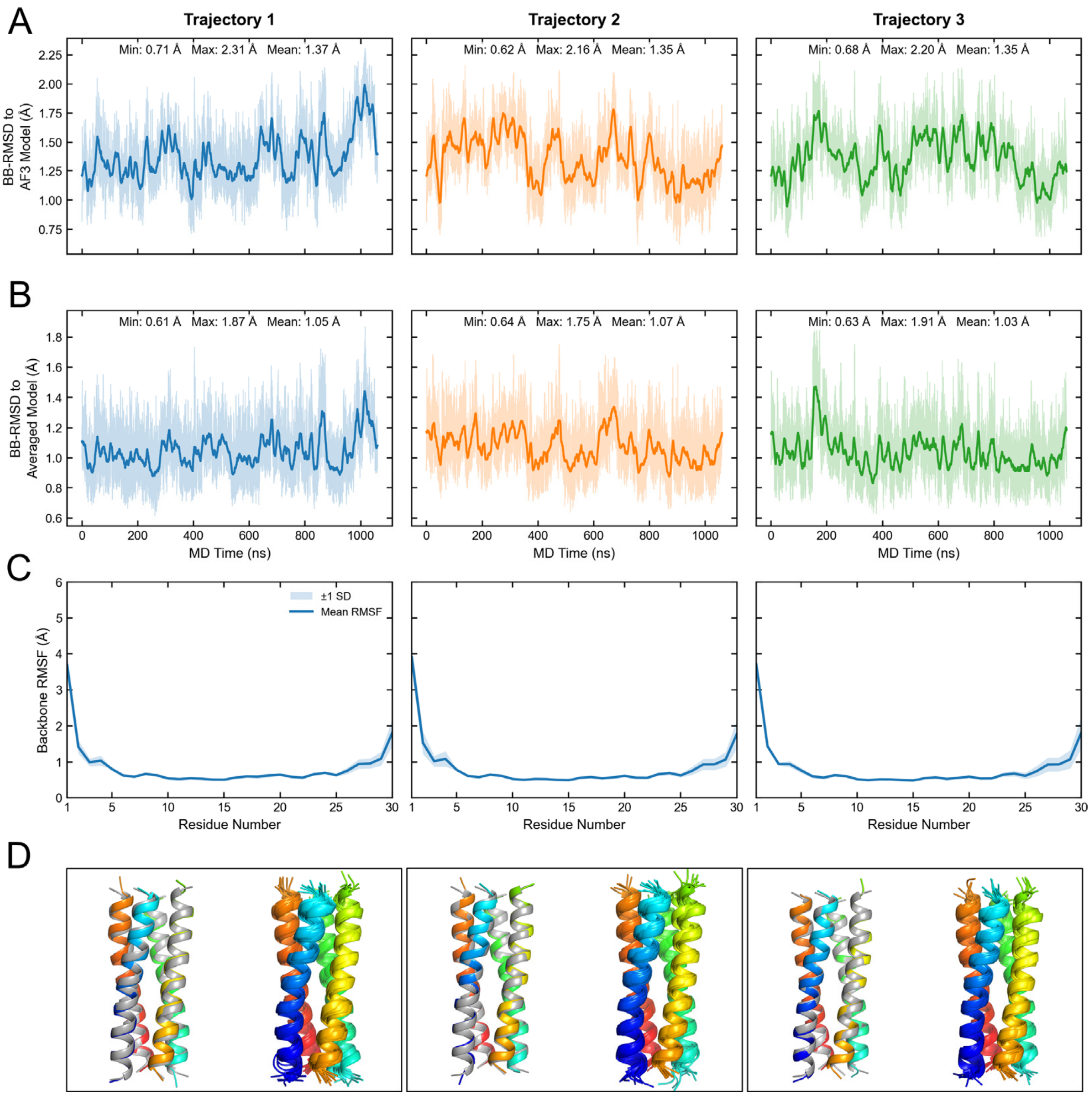


**Figure S8. Structural stability of Heteromer-1 during MD simulations at 30 °C.** (A, B) Time evolution of the backbone RMSD relative to (A) the AlphaFold3 input model and (B) the corresponding trajectory-averaged structure for three independent 1.06 μs MD simulations. Transparent lines represent the original RMSD values, while solid lines show the 10 ns running average. Minimum, maximum, and mean RMSD values are indicated in each panel. (C) Backbone RMSF averaged over the four peptide chains for each MD replicate. Solid lines indicate the mean RMSF, and shaded areas represent ±1 standard deviation. (D) Structural comparison and conformational sampling during the MD simulations. Left: Superposition of the MD structure with the lowest Cα RMSD to the trajectory-averaged structure (rainbow coloring) and the AlphaFold3 input model (grey). Right: Superposition of ten structural snapshots sampled at 100 ns intervals from the 1.06 μs MD trajectory.

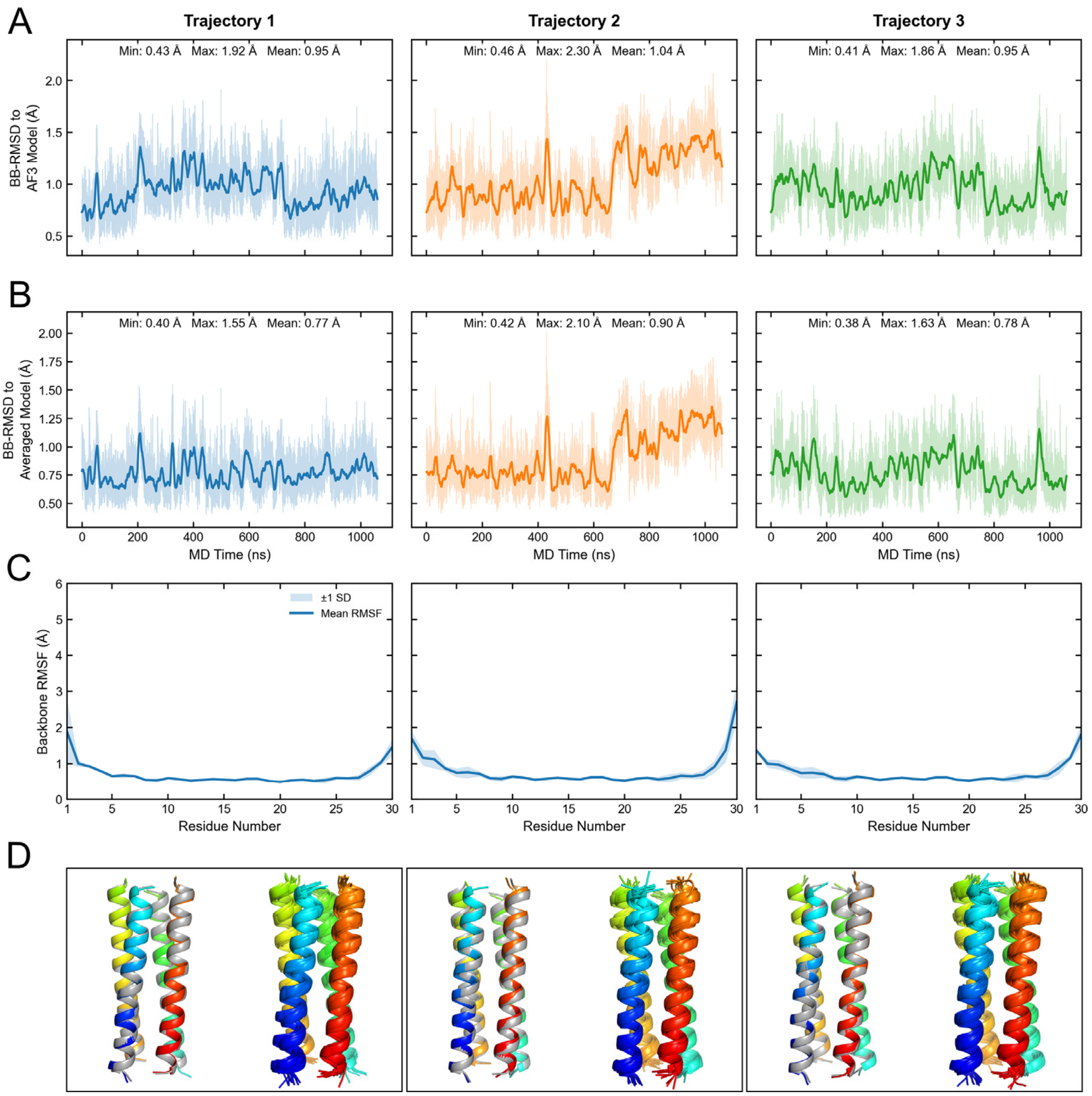


**Figure S9. Structural stability of Heteromer-2 during MD simulations at 30 °C.** (A, B) Time evolution of the backbone RMSD relative to (A) the AlphaFold3 input model and (B) the corresponding trajectory-averaged structure for three independent 1.06 µs MD simulations. Transparent lines represent the original RMSD values, while solid lines show the 10 ns running average. Minimum, maximum, and mean RMSD values are indicated in each panel. (C) Backbone RMSF averaged over the four peptide chains for each MD replicate. Solid lines indicate the mean RMSF, and shaded areas represent ±1 standard deviation. (D) Structural comparison and conformational sampling during the MD simulations. Left: Superposition of the MD structure with the lowest Cα RMSD to the trajectory-averaged structure (rainbow coloring) and the AlphaFold3 input model (grey). Right: Superposition of ten structural snapshots sampled at 100 ns intervals from the 1.06 µs MD trajectory.

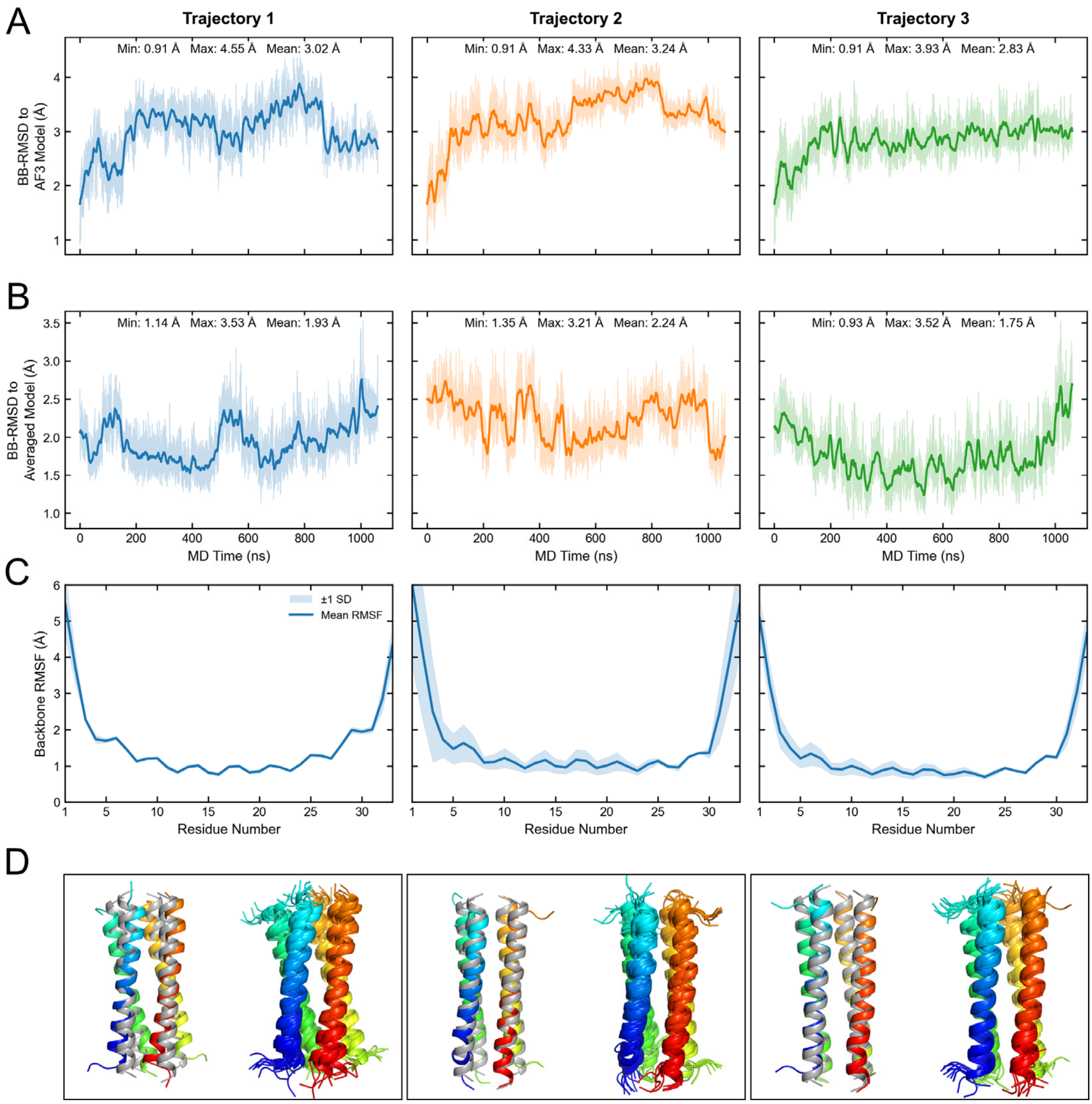


**Figure S10. Structural stability of Homomer-1 during MD simulations at 40 °C.** (A, B) Time evolution of the backbone RMSD relative to (A) the AlphaFold3 input model and (B) the corresponding trajectory-averaged structure for three independent 1.06 μs MD simulations. Transparent lines represent the original RMSD values, while solid lines show the 10 ns running average. Minimum, maximum, and mean RMSD values are indicated in each panel. (C) Backbone RMSF averaged over the four peptide chains for each MD replicate. Solid lines indicate the mean RMSF, and shaded areas represent ±1 standard deviation. (D) Structural comparison and conformational sampling during the MD simulations. Left: Superposition of the MD structure with the lowest Cα RMSD to the trajectory-averaged structure (rainbow coloring) and the AlphaFold3 input model (grey). Right: Superposition of ten structural snapshots sampled at 100 ns intervals from the 1.06 μs MD trajectory.

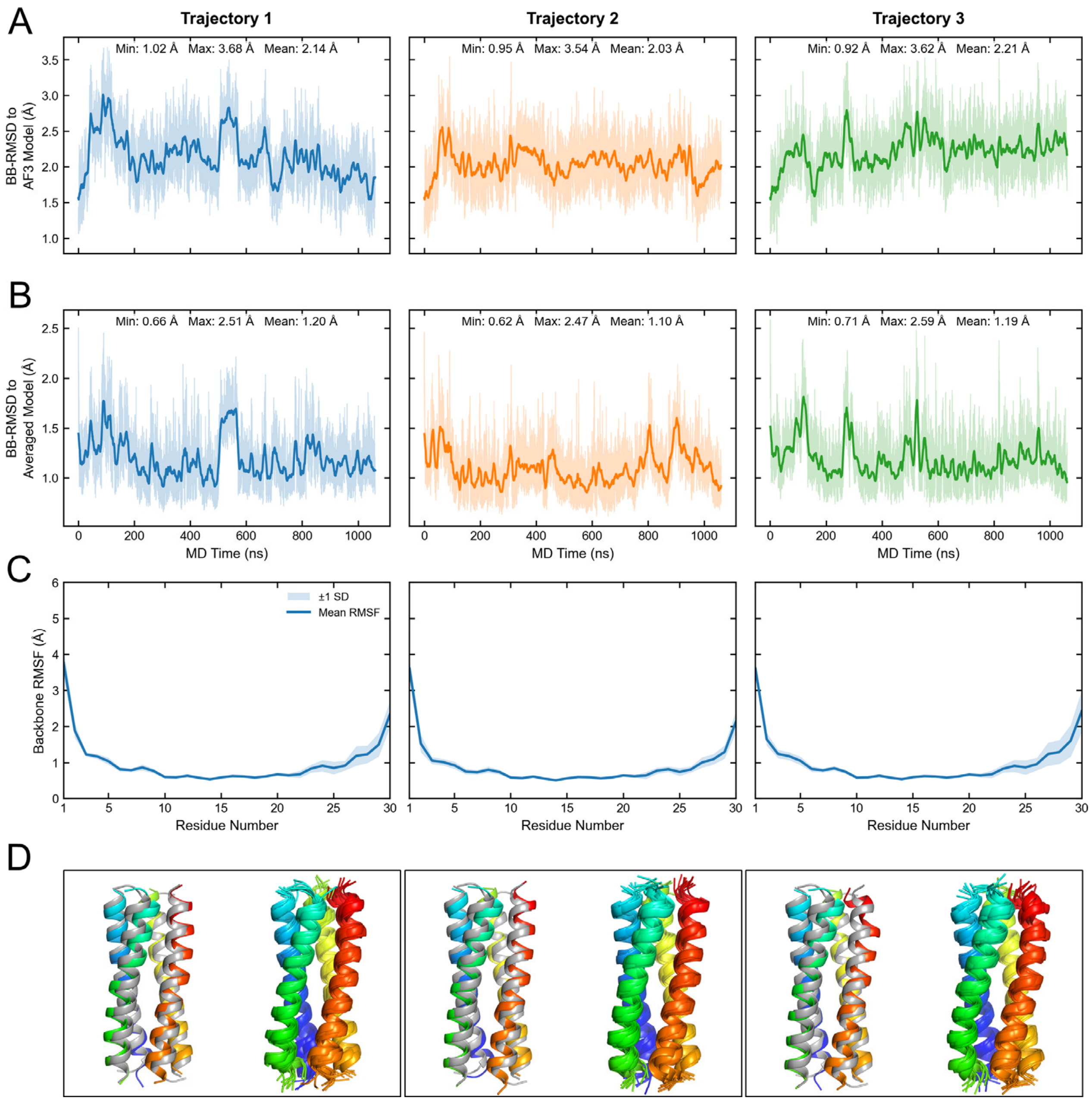


**Figure S11. Structural stability of Homomer-2 during MD simulations at 40 °C.** (A, B) Time evolution of the backbone RMSD relative to (A) the AlphaFold3 input model and (B) the corresponding trajectory-averaged structure for three independent 1.06 μs MD simulations. Transparent lines represent the original RMSD values, while solid lines show the 10 ns running average. Minimum, maximum, and mean RMSD values are indicated in each panel. (C) Backbone RMSF averaged over the four peptide chains for each MD replicate. Solid lines indicate the mean RMSF, and shaded areas represent ±1 standard deviation. (D) Structural comparison and conformational sampling during the MD simulations. Left: Superposition of the MD structure with the lowest Cα RMSD to the trajectory-averaged structure (rainbow coloring) and the AlphaFold3 input model (grey). Right: Superposition of ten structural snapshots sampled at 100 ns intervals from the 1.06 μs MD trajectory.

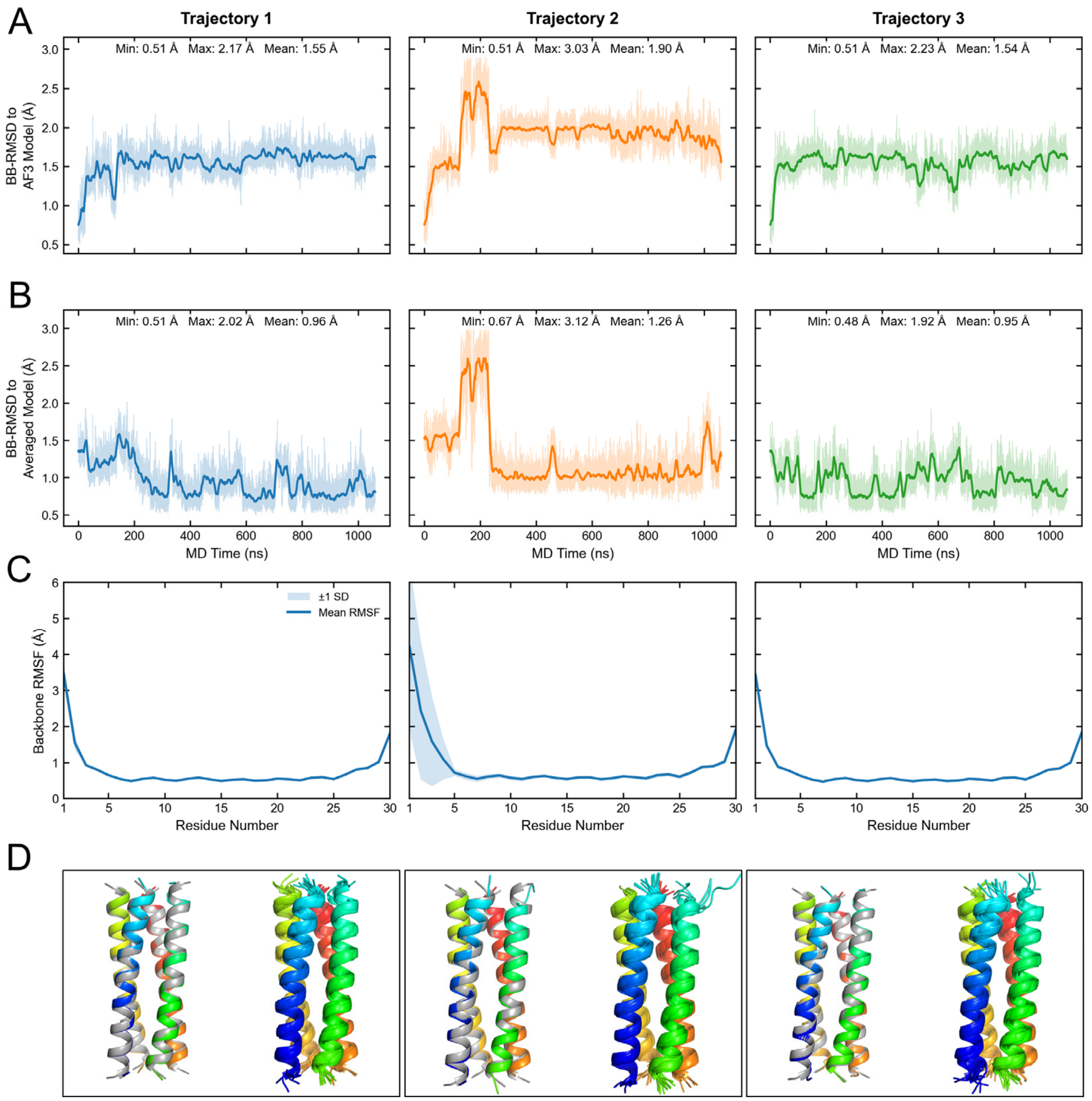


**Figure S12. Structural stability of Homomer-3 during MD simulations at 40 °C.** (A, B) Time evolution of the backbone RMSD relative to (A) the AlphaFold3 input model and (B) the corresponding trajectory-averaged structure for three independent 1.06 μs MD simulations. Transparent lines represent the original RMSD values, while solid lines show the 10 ns running average. Minimum, maximum, and mean RMSD values are indicated in each panel. (C) Backbone RMSF averaged over the four peptide chains for each MD replicate. Solid lines indicate the mean RMSF, and shaded areas represent ±1 standard deviation. (D) Structural comparison and conformational sampling during the MD simulations. Left: Superposition of the MD structure with the lowest Cα RMSD to the trajectory-averaged structure (rainbow coloring) and the AlphaFold3 input model (grey). Right: Superposition of ten structural snapshots sampled at 100 ns intervals from the 1.06 μs MD trajectory.

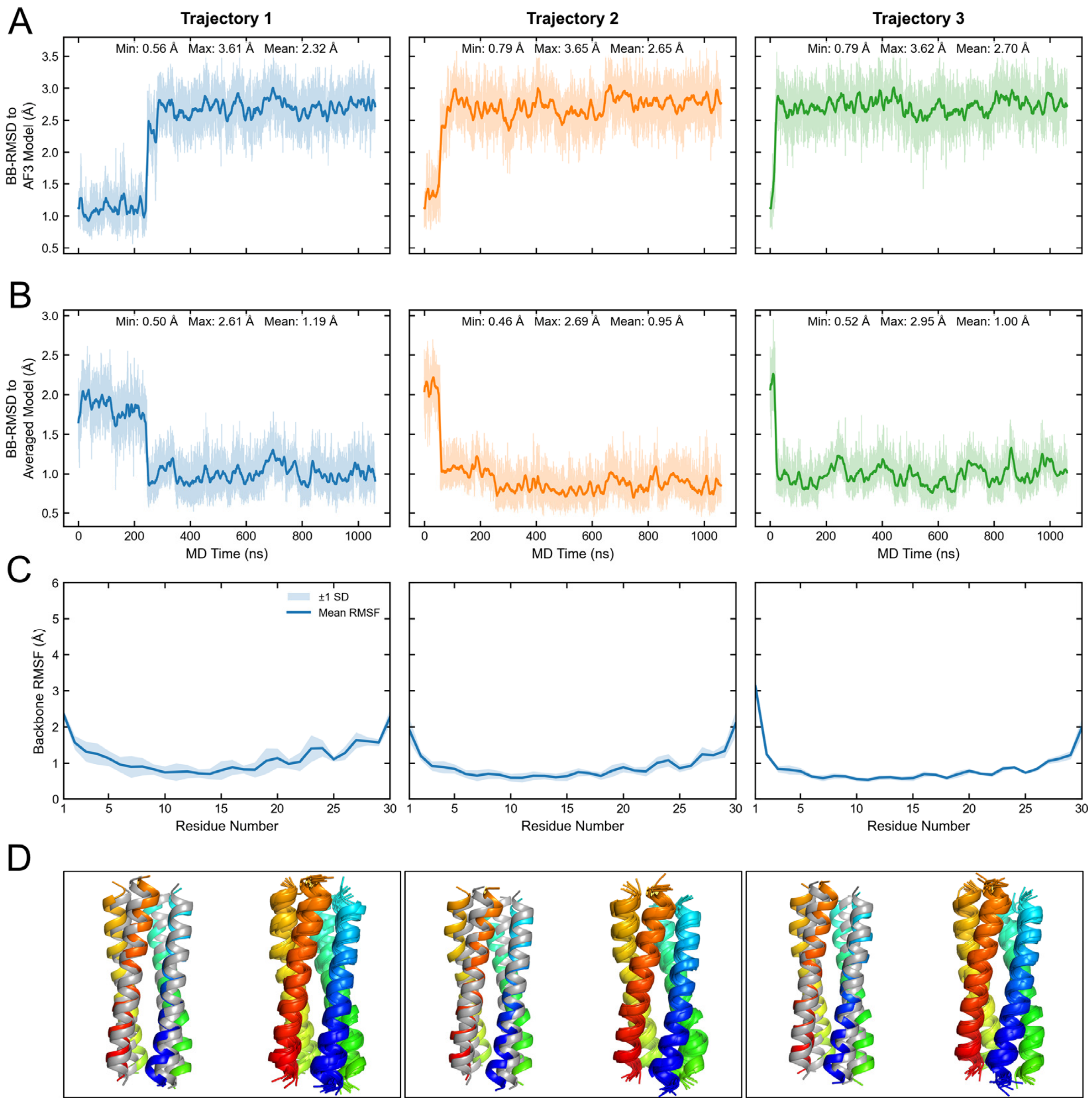


**Figure S13. Structural stability of Homomer-4 during MD simulations at 40 °C.** (A, B) Time evolution of the backbone RMSD relative to (A) the AlphaFold3 input model and (B) the corresponding trajectory-averaged structure for three independent 1.06 μs MD simulations. Transparent lines represent the original RMSD values, while solid lines show the 10 ns running average. Minimum, maximum, and mean RMSD values are indicated in each panel. (C) Backbone RMSF averaged over the four peptide chains for each MD replicate. Solid lines indicate the mean RMSF, and shaded areas represent ±1 standard deviation. (D) Structural comparison and conformational sampling during the MD simulations. Left: Superposition of the MD structure with the lowest Cα RMSD to the trajectory-averaged structure (rainbow coloring) and the AlphaFold3 input model (grey). Right: Superposition of ten structural snapshots sampled at 100 ns intervals from the 1.06 μs MD trajectory.

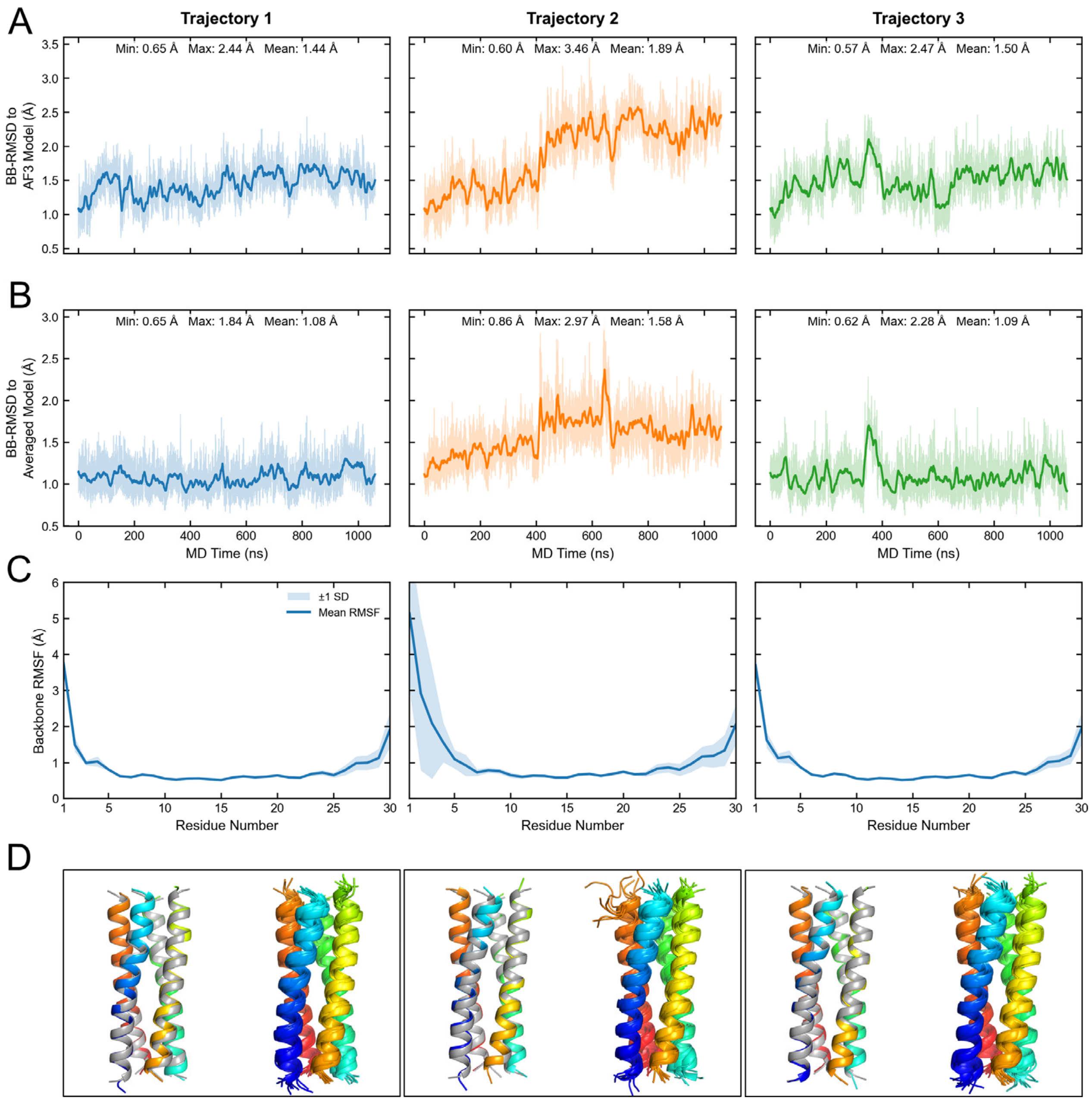


**Figure S14. Structural stability of Heteromer-1 during MD simulations at 40 °C.** (A, B) Time evolution of the backbone RMSD relative to (A) the AlphaFold3 input model and (B) the corresponding trajectory-averaged structure for three independent 1.06 μs MD simulations. Transparent lines represent the original RMSD values, while solid lines show the 10 ns running average. Minimum, maximum, and mean RMSD values are indicated in each panel. (C) Backbone RMSF averaged over the four peptide chains for each MD replicate. Solid lines indicate the mean RMSF, and shaded areas represent ±1 standard deviation. (D) Structural comparison and conformational sampling during the MD simulations. Left: Superposition of the MD structure with the lowest Cα RMSD to the trajectory-averaged structure (rainbow coloring) and the AlphaFold3 input model (grey). Right: Superposition of ten structural snapshots sampled at 100 ns intervals from the 1.06 μs MD trajectory.

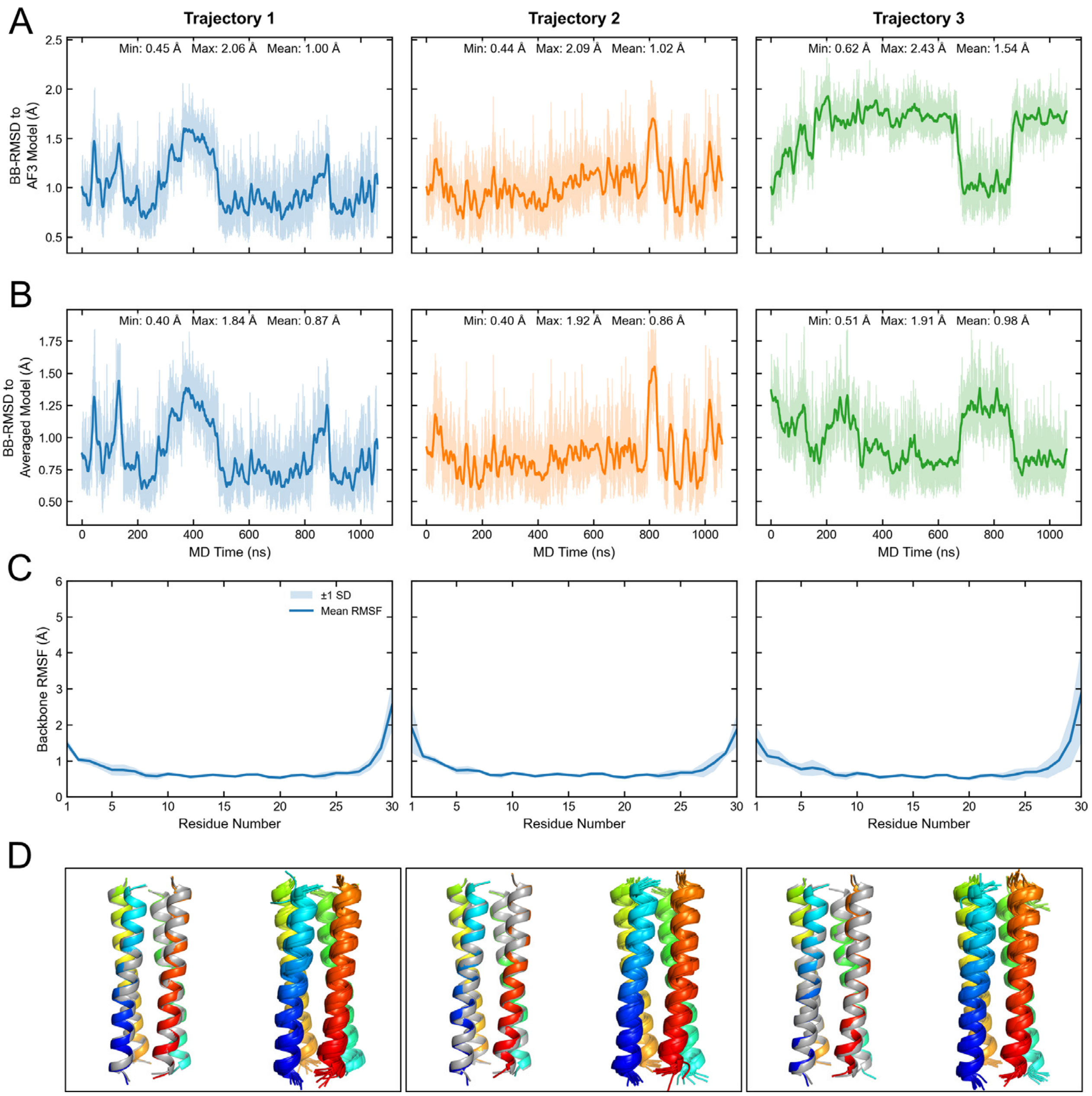


**Figure S15. Structural stability of Heteromer-2 during MD simulations at 40 °C.** (A, B) Time evolution of the backbone RMSD relative to (A) the AlphaFold3 input model and (B) the corresponding trajectory-averaged structure for three independent 1.06 μs MD simulations. Transparent lines represent the original RMSD values, while solid lines show the 10 ns running average. Minimum, maximum, and mean RMSD values are indicated in each panel. (C) Backbone RMSF averaged over the four peptide chains for each MD replicate. Solid lines indicate the mean RMSF, and shaded areas represent ±1 standard deviation. (D) Structural comparison and conformational sampling during the MD simulations. Left: Superposition of the MD structure with the lowest Cα RMSD to the trajectory-averaged structure (rainbow coloring) and the AlphaFold3 input model (grey). Right: Superposition of ten structural snapshots sampled at 100 ns intervals from the 1.06 μs MD trajectory.
